# Firing Without Wiring: How REM-like Sleep Reactivation Rebalances Excitation and Inhibition to Orthogonalize Memories

**DOI:** 10.64898/2026.10.11.757684

**Authors:** Yoshimasa Kubo, Jean Erik Delanois, Maxim Bazhenov

**Affiliations:** Department of Medicine, University of California San Diego, La Jolla, CA, USA

**Keywords:** REM sleep, memory consolidation, memory interference, synaptic plasticity, artificial neural networks

## Abstract

The role of Rapid Eye Movement (REM) sleep in memory consolidation remains poorly understood. Several influential ideas propose that REM sleep serves to eliminate crosstalk between memories, thereby orthogonalizing representations of similar ones; however, mechanistic confirmation remains sparse. Here we used recurrent neural networks trained with Equilibrium Propagation (EP), a biologically plausible training algorithm, to investigate the role of REM-like sleep in memory consolidation. We found that REM-like sleep reduces memory interference by decorrelating task-specific neuronal populations, isolating individual memory traces. During sleep, reactivation was dominated by the population encoding the most recent memory, which had taken over the inputs of older memories during new learning. The excitation/inhibition balance of inputs onto this population shifted toward inhibition, most strongly from older-memory populations, so that older memories again activated their own neurons. Spreading the same total synaptic change evenly across all target populations abolished this recovery. We call this mechanism reactivation-targeted pruning: synaptic weights onto the reactivated population decrease when its neurons fire while their presynaptic partners are silent. The mechanism held under a synapse-level Dale’s law, and excitatory plasticity alone, with inhibitory synapses fixed, was sufficient. These findings suggest mechanistic principles underlying REM sleep’s role in memory consolidation.

## 1 Introduction

Rapid eye movement (REM) sleep has remained mechanistically enigmatic since its discovery seventy years ago [1, 2]. It is phylogenetically conserved, follows a stereotyped developmental trajectory, and occupies roughly a fifth of human sleep [3, 4], yet a causal, cellular-level account of what REM sleep actually does for memory has been difficult to establish. This stands in sharp contrast to non-REM (NREM) sleep, where slow oscillations, spindles, and hippocampal sharp-wave ripples have been directly linked to the offline replay and systems-level consolidation of recently acquired memories [5, 6]. During REM sleep, hippocampal and neocortical activity dynamics closely resemble wakefulness, yet evidence for veridical, content-specific memory replay during REM sleep is comparatively sparse, leaving REM sleep’s contribution to memory consolidation open.

Several influential theories converge on a different function for REM sleep: not further consolidation of individual memories, but removal of crosstalk between memory representations that share overlapping features. Crick and Mitchison’s reverse-learning hypothesis proposed that REM dreaming actively unlearns spurious or parasitic associations in cortical networks [7]. The synaptic homeostasis hypothesis emphasizes slow-wave sleep for global synaptic downscaling, but has also raised the possibility that REM sleep selectively maintains some connections while allowing others to weaken [8, 9]. Other accounts frame REM sleep as an error-correction, or “forgetting,” process that exposes and eliminates inappropriate associative links [6, 10], and a recently proposed Refinement-and-Rescue hypothesis suggests that REM sleep sparsifies memories to their essential components, with pattern separation between overlapping memories emerging as a byproduct of this sparsification [11]. These proposals share one idea: REM sleep removes crosstalk between memories and thereby makes the representations of similar memories more orthogonal, which minimizes interference in distributed associative memories [12].

This proposed function has a plausible neuromodulatory substrate. The distinctive chemistry of REM sleep – reduced noradrenergic and serotonergic tone alongside elevated cholinergic tone – can shift spike-timing-dependent plasticity toward long-term depression [13, 14] and is associated with a decrease in cortical excitation/inhibition balance during REM sleep following learning [15]. Together, these conditions are well positioned to decorrelate memory traces that may have been partially merged during preceding NREM sleep, when replay [16] is thought to integrate new memories with existing ones [5, 6]. What is missing is a testable account, at the level of synapses, of how this decorrelation comes about.

Because complete, longitudinal in vivo access to an evolving synaptic weight matrix is not feasible, computational models offer a tractable route to such a mechanistic account. Here we use recurrent neural networks (RNNs) trained with Equilibrium Propagation (EP) [17, 18], a local, contrastive, biologically plausible learning rule, to ask whether a REM-like reactivation phase, acting through local plasticity alone, can change connectivity so that overlapping memories become separated, and how it does so. The sleep phase was implemented in a spiking version of the model with a local plasticity rule: coincident pre- and postsynaptic spikes strengthened a synapse [19], and postsynaptic spikes without presynaptic input weakened it [20, 21]. We show that sleep dynamics rescue old memories from catastrophic forgetting by reducing between-class correlation and the overlap between task-specific neuronal populations. This separation comes from changes to the synapses onto the population of the most recently trained task, which dominates reactivation during sleep. These synapses shift toward inhibition, through depression of excitatory and potentiation of inhibitory synapses, most strongly those arriving from older-task populations. We call this mechanism *reactivation-targeted pruning* : plasticity during sleep is concentrated on the reactivated population, whose neurons lose the inputs that are silent when they fire. By directly manipulating the sleep-induced weight changes, we show that this targeting of the reactivated population is what restores older memories. The mechanism holds under a synapse-level Dale’s law, and excitatory plasticity alone, with inhibitory synapses held fixed, is sufficient; it also holds in recurrent networks trained with backpropagation through time, in feedforward networks and on natural images. Together, these results give a mechanistic, testable account of how REM sleep could decorrelate memories.

## 2 Results

The paper is organized as follows. We first show that a REM-like sleep phase rescues older memories from interference by new learning in EP-trained recurrent neural networks (MRNN-EP; Fig. 1). We then show that sleep decorrelates class representations, separates the neuronal populations recruited by different tasks, and restores the ability of older-task inputs to drive their own neurons (Fig. 2). Next, we identify how sleep reorganizes synaptic connectivity (Fig. 3), which feature of this reorganization is required for recovery (Fig. 4) and why it emerges from local plasticity (Fig. 5). A minimal spiking network then follows the mechanism from synapses to network dynamics and shows its limit for memories that overlap most with the most recent one (Fig. 6). Finally, we show that the mechanism holds under a synapse-level Dale’s law – including when only excitatory synapses are plastic during sleep – in recurrent and feedforward networks trained with backpropagation, and on natural images (Fig. 7).

**Fig. 1:**
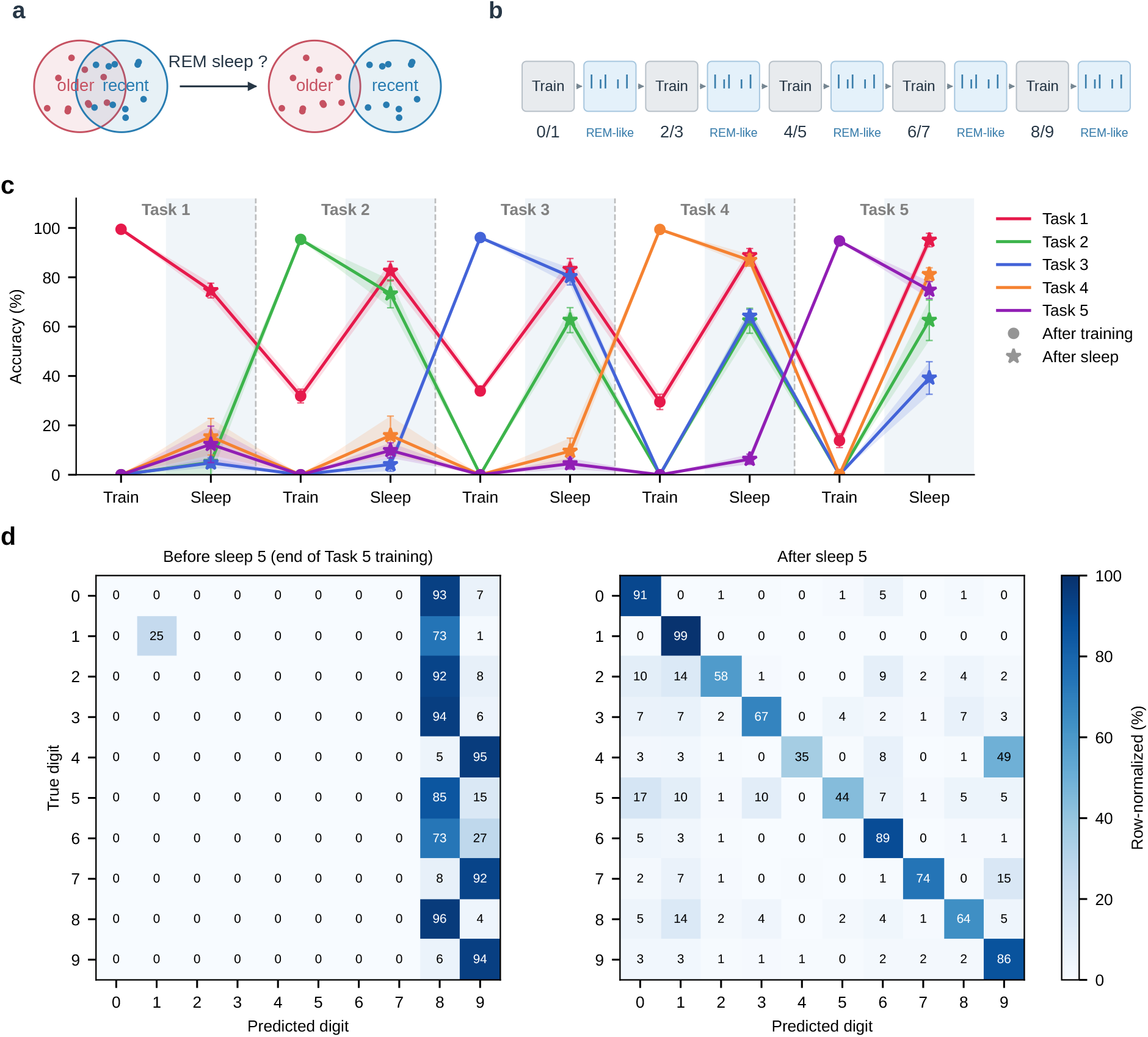
Catastrophic forgetting and sleep recovery in an RNN trained with EP. **(a)** Question addressed: after sequential learning, the neuronal representations of an older and a recently learned memory overlap; can a REM-like sleep phase separate them? **(b)** Protocol. MRNN-EP is trained sequentially on five two-class MNIST tasks (digits 0/1, 2/3, 4/5, 6/7, 8/9); each training block is followed by a REM-like sleep phase (spiking replay driven by Poisson noise with the input statistics of the task just trained). We use T# for the training phase of task # and S# for the sleep phase that follows it. **(c)** Validation accuracy of each of the five tasks at the end of every training (circles) and sleep (stars) phase, mean ± SD over six independent trials (shading and error bars); grey shading marks sleep. After each new task is introduced, prior-task accuracy decreases (catastrophic interference); sleep partially or fully restores it. **(d)** Confusion matrices immediately before (left; end of Task 5 training) and after (right) the final sleep phase, row-normalized (%), averaged over six trials. Before sleep, almost all digits are classified as 8 or 9, the classes of the last trained task.

**Fig. 2:**
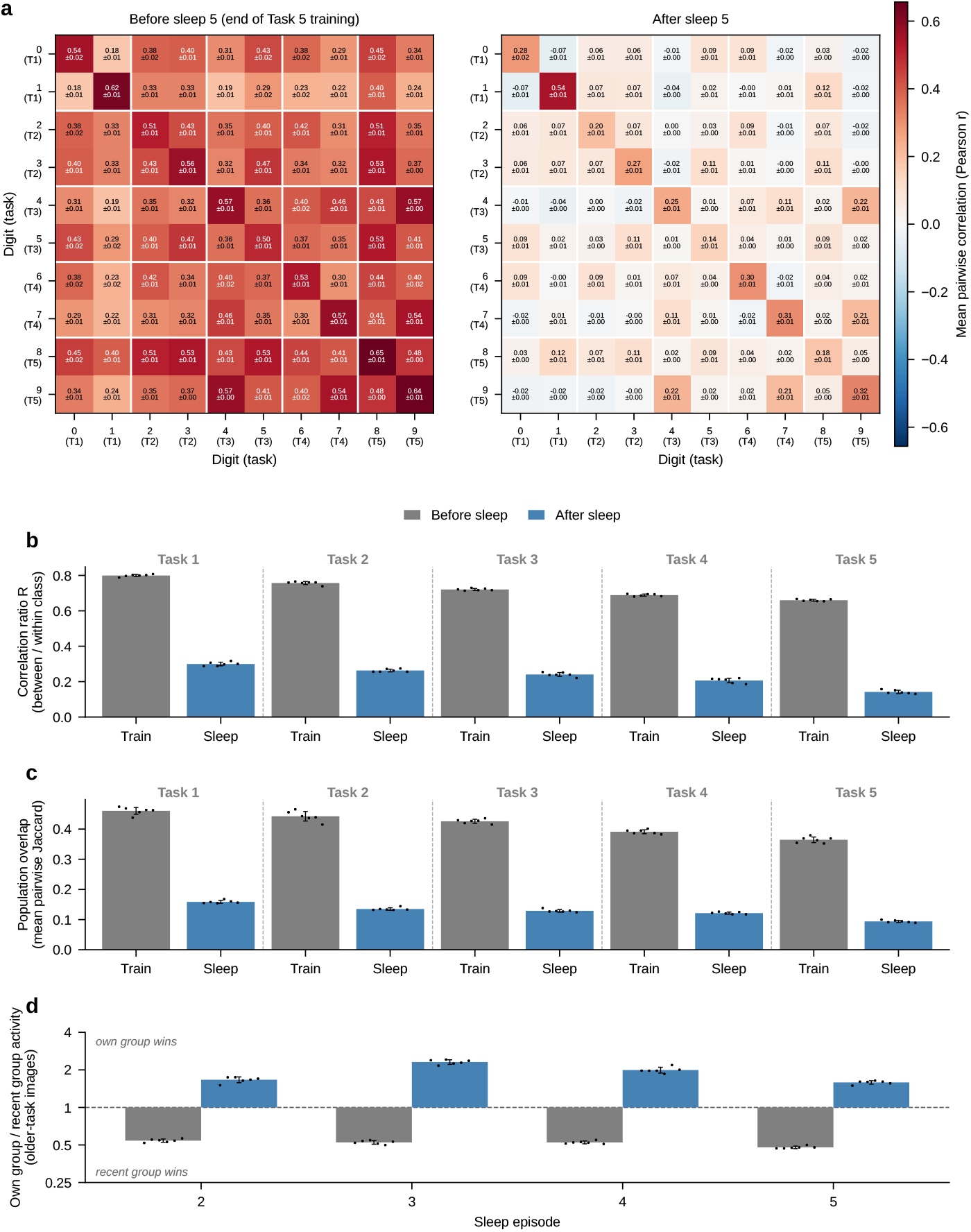
Sleep decorrelates hidden representations, separates task populations and restores the routing of older-task inputs. **(a)** Class-level activation correlation at the end of Task 5 training (left) and after the final sleep (right). Each cell shows the mean ± SD Pearson correlation of hidden activations between classes *i* (row) and *j* (column), averaged across six trials; the task of each digit is given in parentheses. Diagonal entries reflect within-class pairwise sample correlation; off-diagonal entries reflect between-class correlations. After sleep, between-class correlations are strongly suppressed. **(b)** Correlation ratio *R* = mean between-class correlation*/*mean within-class correlation of the hidden layer at the end of each training block (grey) and after the following sleep episode (blue). A lower *R* indicates more class-selective representations. Sleep consistently reduces *R* at every task. **(c)** Population overlap: mean pairwise Jaccard index between the top-10% most active neuron populations of the ten classes, with populations redefined at every phase. Overlap falls sharply after each sleep. **(d)** Intrusion index *Q*: mean activity of an older task’s own hidden population divided by that of the most recent task’s population, measured on the older task’s validation images and averaged over older tasks, before (grey) and after (blue) sleep episodes 2–5 (log scale). *Q <* 1: older-task inputs drive the recent task’s neurons more than their own; *Q >* 1: they drive their own neurons. (b–d) Mean ± SD over six trials; dots, individual trials.

**Fig. 3:**
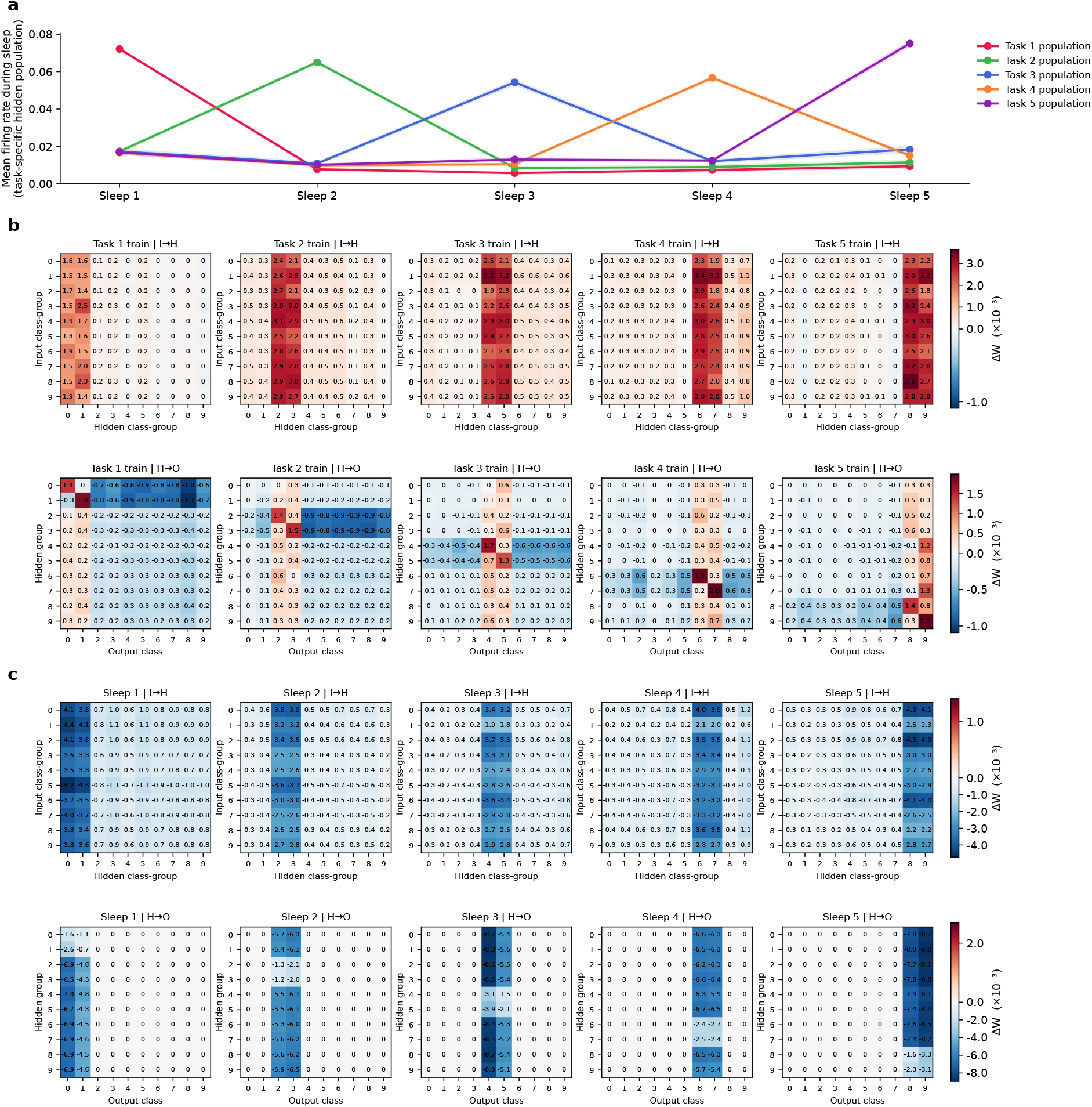
Sleep reactivates the most recently trained task’s population and reorganizes the synapses onto it. **(a)** Mean firing rate of each task’s fixed hidden-neuron population (the union of that task’s two class-specific top-10% most-active sets, computed once at the end of that task’s training and held fixed thereafter) during each of the five sleep episodes, mean ± SD across six independently trained trials. Each task’s population peaks at its own sleep episode and is comparatively silent during every other episode: sleep-driven activity is targeted to the population representing the most recently learned task rather than diffusely engaging the whole network. **(b)** Weight changes during training of each task (Δ*W* = end minus start of that task’s training; Task 1 starts from the initial weights) organized by task-selective groups. Top: input-to-hidden (I → H) block matrices; rows are input class-groups (top-75 most class-discriminative pixels per digit), columns are hidden class-groups (class-specific top-10% most-active sets, as in a). Bottom: hidden-to-output (H → O) block matrices; rows are hidden class-groups, columns are output classes. The shift toward excitation (positive Δ*W*) is concentrated on projections from all input or hidden groups onto the currently-training task’s hidden group and output units. **(c)** Weight changes during each of the five sleep episodes (Δ*W* = post-minus pre-sleep), same layout as (b). At each sleep stage, the negative shift of Δ*W* is concentrated on the synapses onto the hidden group and output units of the most recently trained task (vertical bands); within these, older-task groups shift more strongly than the own-task group. (b,c) Trial 0; each row has its own diverging colour scale centred at zero; cell values and colour bars in units of Δ*W* × 10^*−*3^.

**Fig. 4:**
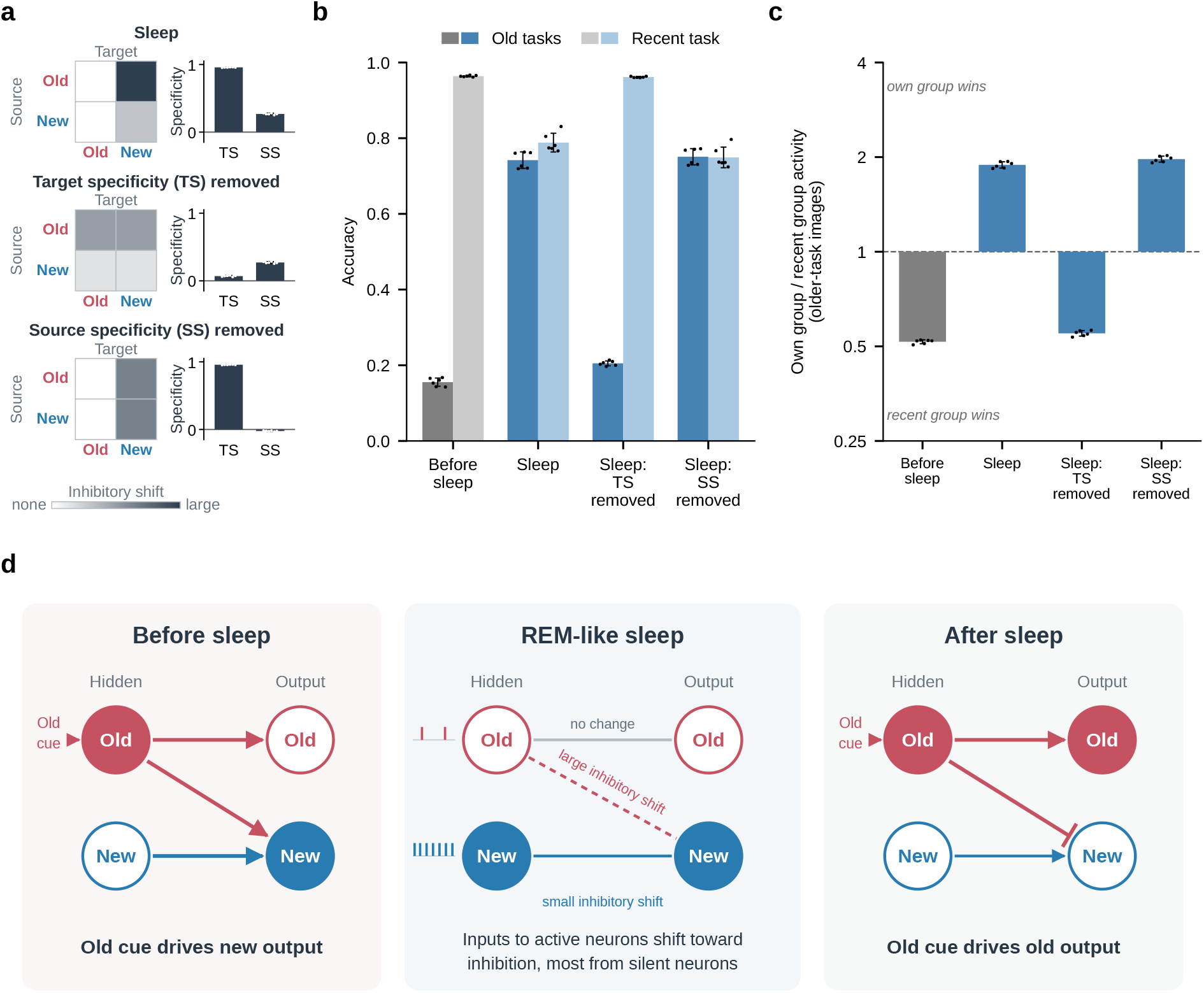
Recovery requires target specificity of sleep’s synaptic changes. **(a)** Weight changes applied in each condition, schematized as a 2 × 2 grid of source (rows: older-task or recent-task presynaptic population) by target (columns: older-task or recent-task postsynaptic population); shading indicates the size of the shift toward inhibition. Sleep: the shift is concentrated onto the recent (new) target population and is larger from old than from new sources. Target specificity removed: each presynaptic unit’s mean change onto the recent population is equalized with its mean change onto all other units. Source specificity removed: each postsynaptic unit’s mean change from older-task inputs is equalized with that from its own-task inputs. Both manipulations use sleep’s own weight changes and preserve each synapse’s deviation from its group mean. Insets: target specificity (TS) and source specificity (SS) of the weight changes applied in each condition, averaged over the I → H and H → O pathways. Both are normalized differences, (*a* − *b*)*/*(|*a*| + |*b*|), between the mean shift toward inhibition (−Δ*W*) of two groups of synapses: for TS, synapses onto the recent versus onto the older populations; for SS, synapses onto the recent population from older versus from own-task populations. Both range from −1 to 1 (0: both groups change equally; 1: the whole shift is in the first group) and are therefore shown on the same axis. **(b)** Accuracy on older tasks (dark) and on the most recent task (light) before sleep (grey), after sleep, and after sleep with target specificity (TS) or source specificity (SS) removed from its weight changes (blue). **(c)** Intrusion index *Q* (as in Fig. 2d) for the same conditions (log scale). **(d)** Summary. Before sleep, an older-task cue drives the new task’s output. During REM-like sleep, the new task’s population is reactivated while older populations are nearly silent; the synapses onto the active (new) neurons shift toward inhibition, most strongly those from silent (older) neurons, while the synapses onto older neurons do not change. After sleep, the older-task cue drives its own output. (a–c) Pooled over Sleeps 2–5, mean ± SD over six trials; dots, individual trials.

**Fig. 5:**
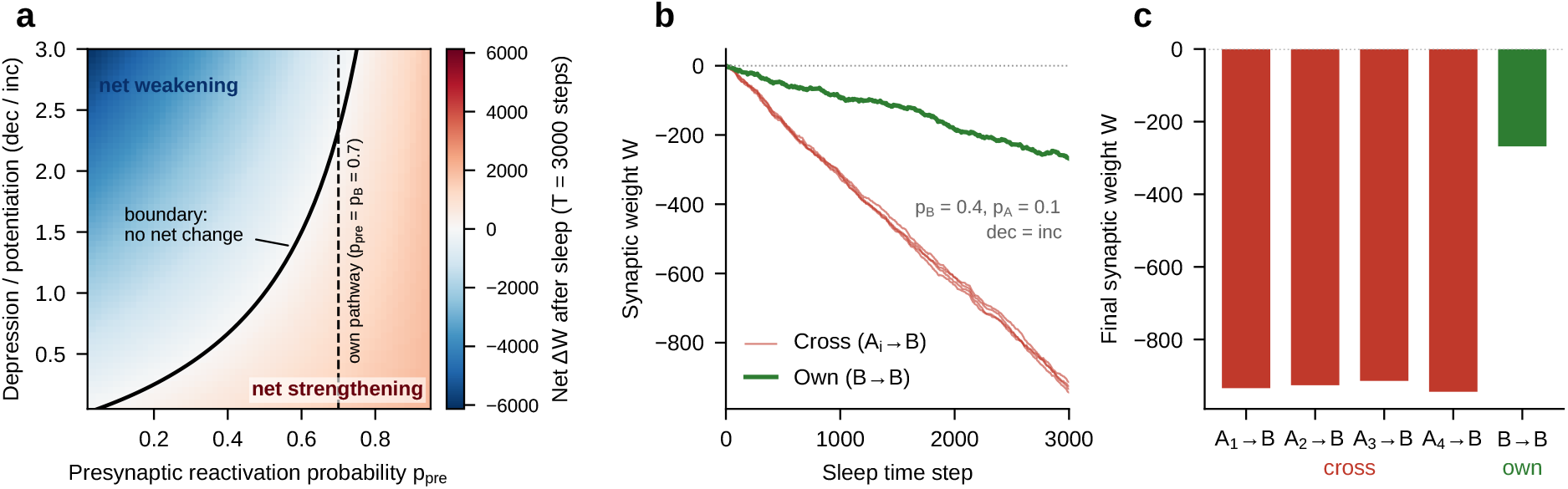
A minimal analytic model reproduces the synaptic reorganization of sleep. **(a)** Analytic phase diagram of Eq. 3: net synaptic weight change after replay (*T* = 3000 steps) as a function of the presynaptic partner’s reactivation probability *p*_pre_ and the plasticity depression/potentiation ratio dec*/*inc, for a fixed dominant firing probability *p*_*B*_ = 0.7. The solid curve marks the analytic depression boundary (Eq. 4); the dashed line marks where an own-pathway synapse sits (*p*_pre_ = *p*_*B*_). **(b)** Simulated weight trajectories over 3000 replay timesteps for one representative parameter setting (*p*_*B*_ = 0.4, *p*_*A*_ = 0.1, dec = inc): four cross-pathway synapses (*A*_*i*_ → *B*, red) versus one own-pathway synapse (*B* → *B*, green). **(c)** Final weights from the same run: cross-pathway synapses are strongly depressed while the own-pathway synapse is only weakly depressed, reproducing the asymmetry seen in the full model (Fig. 3c).

**Fig. 6:**
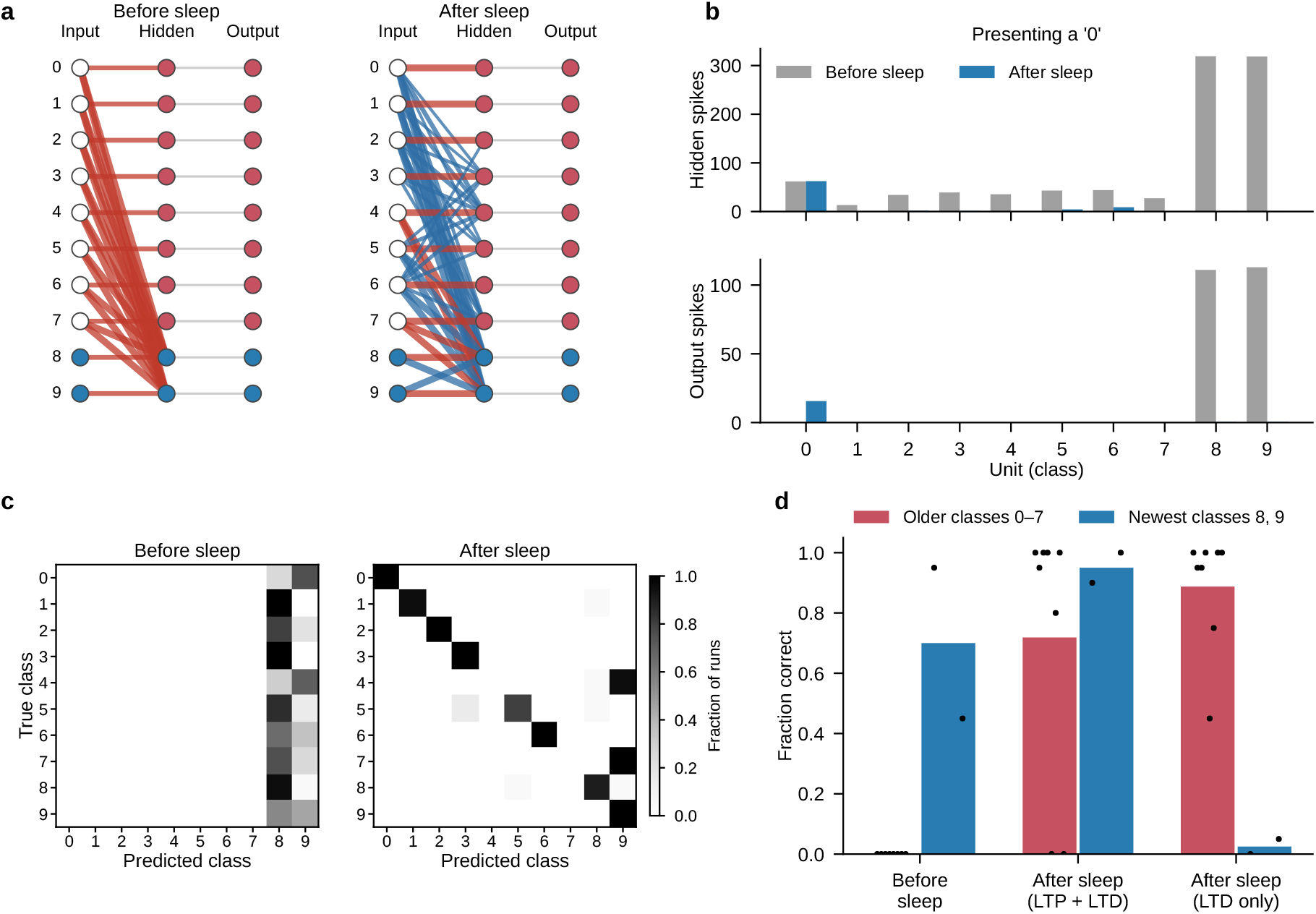
A minimal spiking network makes the mechanism explicit. **(a)** Network with one unit per digit class in each layer, showing input-to-hidden weights before (left) and after (right) one sleep episode (red, excitatory; blue, inhibitory; line width, weight magnitude; weights below 1.5 × 10^*−*3^ not drawn). Before sleep, every older-task input excites the hidden units of the most recent task (8, 9). **(b)** Spike counts of hidden (top) and output (bottom) units when a 0 is presented before (grey) and after (blue) sleep. **(c)** Classification before and after sleep: fraction of runs in which each presented digit (rows) was assigned to each class (columns). **(d)** Fraction correct for older (0–7) and most recent (8, 9) classes before sleep, and after sleep with the full plasticity rule (LTP + LTD) or with depression only (LTD only); bars, mean over classes; dots, individual classes. Input firing probabilities from class-average MNIST images; hidden-to-output weights from the trained network; sleep dynamics and plasticity as in the full model; 20 runs per condition (Methods).

**Fig. 7:**
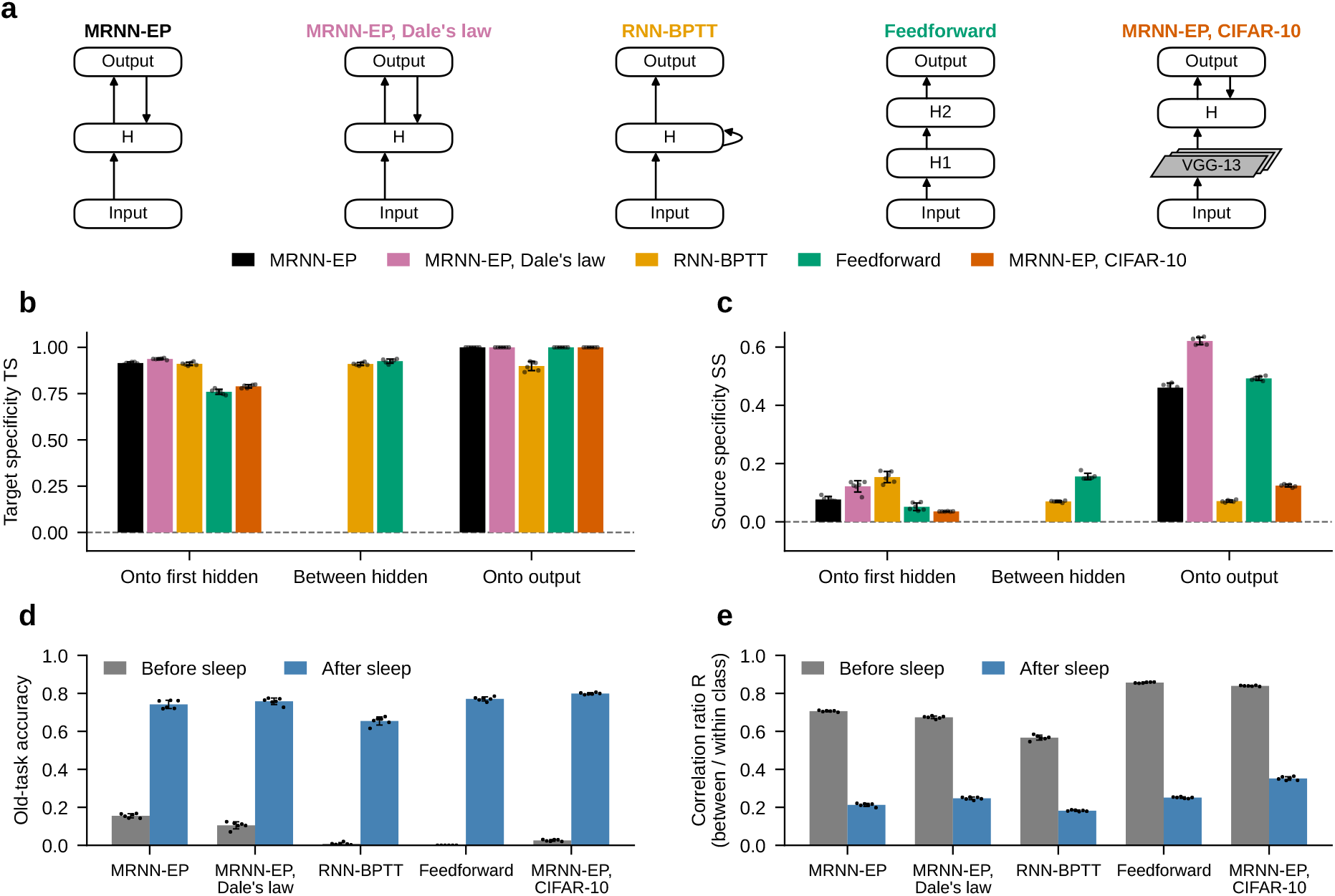
Target specificity, recovery and decorrelation generalize across models. **(a)** Architectures: MRNN-EP (main model), MRNN-EP with synapse-level Dale’s law (50%/50% excitatory/inhibitory), a recurrent network trained with backpropagation through time (RNN-BPTT; hidden-to-hidden recurrence, no output feedback), a feedforward network with two hidden layers (H1, H2) trained with backpropagation and consolidated with SRC, and MRNN-EP trained on CIFAR-10, whose input passes through pretrained convolutional layers of VGG-13 that are held fixed (only the EP network learns and sleeps). All other models were trained on the same five-task MNIST curriculum. **(b)** Target specificity (TS) of sleep’s weight changes for each pathway grouped by depth: onto the first hidden layer (I → H, I → H1), between hidden layers (RNN-BPTT H → H; feedforward H1 → H2) and onto the output layer. **(c)** Source specificity (SS) for the same pathways. **(d)** Accuracy on older tasks before (grey) and after (blue) sleep. **(e)** Correlation ratio *R* of the last hidden layer before and after sleep. (b–e) Per trial, mean over Sleeps 2–5; mean ± SD over six trials; dots, individual trials.

### 2.1 REM-like sleep rescues older memories from interference

Our main model, MRNN-EP, is a recurrent neural network (RNN) trained with Equilibrium Propagation (EP). Each training input is processed in two phases: a free phase, in which the network settles to a stable state without a target, and a weakly clamped phase, in which the output units are gently nudged toward the correct label [17, 22]. Weights are updated by a local rule that compares neuron activity between the two phases, without backpropagating errors. We use a simplified version of the predictive learning rule of Luczak et al. [23], in which the predicted signal is replaced by the actual free-phase activation

[24] (Methods). MRNN-EP lacks the biophysical detail of more realistic models, but it can learn many complex memories with local plasticity alone, which our question requires.

To illustrate catastrophic forgetting in an RNN trained with EP, and the effect of sleep-like reactivation in enabling continual learning, we started with the simple 10-class MNIST dataset. We trained a 1024-unit RNN sequentially on five binary MNIST digit-classification tasks (digit pairs 0/1, 2/3, 4/5, 6/7, and 8/9), each followed by a REM-like sleep episode (Fig. 1a,b). The question we ask is whether such an episode can separate the overlapping representations of older and recent memories (Fig. 1a).

To implement sleep, we mapped the trained MRNN-EP onto a spiking network of identical structure with scaled weights, following the Sleep Replay Consolidation (SRC) approach developed for feedforward networks [25]. The spiking network was driven by Poisson spike trains generated from the input statistics of the most recently trained task only, with no labels or targets; unless stated otherwise, sleep was driven this way in all models. Weights changed by the local rule described above: coincident pre- and postsynaptic spikes increased a weight, and postsynaptic spikes without presynaptic input decreased it. We call a decrease of a weight (Δ*W <* 0) depression or a shift toward inhibition, and an increase (Δ*W >* 0) potentiation or a shift toward excitation. Because weights in the main model can be positive or negative, a decrease either weakens an excitatory connection or strengthens an inhibitory one, shifting the excitation/inhibition balance of the postsynaptic neuron’s input. After sleep, the weights were mapped back onto the original network to test performance on all tasks learned so far.

Figure 1c shows mean validation accuracy across six trials for each of the five MNIST tasks, plotted at the end of every training and sleep phase. A characteristic pattern of *forgetting and recovery* is evident at every task transition. After training on a new task, the classification accuracy of all previously learned tasks falls substantially, often close to zero, consistent with catastrophic interference in shared synaptic weights. Following each sleep episode, however, accuracy for the task that had just been trained is preserved or decreases slightly, while the accuracy of earlier tasks shows partial or full recovery. To quantify this recovery, for each Sleep *k* (*k* = *2* –*5*), which follows the training of Task *k*, we averaged the accuracy of Tasks *1* to *k* − *1* before vs. after sleep and then averaged over the four sleeps. This old-task accuracy rose from 15.5 ± 1.1% immediately before sleep to 74.2 ± 2.2% immediately after sleep (mean ± SD over six trials). After the final sleep (Sleep 5), mean accuracy over all five tasks, including the most recently trained Task 5, was 70.6 ± 3.2%. The magnitude of sleep-related recovery diminishes for older tasks over successive task transitions, reflecting the accumulating interference from multiple overwriting steps, yet a measurable benefit of sleep remains detectable across all five tasks. The small standard deviations across six trials show that the recovery is robust.

To understand how sleep affects task-specific predictions, we examined the confusion matrices before and after the final sleep phase, averaged over six independent runs. Before the final sleep, predictions were largely biased toward the most recent task: almost all input digits were classified as digits 8 or 9 from the last trained task (Fig. 1d, left). After the final sleep, the models successfully retained and recalled knowledge from previous tasks (Fig. 1d, right).

### 2.2 Sleep decorrelates class representations and separates task populations

To understand the neural mechanisms underlying the observed performance recovery during sleep, we analyzed the hidden-layer activation patterns recorded on validation data at the end of every training and every sleep epoch. We quantified three complementary properties: (i) inter-class correlation structure, (ii) the overlap between the neuronal populations recruited by different classes, and (iii) whether inputs of an older task drive their own neurons or are captured by the neurons of the most recent task.

Figure 2a depicts the correlation matrices of activations in the hidden layer for the MRNN-EP model on the MNIST dataset, prior to and following the last sleep application. After sleep, the hidden-layer representations of different classes were much less correlated, indicating that sleep helps disentangle class-specific features and reduce task interference.

To summarize this across all training and sleep stages, we defined two statistics: the *within-class* correlation (diagonal; how consistently the same digit activates the same neurons across different samples) and the *between-class* correlation (off-diagonal; how similar the representations of different digits are). Sleep reduces both correlations, but the reduction is substantially larger for the between-class correlation (Supplementary Fig. S1c), so sleep mainly removes overlap between classes rather than scaling all correlations down. The ratio *R* = off-diagonal*/*diagonal combines these two measures into a single index of representational discrimination: *R* ≈ 1 implies indiscriminate coding, *R* ≪ 1 implies class-selective coding. Because between-class correlation falls faster than within-class correlation, *R* decreases markedly after each sleep episode, from 0.66–0.80 at the end of the five training blocks to 0.14–0.30 after the corresponding sleep episodes (Fig. 2b). These observations suggest that sleep helps allocate distinct neurons to different tasks, which would promote independent class representations.

Next, we asked whether this decorrelation reflects a shift from shared to largely separate neuronal populations. For each class, we defined its population as the 10% of hidden neurons most active on that class’s validation data, recomputed after every training block and every sleep episode so that overlap could be tracked over time. For each phase, we computed the Jaccard index between the populations of every pair of classes, i.e. the fraction of shared neurons relative to the union of the two populations, and summarized it as the mean over all class pairs. After each sleep episode, the mean pairwise overlap dropped substantially, from 0.37–0.46 at the end of the training blocks to 0.09–0.16 after the corresponding sleep episodes (Fig. 2c). The full pairwise overlap matrices show that before the final sleep, off-diagonal entries are uniformly high, whereas after sleep they drop towards 0.1–0.2, consistently across all six trials (Supplementary Fig. S1a).

Consistent with this, the fraction of each class’s population that is not shared with any other class – the *exclusive fraction* – rose from ~ 10% at training end to ~ 25% after sleep on average (Supplementary Fig. S1b). The only exception was class “1”, which maintained a high fraction of exclusive neurons even after each new task training. This effect is likely related to digit 1’s distinctive pixel-level statistics: it has both the sparsest raw pixel representation of any MNIST digit (the highest fraction of near-zero pixels) and the lowest mean pixel-intensity correlation with every other digit, suggesting its input pattern is intrinsically the most separable from the rest of the digit distribution even before training begins.

Decorrelated, separated populations predict a specific functional consequence: after sleep, inputs of an older task should again drive their own neurons, rather than being captured by the neurons of the most recently trained task. To test this directly, for every sleep episode *k* ≥ 2 and every older task *j < k* we presented task-*j* validation images and computed *Q*, the mean activity of task *j*’s own hidden population divided by the mean activity of the most recent task *k*’s population. Each task’s population was a fixed hidden-neuron group – the union of that task’s two class-specific top-10% most-active sets, computed once at the end of that task’s training and held fixed thereafter (Methods). *Q <* 1 means that the recent task’s neurons respond more strongly to older-task inputs than the older task’s own neurons do. Before sleep this was the case at every sleep episode: pooled over Sleeps 2–5, *Q* = 0.52 ± 0.01, i.e. older-task images drove the recent task’s population about twice as strongly as their own (Fig. 2d). After sleep, *Q* rose to 1.89±0.04 (increase in all six trials at every sleep episode; exact paired Wilcoxon *p* = 0.031, the minimum attainable with six trials), and the same reversal was seen when the neurons belonging to both populations (on average 21 of about 160 units per population) were excluded from both before computing *Q* (0.43 before vs. 2.22 after sleep). Together, these results show that sleep gives each task a largely separate set of neurons and lets each task’s inputs drive that set again, reducing the overlap that causes interference between tasks.

### 2.3 Sleep reactivates the most recent task’s population and reorganizes the synapses onto it

How does sleep-like replay lead to decorrelation of memory representations? We first asked whether activity during sleep equally engaged all network neurons or was specific to neurons previously involved in a given trained task. For each of the five tasks, we used the fixed hidden-neuron group defined above (the same groups as in Fig. 2d) and measured its mean firing rate during each sleep episode. Activity during sleep was dominated by the group of the task that had just been trained. Pooled across all sleep episodes and six trials, this group fired at a mean rate of 0.065, compared with 0.012 for the other tasks’ groups during the same episode, a more than fivefold difference present in every episode (Fig. 3a). While the noisy input to the network remained constant, overall activity in the hidden and output layers progressively decreased over the course of each sleep episode (Supplementary Fig. S2).

While this is clearly a signature of reactivation, it does not answer the question of how this reactivation changes synaptic connectivity between tasks. Indeed, the classic account of NREM reactivation holds that task-specific firing order strengthens synapses among neurons of the most recently learned task, further augmenting that representation. Given that the last task is already dominant immediately after training, this account alone does not explain how REM-like sleep could achieve the opposite goal – recovering older tasks from forgetting.

To answer this, we analyzed the changes of synaptic weights between groups of neurons representing each learned task, both during training and during sleep. For the *hidden-to-output* (H → O) pathway, hidden neurons were grouped into the ten class-specific top-10% most-active sets defined above (two per task). Each cell of the resulting 10 × 10 block matrix reports the mean Δ*W* from that hidden group to the corresponding output unit. For the *input-to-hidden* (I → H) pathway, input pixels were grouped by discriminative relevance (top 75 pixels per class), and weight changes to each hidden group were aggregated analogously.

Why does new training cause catastrophic forgetting? During training of each task (Fig. 3b), the synapses onto that task’s hidden group shifted toward excitation from almost all input pixel groups, not only from the task’s own discriminative pixels, producing a vertical band of positive Δ*W* at the task’s hidden columns. The H → O pathway showed the same pattern: synapses from all hidden groups onto the current task’s output units shifted toward excitation. At the same time, synapses from the current task’s hidden group onto the output units of all other tasks shifted toward inhibition, producing a horizontal band of negative Δ*W*. The new hidden representation thus recruits input from all pixel groups and suppresses the outputs of other tasks, “stealing” input and output weight from previously learned tasks. This is the synaptic basis of the catastrophic forgetting in Fig. 1.

Sleep reverses this pattern (Fig. 3c). During sleep, most weights shifted toward inhibition, weakening excitatory and strengthening inhibitory connections, while only a small fraction shifted toward excitation (Supplementary Fig. S3). This shift was not uniform: it was concentrated on the synapses onto the population of the task that had just been trained – the population that dominated reactivation – producing, at every sleep stage, vertical bands of negative Δ*W* at the recent task’s hidden columns (I → H) and output columns (H → O). In the H → O pathway, synapses onto the output units of older tasks did not change at all, and within the recent task’s output columns the shift was far stronger from older-task hidden groups than from the recent task’s own group (e.g., sleep 5: −7.3 to −9.0 for older-task groups vs. −1.6 to −3.3 for the own group, in units of 10^*−*3^). In the I → H pathway, almost all connections between input and hidden groups shifted toward inhibition. The shift was strongest from older-task input groups onto the hidden group of the most recent task (e.g., sleep 3: input groups 0, 2, 3, and 6 onto hidden groups 4/5, Δ*W* ≈ −3.1 to −3.7 × 10^*−*3^), weaker from the recent task’s own pixels onto its own hidden group (Δ*W* ≈ −2.4 to −2.8 × 10^*−*3^), and weakest onto the hidden groups of other tasks (Δ*W* ≈ −0.1 to −0.8 × 10^*−*3^). As a result, after sleep, older-task populations suppress the neurons of the most recent task, isolating the memory traces.

This pattern is robust and follows directly from the local plasticity rule. The asymmetry between older-task and own-task groups onto the recent task’s output units was present in all 30 of 30 trial × sleep-stage combinations (mean Δ*W* from older-task groups −0.0055 to −0.0086 vs. from the own-task group −0.0009 to −0.0033; the two ranges never overlapped). At the level of individual synapses, Δ*W* tracked the joint spiking statistics of the pre- and postsynaptic neurons, with synapses from older-task sources onto the recent task’s neurons experiencing more “postsynaptic spike without presynaptic spike” events than synapses from the recent task’s own sources (Supplementary Fig. S6). No rule that singles out old tasks is needed: the same local plasticity rule, acting on the activity that arises during sleep, produces this pattern.

### 2.4 Recovery requires that sleep’s changes target the inputs to the reactivated population

The block matrices of Fig. 3c contain two distinct kinds of specificity (Fig. 4a). *Target specificity* : the shift toward inhibition is concentrated on synapses onto the population of the recent task – the population reactivated during sleep – and not onto the populations of older tasks (the vertical bands). *Source specificity* : among the synapses onto the recent task’s population, those from older-task sources shift more than those from the recent task’s own sources (the gradient within the bands). We quantified both properties for each pathway and sleep episode *k* from the per-synapse shift toward inhibition, *s* = −(*W*_post_ − *W*_pre_), averaged over the synapses between task populations (Methods). Target specificity compares the mean shift onto the recent population, 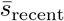, with that onto the older populations,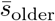:

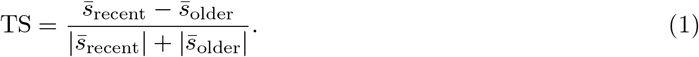

Source specificity compares, among the synapses onto the recent population only, the mean shift from older-task sources, 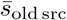, with that from the recent task’s own sources, 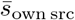:

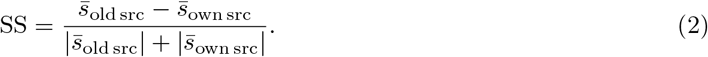

Both indices range from −1 to 1. TS = 1 means that only synapses onto the recent population shifted, and TS = 0 that synapses onto recent and older populations shifted equally; SS *>* 0 means that inputs from older tasks shifted more than the recent task’s own inputs, and SS = 0 that all inputs onto the recent population shifted equally. Sleep’s changes were almost perfectly target-specific (TS = 0.92 ± 0.01 in the I → H and 1.00 in the H → O pathway) and moderately source-specific (SS = 0.08 ± 0.01 in the I → H and 0.46 ± 0.02 in the H → O pathway; mean ± SD over six trials, Sleeps 2–5).

To determine which form of specificity was required for recovery, we modified the weight changes produced by sleep and applied them to the pre-sleep network (Methods). Each manipulation removed either target or source specificity by equalizing the relevant group means, while preserving each unit’s total weight change and each synapse’s deviation from its original group mean. Removing target specificity abolished recovery. In this manipulation, each presynaptic unit’s mean change onto the recent population was equalized with its mean change onto all other units (TS: 0.96 → 0.07, averaged over both pathways; SS unchanged). Pooled over Sleeps 2–5, old-task accuracy was then 0.21 ± 0.01, barely above the 0.16 before sleep and far below the 0.74 after real sleep (Fig. 4b), and *Q* stayed at 0.55 (Fig. 4c). Removing source specificity left recovery intact. Here, each postsynaptic unit’s mean change from older-task inputs was equalized with its mean change from own-task inputs (SS: 0.27 → −0.02; TS unchanged), and old-task accuracy (0.75 ± 0.02) and *Q* (1.97) were as high as after real sleep. Its only cost was to the recent task, whose accuracy fell from 0.79 to 0.75 (all six trials). Removing target specificity from only one of the two pathways reduced recovery partially (old-task accuracy 0.50 when only the H → O and 0.36 when only the I → H pathway was equalized; Supplementary Fig. S4); randomly redistributing sleep’s own weight changes across synapses likewise reduced or abolished recovery (Supplementary Fig. S5).

Thus the property of sleep’s synaptic changes that rescues old memories is target specificity: plasticity concentrates on the inputs to the reactivated, most recently trained population and shifts them toward inhibition, so that this population no longer captures the inputs of older tasks (Fig. 4d). Source specificity – the stronger shift from older than from own sources – is present, but it mainly spares the recent task’s own pathway. In the current model, a shift toward inhibition can come from depression of excitatory synapses or from potentiation of inhibitory ones. Section 2.7 confirms our conclusion with a Dale’s-law model in which only excitatory synapses could change during sleep. Its weight changes were target-specific but not source-specific, and older tasks still recovered about as well as in the fully plastic model.

### 2.5 A minimal analytic model explains target and source specificity

To see why local plasticity acting on biased reactivation is enough, consider a minimal analytic model without any network architecture. It has two kinds of binary units, redrawn independently at every time step of replay: a *dominant* population *B*, representing the most recently trained task, whose units fire with probability *p*_*B*_, and one or more *quiescent* populations *A*_*i*_, representing older tasks, whose units fire with a lower probability *p*_*A*_ ≪ *p*_*B*_. Reactivation is biased toward the most recent task but is otherwise random; no replay of specific sequences or patterns is assumed. A synapse from a presynaptic unit to a postsynaptic unit is updated only on timesteps when the *postsynaptic* unit fires, exactly as in the Hebbian plasticity rule of Algorithm 1: coincident firing potentiates the synapse by a fixed amount *inc*, while postsynaptic firing with a silent presynaptic partner depresses it by a fixed amount *dec*. No structure is imposed beyond this: which units belong to which population, and how often each population fires, are the only free quantities.

For a synapse from a presynaptic unit with firing probability *p*_pre_ onto a postsynaptic unit in *B*, the expected weight change per timestep of replay follows directly from the two ways a postsynaptic spike can be accompanied by presynaptic activity or silence:

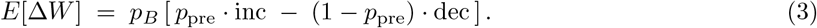

This expression separates the two properties identified above. Its magnitude is set by the *postsynaptic* firing probability. The same expression holds for a synapse onto a unit of a quiescent population *A*, with *p*_*B*_ replaced by *p*_*A*_: because *p*_*A*_ ≪ *p*_*B*_, synapses onto older populations are updated rarely and change little, whereas synapses onto the reactivated population *B* change at the much higher rate *p*_*B*_ – target specificity. Its sign is set by the *presynaptic* firing probability: the expectation is negative – net depression – whenever

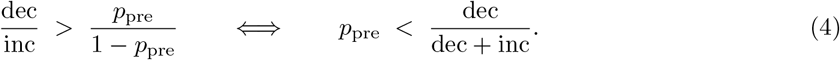

This threshold depends only on the *presynaptic* partner’s reactivation probability and the fixed ratio *dec/inc*. It does not depend on *p*_*B*_, which sets only how often updates occur, or on which task or population is involved. Applying this to the two cases of interest: an *own-pathway* synapse, whose presynaptic partner is itself drawn from the dominant population (*p*_pre_ = *p*_*B*_), sits close to or beyond this threshold and depresses weakly or not at all, while a *cross-pathway* synapse, whose presynaptic partner is drawn from a quiescent old-task population (*p*_pre_ = *p*_*A*_ ≪ *p*_*B*_), sits well below the threshold and depresses strongly – source specificity. The same rule, applied to units that differ only in how often they happen to be reactivated, is therefore sufficient to generate both properties, without any explicit rule that treats “own” and “cross” synapses differently.

Figure 5a visualizes Eq. 3 directly as a phase diagram over *p*_pre_ and *dec/inc*, with the boundary of Eq. 4 overlaid; own- and cross-pathway synapses occupy well-separated regions of this diagram simply because they sit at different values of *p*_pre_. Figure 5b,c confirm this with a direct stochastic simulation of the model above (four old-task populations converging on one dominant population, *p*_*B*_ = 0.4, *p*_*A*_ = 0.1, *dec* = *inc*, 3000 replay timesteps, no fitted parameters): cross synapses drift to strongly negative weights while the own synapse drifts only weakly negative, reproducing the qualitative asymmetry reported for the full model in Fig. 3c from a model with no trained weights, no task structure, and no free parameters beyond the reactivation probabilities and plasticity ratio themselves.

To test how well this minimal model describes the full network, we measured, directly from the spike trains recorded during Sleeps 2–5, the probability that the presynaptic unit was active in the time step before a postsynaptic spike,

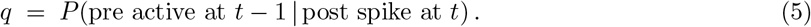

Eq. 3 assumes that pre- and postsynaptic spikes are independent, in which case *q* equals the presynaptic firing probability *p*_pre_. This assumption held for input-to-hidden synapses (*q* ≈ *p*_pre_), whose Poisson input does not depend on network activity, but not for hidden-to-output synapses, where *q* was about three times higher than *p*_pre_ because hidden spikes drive the output units one step later. With *q* in place of *p*_pre_, Eq. 3 reproduced every synapse’s weight change exactly, and all measured own and cross synapses lay on the depression side of the boundary (*q* = 0.01–0.20, below dec*/*(dec+inc) = 0.32), with cross below own in both pathways (Supplementary Fig. S6). Correlated firing therefore does not change the regime: own and cross synapses still weaken. It changes only how much own-pathway hidden-to-output synapses weaken. Because these hidden units help drive the output units they project to, their synapses weaken only about half as much as the independence assumption predicts. As a result, the difference between cross and own synapses in this pathway is about three times larger than predicted by the model assuming independent pre- and postsynaptic firing.

This is reactivation-targeted pruning: plasticity is concentrated on the reactivated population, where neurons that fire while their inputs are silent lose those inputs.

### 2.6 A minimal spiking network shows how sleep restores older memories, and why the most overlapping ones resist

To follow the mechanism from synapses to network dynamics, we built a minimal spiking network with one unit per digit class in each layer (10 input, 10 hidden and 10 output units; Fig. 6a; Methods). Each input unit fired with a probability given by the pixels most active for its class in the class-average MNIST image, so digits that share pixels also share input activity. Input-to-hidden weights were set to reproduce the interference left by waking training: each input unit excited its own hidden unit, and every older-task input unit excited the hidden units of the most recent task (digits 8 and 9) 1.5-fold more strongly. Hidden-to-output weights were taken from the trained network, and sleep used the same spiking dynamics, plasticity rule and parameters as the full model, driven by the most recent task’s average input.

Before sleep, every digit drove hidden units 8 and 9 and was classified as 8 or 9 (Fig. 6b,c). During sleep, the most recent task’s hidden units dominated activity, so the plasticity rule depressed the synapses onto them – most strongly those from older-task inputs, which were mostly silent when units 8 and 9 fired – and turned most interference synapses inhibitory (Fig. 6a). After a single sleep episode, presenting a 0 activated hidden unit 0 and output 0 instead of outputs 8 and 9 (Fig. 6b), and the network correctly classified 72% of older-class presentations, and 95% of the most recent ones (Fig. 6c,d). With depression only (no potentiation term), sleep still restored the older classes (89%) but nearly erased the most recent ones (3%; Fig. 6d). Depression of the inputs to the reactivated population therefore restores old memories, and potentiation of that population’s own inputs protects the newest one.

The minimal network also exposed a limit of the mechanism. Digits 4 and 7, whose pixels overlap most with digit 9 of the most recent task, remained classified as 9 after sleep (Fig. 6c): because their input units were often co-active with the reactivated population during sleep, potentiation offset depression on their synapses onto it, and their interference synapses stayed excitatory (Fig. 6a). The full network showed the same weakness in milder form. After the final sleep, digit 4 was the least recovered of all older digits (36 ± 16% correct, mean ± SD over six trials), and half of its presentations (49%) were still classified as 9; digit 7 was most often confused with 9 (15%). Sleep therefore separates memories most effectively when they overlap little with the reactivated one: inputs that share the most features with the most recent memory are the hardest to disentangle.

### 2.7 The mechanism holds under a synapse-level Dale’s law, and excitatory plasticity alone is sufficient

A biological neuron releases either an excitatory or an inhibitory neurotransmitter, so all of its outgoing synapses have the same sign. Moreover, while depression of excitatory synapses during sleep is well documented, direct evidence for potentiation of inhibitory synapses during sleep is less consistent (Discussion). It is therefore important to ask whether the mechanism survives sign-constrained synapses and, in particular, whether it requires inhibitory plasticity at all.

Enforcing Dale’s law literally in the MRNN-EP architecture is not possible, however. EP’s local, energy-based update rule is only a valid estimator of the true gradient when the network’s weight matrix is symmetric (*W*_*ij*_ = *W*_*ji*_), since the same connectivity must define both the free-phase and nudged-phase fixed points from which the update is computed. A symmetric weight matrix ties *W*_*ij*_ and *W*_*ji*_ together, which is only consistent with a neuron-level Dale’s law when the two neurons sharing a connection have matching identities (both excitatory or both inhibitory): wherever an excitatory neuron connects to an inhibitory one, Dale’s law requires *W*_*ij*_ ≥ 0 (set by the excitatory source) and *W*_*ji*_ ≤ 0 (set by the inhibitory source), which directly contradicts *W*_*ij*_ = *W*_*ji*_ for any nonzero connection.

As the closest structurally compatible approximation, we instead imposed Dale’s law at the level of individual synapses rather than individual neurons: each entry of the weight matrix was independently assigned a fixed excitatory or inhibitory sign at initialization, held fixed throughout training and sleep, with the effective weight computed as *W*_eff_ = |*W*_raw_| × sign at every forward pass (Supplementary Section S2). This sacrifices the biological requirement that a synapse’s sign follow its source neuron’s identity, but preserves the structural property that ultimately matters for the mechanism under test: every connection has a fixed, non-switchable sign throughout learning. With sleep hyperparameters re-optimized by genetic algorithm for this model (50%/50% excitatory/inhibitory synapses), it reached 73.9 ± 1.7% mean task accuracy after the final sleep across 6 independently trained trials, comparable to the unconstrained recurrent model of the main text. All signatures of the main model were reproduced (Fig. 7, pink; Supplementary Figs. S7, S8): old-task accuracy rose from 0.11 ± 0.02 before to 0.76 ± 0.02 after sleep (pooled over Sleeps 2–5), *R* fell from 0.67 to 0.25, *Q* rose from 0.55–0.64 before to 1.40–2.12 after each sleep, and sleep’s weight changes were target-specific (TS = 0.94 for I → H and 1.00 for H → O) and source-specific (SS = 0.12 for I → H and 0.62 for H → O).

The Dale’s-law model allows us to separate the contributions of excitatory long-term depression (LTD) and inhibitory long-term potentiation (LTP). We trained two additional models, identical to the one above except for which synapses were allowed to change during sleep: in the first, only excitatory synapses were plastic and inhibitory synapses were held fixed; in the second, only inhibitory synapses were plastic and excitatory synapses were held fixed. Sleep hyperparameters were independently re-optimized by genetic algorithm for each model (Methods). Excitatory plasticity alone, in which sleep could change only excitatory synapses, restored old tasks about as well as the fully plastic model: averaged over Sleeps 2–5, old-task accuracy rose from 0.08 ± 0.02 to 0.74 ± 0.02, compared with 0.11 → 0.76 for the fully plastic and 0.11 → 0.76 for the inhibitory-only model (Supplementary Table S1). Old-task recovery matched or exceeded that of the fully plastic model at Sleeps 2–4 (0.78 vs. 0.76 on average) and fell behind only at Sleep 5 (0.63 vs. 0.76), so its final accuracy measured after Sleep 5 was somewhat lower (66.9 ± 1.6%, vs. 73.9 ± 1.7% for the fully plastic and 72.7 ± 2.1% for the inhibitory-only model). Both ablations reproduced the representational signatures (*R*: 0.71 → 0.25 with excitatory-only and 0.69 → 0.26 with inhibitory-only plasticity; Supplementary Figs. S9, S11) and the vertical bands of weight change onto the recently reactivated population (Supplementary Figs. S10, S12).

The excitatory-only model also provides an independent test of the conclusion drawn from Fig. 4. Its weight changes were as target-specific as those of the other models (TS = 0.89 for I → H and 1.00 for H → O), but had essentially no source specificity (SS = 0.03 ± 0.02 for I → H and 0.01 ± 0.04 for H → O, vs. 0.12 and 0.62, respectively, in the fully plastic model; Supplementary Table S1): all excitatory inputs onto the reactivated population were depressed to a similar extent, regardless of whether they came from older or own-task neurons. Nevertheless, recovery of old tasks, averaged over Sleeps 2–5, was undiminished – exactly as predicted by the manipulation in which source specificity was removed from the main model’s weight changes (Fig. 4). Reactivation-targeted pruning therefore requires neither inhibitory plasticity nor source specificity.

### 2.8 The mechanism generalizes across architectures and learning rules

The mechanistic account above was derived from recurrent, EP-trained models. To test whether its signatures are general properties of sleep replay, rather than artifacts of this specific recurrent architecture or its local EP training rule, we analyzed several further models, first on the same five-task MNIST curriculum and then on natural images (Fig. 7a). First, Equilibrium Propagation requires a symmetric weight matrix and a feedback pathway from the output layer back to the hidden layer, both biologically motivated but neither universal among recurrent network models. We therefore trained an architecturally matched recurrent network (same layer sizes, hidden-to-hidden recurrence, no output-to-hidden feedback pathway) using standard backpropagation-through-time (BPTT) in place of EP (RNN-BPTT). Second, we applied the original SRC procedure [25] to feedforward networks with two hidden layers trained with backpropagation. Finally, to test whether the mechanism depends on the simple statistics of MNIST, we trained MRNN-EP (Fig. 7a; Supplementary Section S4) and a feedforward network (Supplementary Section S5) on five two-class tasks of CIFAR-10 natural images. Their input features came from pretrained convolutional layers that were held fixed (VGG-13 for MRNN-EP; Methods). Sleep hyperparameters were independently re-optimized by genetic algorithm for each model.

In every model, sleep restored old-task accuracy (Fig. 7d; pooled over Sleeps 2–5: MRNN-EP 0.16 → 0.74, Dale’s law 0.11 → 0.76, RNN-BPTT 0.01 → 0.66, feedforward 0.00 → 0.77, CIFAR-10 0.03 → 0.80) and decorrelated the last hidden layer (*R*: 0.71 → 0.21, 0.67 → 0.25, 0.57 → 0.18, 0.86 → 0.25, 0.84 → 0.35; Fig. 7e; all paired Wilcoxon *p* = 0.031). Sleep’s weight changes were target-specific in every model and every pathway, from the input onto the first hidden layer, between hidden layers (RNN-BPTT recurrence and the feedforward network’s H1 → H2 pathway) and onto the output (TS ≥ 0.76 in every pathway; one-sided exact Wilcoxon *p* = 0.016, the minimum attainable with six trials; Fig. 7b). Source specificity was positive in every pathway but varied between models (Fig. 7c): it increased with depth in MRNN-EP (I → H 0.08, H → O 0.46), in the Dale’s-law model (0.12, 0.62), in MRNN-EP on CIFAR-10 (0.04, 0.12) and in the feedforward network (0.05, 0.16, 0.49 for I → H1, H1 → H2 and H2 → O), but not in RNN-BPTT (0.15, 0.07, 0.07). Consistent with the manipulations of Fig. 4, models with larger source specificity onto the output did not show larger recovery: RNN-BPTT, with little source specificity (0.07), recovered as much as MRNN-EP (0.46). Full representational and synaptic analyses of these models are given in Supplementary Figs. S13–S20 (Supplementary Sections S3–S5). Finally, the mechanism did not depend on driving sleep with the most recent task alone: when the sleep input combined the statistics of all tasks learned so far, the most recent task’s population still dominated reactivation, and sleep still decorrelated representations and reorganized the synapses onto the reactivated population (Supplementary Figs. S21, S22; Supplementary Section S6).

Together, these results show that reactivation-targeted pruning – the reorganization of the synapses onto the reactivated population – and the resulting decorrelation and recovery are not specific to the recurrent, EP-trained architecture: they are reproduced under a synapse-level Dale’s law, with excitatory plasticity alone, under a gradient-based waking learning rule, in feedforward networks and on natural images.

## 3 Discussion

We identify a synaptic mechanism by which REM sleep can separate memories that would otherwise interfere with one another. Its logic runs counter to the classic view of replay. During sleep, the most recently learned memory dominated reactivation. Replay during NREM sleep is thought to strengthen the memories it reactivates; here, instead, the reactivated memory lost the synaptic input it had taken over from older memories during learning. The synapses onto the reactivated population shifted toward inhibition, most strongly those from the largely silent populations of older memories, while the synapses onto older populations remained nearly unchanged. We call this mechanism reactivation-targeted pruning. It undoes the signature of catastrophic forgetting – the capture of older inputs by the neurons of the newest memory – and thereby decorrelates class representations, separates task populations and restores the routing of older-task inputs to their own neurons, as long-standing REM sleep orthogonalization theories propose [7, 10, 11].

To reach this conclusion, we used recurrent neural networks trained with Equilibrium Propagation (EP), a biologically plausible, local learning rule, and simulated sleep with our previously proposed Sleep Replay Consolidation (SRC) algorithm [25], in which the trained network is converted into a spiking network driven by noisy input and its synapses are modified by a local plasticity rule. SRC was shown to improve continual learning in feedforward networks [25] and, more recently, in MRNN-EP on a variety of datasets, including Fashion-MNIST, Kuzushiji-MNIST, CIFAR-10 and ImageNet, where it matched or exceeded BPTT-trained recurrent networks [26]. Here, by manipulating sleep’s own weight changes, we showed that it is the targeting of the reactivated population (target specificity) that restores older memories: removing only this targeting, while keeping the overall amount of change, abolished recovery. The targeting follows directly from the plasticity rule. The most recently trained populations were the most active, so their incoming synapses were updated most often. Inputs from older-task populations were mostly silent when these neurons fired, so they experienced many “post-spike, no pre-spike” (0 → 1) events, which shifted them toward inhibition. Conversely, coincident firing (1 → 1 events) counteracted this depression on the population’s own inputs, which were often active together with it, and thereby protected the newest memory: in the minimal spiking network, removing the potentiation term still restored older memories but erased the most recent ones (Fig. 6d). Depression of excitatory and potentiation of inhibitory connections are complementary routes to this outcome, each independently sufficient: in a Dale’s-law model, excitatory plasticity alone and inhibitory plasticity alone each restored older memories. The same targeted reorganization, with the resulting decorrelation and recovery, appeared in every model we tested – recurrent networks trained with EP or with backpropagation through time, feedforward networks and networks with a synapse-level Dale’s law – and on both MNIST and CIFAR-10 natural images (Fig. 7).

Figure 8a relates the main results of the model to existing observations of REM sleep. In the model, the most recently trained task’s population dominated reactivation (Fig. 3a); in rodents, learning-recruited ensembles reactivate during REM sleep, and silencing them specifically during REM sleep impairs consolidation [27, 28]. In the model, the shift toward inhibition landed on the inputs to the reactivated neurons (Fig. 3c); in vivo, REM-specific dendritic calcium spikes prune a subset of newly formed spines while strengthening others [29], and REM sleep is required for experience-dependent elimination of preexisting spines [30]. In the model, hidden-layer activity declined over each sleep episode (Supplementary Fig. S2), as hippocampal firing rates decline across REM episodes in proportion to theta power [31–33]. Competing memories became more distinct (Fig. 2d); in humans, a nap containing REM sleep, but not NREM sleep alone, differentiates competing memories in hippocampal CA2/3/dentate gyrus [34]. Older memories recovered after new learning (Fig. 1c). Similarly, REM sleep after training rescues perceptual learning from interference by a competing task [35] and lowers the cortical excitation/inhibition balance, which stabilizes learning [15]. The model also makes several predictions (Fig. 8b). First, target specificity: synaptic plasticity during REM sleep should concentrate on the inputs to the ensemble that is reactivated, rather than being distributed evenly across ensembles. Second, a depth gradient: because source specificity increased from the first to the last processing stage in four of the five models tested (Fig. 7c), REM-dependent separation of overlapping memories should be strongest at the top of the processing hierarchy (hippocampus and association cortex) and weakest in early sensory cortex. Third, graded recency: when sleep replayed a mixture of all learned tasks, reactivation of each task’s population declined with the age of the memory (Supplementary Fig. S22a), predicting that REM reactivation should decline with memory age.

**Fig. 8:**
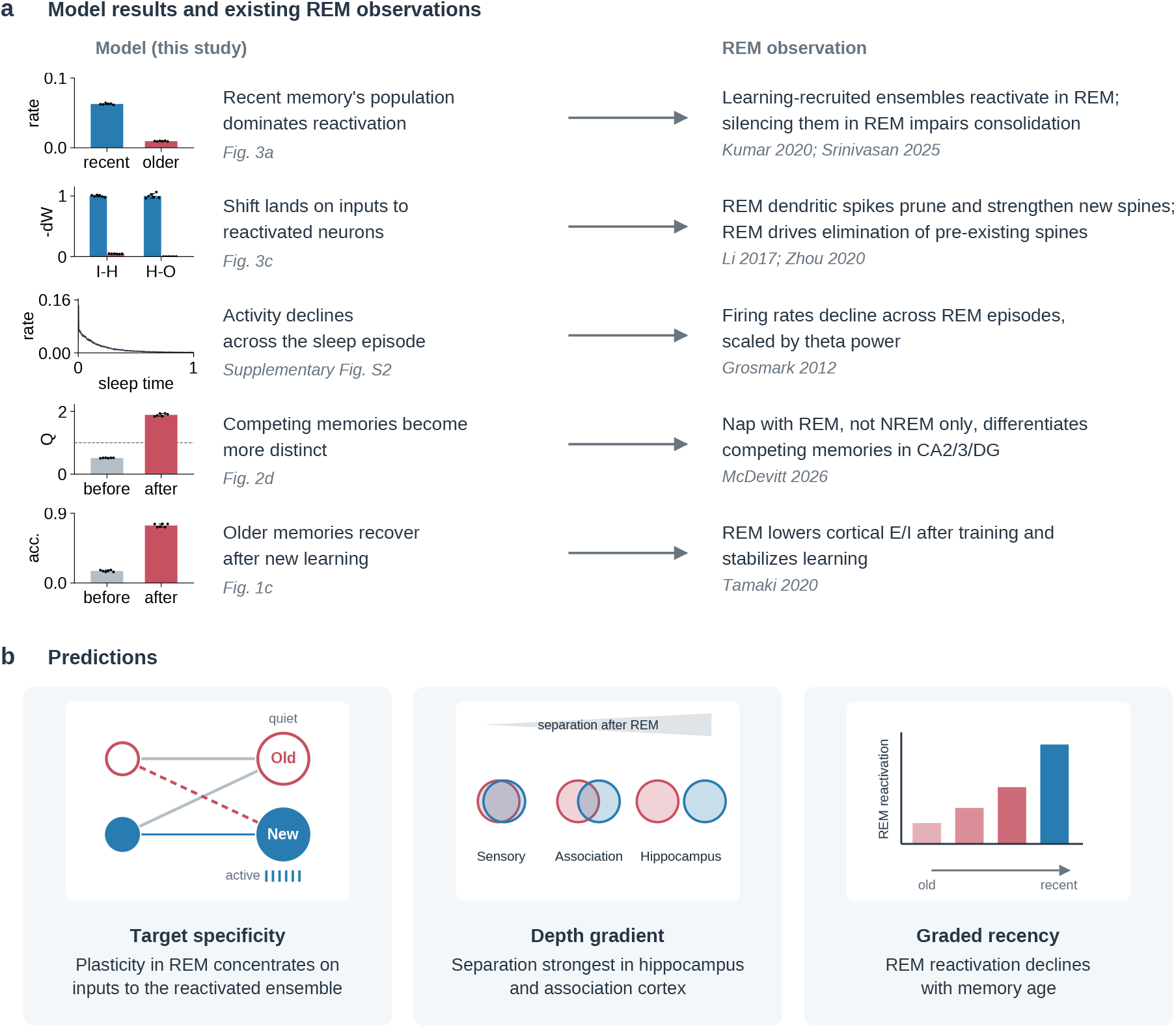
Model results, existing REM-sleep observations and predictions. **(a)** Left: five results of the model, each with a summary plot and the figure it comes from. The most recent task’s population dominates reactivation (firing rate of the recent vs older task populations during sleep; Fig. 3a). The shift toward inhibition lands on the inputs to the reactivated neurons (−Δ*W* from older-task sources onto the recent vs older populations, normalized to the recent value, I → H and H → O; Fig. 3c). Hidden-layer activity declines across the sleep episode (Supplementary Fig. S2). Competing memories become more distinct (intrusion index *Q* before and after sleep; Fig. 2d). Older memories recover after new learning (old-task accuracy before and after sleep; Fig. 1c). Right: corresponding observations of REM sleep. Bar plots: pooled over Sleeps 2–5, mean ± SD over six trials; dots, individual trials. **(b)** Predictions of the model (see text): target specificity, a depth gradient of memory separation, and graded recency of REM reactivation.

### 3.1 REM vs NREM sleep in memory consolidation

Several theories assign REM sleep a role in reducing interference between memories, each through a different proposed process. Crick and Mitchison suggested that dreaming actively unlearns spurious associations [7]; error-correction accounts hold that REM sleep exposes and removes inappropriate links [6, 10]; and the Refinement-and-Rescue hypothesis proposes that REM sleep sparsifies memories, thereby separating similar ones [11]. Despite their differences, these proposals predict the same outcome: overlapping memories become more distinct after REM sleep. Our results identify a single local mechanism that is sufficient to produce this outcome: synaptic plasticity, driven by asymmetric reactivation, that removes cross-talk between memories.

In a classic network model, Hopfield and colleagues let the network settle from random starting states and then weakened the connections between neurons that were active together [36]. This “unlearning” removed spurious memories, but it also weakened real ones, so it helped only in small amounts. In a model of the hippocampus and neocortex, REM sleep replayed older memories and protected them from interference by new learning [37]. In a biophysical model, high acetylcholine during REM sleep increased inhibition, and spike-timing-dependent plasticity then pruned connections and separated the representations of different memories [38]. In our model, reactivation is dominated by the newest memory, and sleep weakens the inputs that this memory took over from older ones. The newest memory itself is protected because its neurons fire together with their own inputs. Other models assign REM sleep a complementary, integrative role: in a two-stage model of sleep, REM-stage reactivation of weak patterns supports selective generalization of memories through synaptic tagging and capture [39]. The two roles need not conflict: by removing indiscriminate overlap between memories, reactivation-targeted pruning may provide the separated representations on which such selective linking can operate.

The dynamics of sleep in our model support assigning this mechanism to REM sleep specifically, rather than to NREM sleep or to sleep in general. NREM sleep consolidation is characterized by large-scale synchronized events – slow oscillations, spindles, and hippocampal sharp-wave ripples – that support coordinated reactivation of recent experience [16, 40]. Sleep in our model lacks these large-scale synchronized events. Activity is driven by unstructured Poisson input. Although neurons representing the most recent task are more active, we found no evidence of precise replay of learned patterns during sleep. Instead, the dominant effect of sleep on network dynamics is a net decrease in spiking activity and a systematic shift of the synaptic E/I balance toward inhibition. This combination of desynchronized, non-replay-like dynamics and a shift toward inhibition matches what is known about REM sleep: it lacks the slow-oscillation-locked synchrony of NREM sleep [40], and its neuromodulatory and synaptic environment favors depression and pruning [29, 30, 41].

### 3.2 REM sleep and synaptic plasticity

A postsynaptic spike is necessary for any synaptic modification in our model; given one, the sign of the change depends on whether the presynaptic partner was simultaneously active or silent. Coincident pre- and postsynaptic firing potentiates the synapse, a conventional Hebbian term. Postsynaptic firing in the absence of presynaptic input depresses excitatory synapses and potentiates inhibitory synapses. Plasticity at excitatory synapses that were not presynaptically active during induction (heterosynaptic plasticity [42]) is driven by postsynaptic activity alone, and is precisely the event our depression term implements. Experiments in cortical slices have shown that postsynaptic spiking or depolarization without paired presynaptic input produces weight-dependent changes at non-activated synapses, with net depression at strong synapses [42, 43]. This depression counteracts the positive feedback intrinsic to Hebbian potentiation and prevents runaway synaptic dynamics [44]; our model contains both terms in exactly that configuration. The input specificity of synaptic plasticity is furthermore only partial at individual synapses, so that postsynaptic activity modifies synapses that were themselves silent [21]. Heterosynaptic plasticity also operates in vivo and shapes the balance of a neuron’s inputs: in mouse visual cortex, Hebbian potentiation of activated spines is accompanied by heterosynaptic weakening of neighbouring spines [45], and in auditory cortex heterosynaptic depression of the strongest unpaired inputs normalizes synaptic strengths and sets the neuron’s excitation/inhibition balance [46]. Early work in amygdala neurons found that heterosynaptic changes nearly conserve the total synaptic weight [47]; more recent data from cortex indicate that they are weight-dependent and normalize the distribution of synaptic strengths, acting mainly on the strongest inputs, rather than strictly conserving their sum [42, 46].

For inhibitory synapses the experimental picture is inconsistent. At GABAergic synapses onto thalamocortical relay cells, iLTP is induced by postsynaptic burst firing alone and is independent of pre–post spike timing: pre-then-post, post-then-pre, and uncorrelated pairings produce equivalent potentiation [48]. Inhibitory inputs onto layer 5 neocortical pyramidal neurons show the same character, the direction of change being gated by postsynaptic membrane potential and firing mode rather than by presynaptic timing, with hyperpolarized bursting yielding iLTP and depolarized tonic firing yielding iLTD [49]; postsynaptic depolarization or brief spike bursts alone are likewise sufficient to potentiate perisomatic GABAergic synapses in this population [50]. At these synapses, inhibitory plasticity is gated by the post-synaptic state, as in BCM-type plasticity, rather than by spike timing as in classical STDP. Inhibitory plasticity at parvalbumin and somatostatin synapses onto hippocampal CA1 pyramidal neurons, by contrast, is strictly coincidence-dependent: presynaptic input alone and postsynaptic spikes alone are each insufficient, and the timing dependence is Hebbian [51]. Inhibitory plasticity rules thus appear not to be conserved across brain regions and interneuron subtypes. This uncertainty does not affect our conclusions: in the Dale’s-law model, excitatory plasticity alone was sufficient to restore older memories, and its weight changes showed the same reactivation-targeted pruning.

The central claim of our model is that sleep-driven plasticity is concentrated on the synapses onto the neurons of the most recently trained task, which are active during reactivation, and most strongly on those from older-memory neurons, which are silent (Figs. 3c, 4). In vivo two-photon spine imaging provides a structural analog of this selectivity. Li et al. [29] showed that REM sleep selectively prunes a subset of newly formed dendritic spines in mouse motor cortex while sparing and strengthening others, gated by REM-specific dendritic calcium spikes. Zhou et al. [30] extended this finding by showing that REM sleep specifically – not slow-wave sleep of comparable duration – is required for experience-dependent spine elimination following monocular deprivation or fear conditioning, and that REM sleep deprivation blocks both the structural elimination and the associated reduction in evoked neuronal activity. Neither study reports uniform, network-wide synaptic weakening; both instead describe activity- and experience-dependent selectivity, consistent with the block-structured, task-selective pattern of depression our model predicts rather than a purely diffuse downscaling process.

In our model, a synapse is depressed when its postsynaptic neuron fires while its presynaptic neuron is silent. Because units of the most recent task are the most active during sleep and older-task units are mostly quiet, these events concentrate on the synapses onto the recent task’s neurons. This leads to a testable prediction: how much the synapses between two overlapping memory representations change during REM sleep should be predictable from how often each population is reactivated during REM sleep. This prediction is directly testable using short-latency functional-connectivity methods for tracking synaptic change following sleep manipulations in vivo, and links the model to REM sleep manipulation experiments on memory consolidation [27, 28], representational differentiation [34] and recovery from interference [35].

The reactivation asymmetry itself is well supported: sleep preferentially reactivates recently acquired experience, with reactivation strongest shortly after learning and decaying over subsequent sleep [16, 52–54], and ensembles recruited during recent learning are also reactivated during REM sleep [27, 28, 55]. REM-specific neuromodulation may further bias processing toward recent memories: elevated acetylcholine suppresses recurrent excitatory transmission and reduces interference from stored associations [56], and REM theta activity has been proposed to process novel and familiar information differently, through phase-dependent potentiation and depotentiation [57].

At the representational level, our model predicts that between-class correlation should decrease substantially more than within-class correlation after sleep, and that overlapping neuronal populations should become progressively more disjoint. Direct recordings of hippocampal or cortical population overlap across a full human sleep cycle remain technically difficult, but related evidence from waking pattern-separation studies is consistent with our model predictions. Zotow, Bisby and Burgess [58] provide behavioral evidence that overlapping, but not unrelated, events are actively separated during human episodic encoding. Using high-resolution fMRI of human CA3/dentate gyrus, Guo et al. [59] show that resolution of behavioral memory interference is temporally coupled to an abrupt decorrelation of hippocampal activity patterns for the two competing memories, with greater initial pattern overlap predicting larger subsequent decorrelation – closely paralleling the relationship we observe between pre-sleep population overlap and post-sleep Jaccard index. These studies do not show that decorrelation happens during REM sleep. They do show that the outcome our model reaches, decorrelation of overlapping representations that reduces interference, occurs in biological memory, which makes REM sleep a candidate offline mechanism for it.

### 3.3 Model design and limitations

Our choice to model the REM-sleep-like phase with a recurrent neural network, rather than a biophysically detailed spiking model, was motivated by the nature of the memories being consolidated. Biophysically detailed spiking models are well suited to represent replay of simple, well-defined sequences, and are a natural framework for NREM sleep, where consolidation is thought to involve compressed replay of specific, previously experienced trajectories [60–63]. REM sleep, by contrast, is hypothesized to act on complex, high-dimensional memories – in our case, entire learned classification tasks – for which comparably detailed biophysical implementations are not yet tractable at scale. RNNs provide a framework capable of storing and reactivating memories of this complexity, but RNNs are conventionally trained with backpropagation through time, which is not biologically plausible. We therefore trained our RNN with EP, which preserves a fully local, biologically plausible learning rule while still supporting the more complex memory representations that REM-sleep-like processing requires.

This choice also limits the model. MRNN-EP is a strongly simplified abstraction of thalamocortical and hippocampal circuitry; it lacks the spiking sharp-wave-ripple dynamics of NREM sleep, and SRC as implemented here models a REM-sleep-like phase only, without a preceding NREM-sleep-like integration phase. Consequently, our model cannot speak to how NREM-sleep-driven integration and REM-sleep-driven separation are jointly balanced in the biological sleep cycle, and we cannot test whether the crosstalk sleep removes is crosstalk that a prior NREM-sleep-like phase would have introduced [5, 6].

This work is a tractable first step toward that larger question. Spiking implementations of EP are available [64–67]. Extending them with an explicit NREM-sleep phase – hippocampal sharp-wave-ripple-driven replay – followed by a REM-sleep-like desynchronized, LTD-biased reactivation phase would allow direct tests of whether interleaving integration and separation phases is required for interference-free consolidation, as opposed to REM-sleep-like separation alone. It would also allow the synaptic and representational predictions from this model to be compared directly against functional connectivity changes measured in vivo under chemogenetic and optogenetic manipulation of REM and NREM sleep.

### 3.4 Conclusions

Our study proposes reactivation-targeted pruning as a synaptic mechanism for the role of REM sleep in reducing memory interference. Its principle is simple: reactivation determines where plasticity acts, and local plasticity determines what it does there. Because the newest, not yet consolidated memory dominates reactivation, the connections through which it has captured the inputs of older memories are weakened, while the connections onto older memories are left intact. The newest memory’s own inputs, which fire together with its neurons, are protected by Hebbian potentiation. The mechanism emerged in every network we analyzed, regardless of architecture, waking learning rule or Dale’s-law constraints, suggesting that it is a general consequence of local plasticity acting on biased reactivation rather than a property of a particular circuit.

More broadly, our results show how artificial neural networks that face a problem the brain must also solve, learning new memories without overwriting old ones, can be used to understand how the brain solves it. The insight can then return to artificial intelligence: the principle identified here, that offline reactivation should prune the inputs a new memory has taken over from older ones, suggests how sleep-like phases could be designed to help AI models adapt to change and learn continually. Moving in this way between artificial and biological networks, with models from AI generating hypotheses about the brain and the mechanisms they reveal informing new AI, may benefit both fields.

## 4 Methods

### 4.1 Equilibrium Propagation (EP)

Equilibrium Propagation (EP) is a biologically plausible learning algorithm that trains networks by minimizing an energy function through local, contrastive updates [17, 22]. This approach specifically leverages *convergent recurrent neural networks*, whose internal dynamics are designed to settle into a stable, fixed-point state. The energy function *E* defines the network’s internal configuration: neurons adjust their activations to reach states that locally minimize *E*. The state update dynamics follow the negative gradient of the energy function with respect to the neuron activations.

The convergent property is crucial for EP. It ensures that when a static input is provided, the network’s recurrent dynamics evolve over time until the neuron activations stop changing and reach a stable equilibrium. This convergence to an energy minimum underlies how EP generates learning signals using only local information.

In continuous time, the neuron states follow gradient flow:

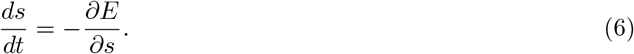

This means the neuron activations evolve continuously to reduce the total energy. The discretized version of this flow aligns with the update formulation described by [67] for energy-based recurrent dynamics, providing a clear link to modern formulations of equilibrium models.

EP training consists of two distinct phases. In the *free phase*, the network settles naturally into a local energy minimum given an input, without any external target. In the *weakly clamped phase*, a small nudging term gently pushes the output units towards the desired target, perturbing the equilibrium slightly. By contrasting the network states between these two conditions, synaptic weights are updated through purely local Hebbian-like rules.

#### Energy and Cost Relationship

To incorporate supervised learning, the total objective combines the internal energy *E* and an external cost function *C* that measures output error (e.g., squared difference from the target). During the weakly clamped phase, the dynamics minimize:

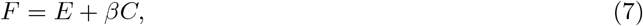

where *β* is a small nudging parameter. This coupling ensures that the output units are biased to reduce the cost, while the system still performs local energy minimization, producing a contrastive signal that approximates backpropagation but with local updates.

#### Energy Function

The network’s state dynamics are governed by minimizing an energy function *E*(*s, x, w*), where *s* denotes neuron activations, *x* is the input, and *w* are the synaptic weights. For example, a Hopfield-like recurrent network can be described by:

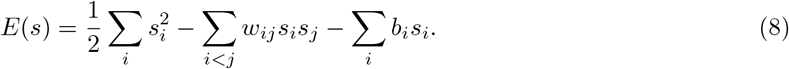

This structure encourages the network to settle into stable configurations that minimize *E*.

#### Free Phase

In the free phase, the network relaxes to a local minimum of the energy function given an input *x*. The continuous-time dynamics (Eq. 6) are discretized using an Euler method. For a simple multilayer setting with one hidden layer:

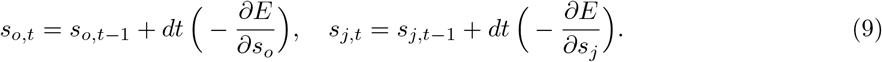

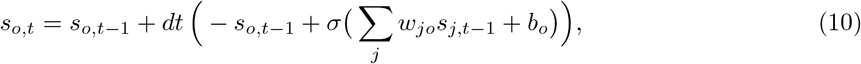

These expand to:

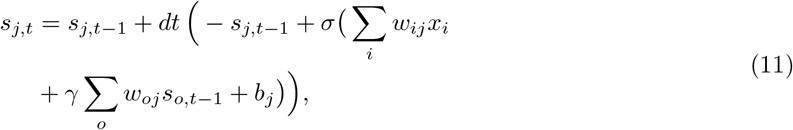

where *σ*(·) is an activation function, *γ* is a feedback scaling parameter, and *dt* is the integration step size.

#### Weakly Clamped Phase

In the weakly clamped phase, the output neurons receive a small nudging force that biases them towards the target *y* by adding the cost gradient:

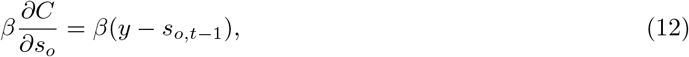

assuming *C* is mean squared error. The output dynamics become:

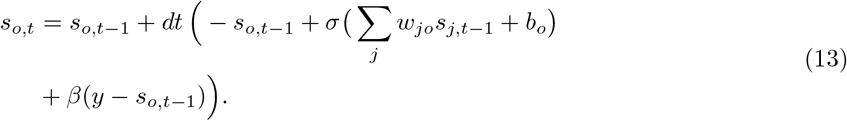

The hidden states evolve as in the free phase (Eq. 11).

#### Weight Update Rule

Finally, the synaptic weights are updated by contrasting neuron correlations between the free and weakly clamped phases:

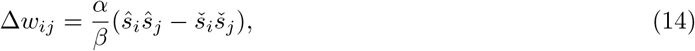

where *ŝ* denotes activations in the weakly clamped state, *š* those in the free state, and *α* is the learning rate.

This local contrastive update approximates the true gradient of the cost with respect to the weights, demonstrating how EP can perform supervised learning using only local, biologically plausible signals [17].

### 4.2 Predictive Learning Rule

Luczak et al. [23] explored a biologically inspired variant of Hebbian learning for Equilibrium Propagation (EP) and similar local training algorithms. The update rule they introduced is called the *predictive learning rule*. This formulation arises naturally if one assumes that each neuron strives to maximize its own metabolic energy balance by predicting its future activity and adjusting synaptic strengths accordingly.

The predictive learning rule is formally expressed as:

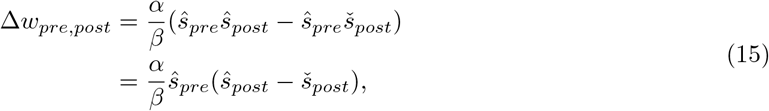

where:

- *w*_*pre,post*_ denotes the synaptic weight connecting the *presynaptic* neuron (*pre*) to the *postsynaptic* neuron (*post*).
- *ŝ*_*pre*_ is the activity of the presynaptic neuron during the weakly clamped phase.
- *ŝ*_*post*_ is the actual activity of the postsynaptic neuron in the weakly clamped phase.
- *š*_*post*_ is the actual activity of the postsynaptic neuron in the free phase.

In the original version, Luczak et al. used the *predicted postsynaptic activity*, denoted by 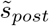, in place of the free-phase activity (*š*_*post*_). Here, the “predicted postsynaptic activity” means the neuron’s internal estimate of what its activation will be in the near future, based on its internal state and local signals. This predictive signal represents the neuron’s capacity to anticipate its own next equilibrium value, which is thought to align with observations of predictive coding in biological neurons.

Their results showed that the correlation between the predicted 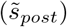 and actual (*š*_*post*_) postsynaptic activity during the free phase was nearly perfect (*R* = 1 ± 0.0001 SD), demonstrating that the explicit prediction provides negligible additional signal under these conditions.

The rule builds on predictive neuronal adaptation [68]; following [24], we use a simplified version in which the predicted signal is replaced by the actual free-phase activation.

Therefore, in this study, which focuses on isolating the effect of sleep-like replay in continual learning, we adopt the simpler form without the explicit prediction term.

### 4.3 Sleep Replay Consolidation (SRC)

Tadros et al. [25] introduced Sleep Replay Consolidation (SRC), a sleep-like, unsupervised replay mechanism originally designed for feedforward neural networks (FFs) trained with backpropagation. In their approach, the trained artificial neural network (ANN) is converted to a spiking neural network (SNN) with the same architecture. During the sleep phase, the SNN’s activity is driven by randomly distributed Poisson spiking input. Each input element (e.g., a pixel) spikes with a probability proportional to its mean intensity across the training data, so pixels that are consistently bright spike more often.

During the sleep phase, the learning rule used is a local Hebbian plasticity mechanism similar to spike-timing-dependent plasticity (STDP): synaptic weights are increased when presynaptic spikes precede postsynaptic spikes and decreased when postsynaptic spikes occur without preceding presynaptic input. This local STDP update replaces the standard EP-based predictive update rule during replay because the network operates in the spiking domain. After the sleep replay phase, the updated SNN weights are mapped back to the ANN used for task learning.

We used the recurrent implementation of SRC [26], which applies SRC to both the output and hidden recurrent layers. Feedback connections (transposed hidden-to-output weights) govern the membrane potential dynamics in a manner consistent with the convergent EP updates used during wake learning. When the membrane potential crosses a threshold, the neuron spikes, and weights are adjusted using the local STDP rule.

In our training procedure, these phases are interleaved: the network is first trained on tasks using the EP-based learning rule in the awake phase, and then SRC with the STDP rule is applied during a sleep phase after each new task. This combined approach mirrors the biological interleaving of awake learning and sleep-dependent consolidation observed in the brain [25].

Prior work has shown that SRC-equipped models increase representational sparsity, reduce class overlap, and improve robustness to noise and adversarial perturbations [25, 26, 69–71]. Here we use SRC to identify the synaptic changes through which sleep produces these effects.

### 4.4 Model and Dataset Specification

We used MNIST [72] for the main analyses and CIFAR-10 [73] to test the mechanism on natural images. The MNIST dataset comprises 60,000 28 × 28 grayscale images of handwritten digits (10 classes).

Our main architecture is a recurrent neural network trained using Equilibrium Propagation (MRNN-EP). For MNIST, we use 784 input neurons, 1024 hidden neurons, and 10 output neurons (784–1024–10). Hidden layers use ReLU activations; the output uses a hard sigmoid as in [74].

For the generality analysis, we compared MRNN-EP with a standard RNN with recurrent connections only in the hidden layer, trained with Backpropagation Through Time (RNN-BPTT). Supplementary Fig. S23 illustrates the architectural differences between the two networks.

#### Computational cost

On an NVIDIA GeForce RTX 3080 Ti, training MRNN-EP on five tasks takes about 4 minutes without SRC and 7 minutes with SRC. The five sleep phases therefore add about 3 minutes, comparable to the cost of training a single task, as in previous work [25].

#### Phase Notation

Throughout the paper — especially in the analysis sections — we use T# to refer to the #-th training phase, during which the model is trained on a new task, and S# to denote the sleep-like replay phase that follows it. This notation is used consistently in our confusion matrix visualizations and other analysis figures to indicate the position in the training sequence, rather than a fixed task identity.

##### CIFAR-10

To test the mechanism on natural images, we trained MRNN-EP on the CIFAR-10 dataset [73], which comprises 60,000 color images (32 × 32 pixels) across 10 object categories, split into five two-class tasks as for MNIST. Because of the higher dimensionality of these images, we extracted features using a pretrained convolutional network composed of VGG-style blocks [75], following previous studies [25, 26]; its weights were held fixed during training and sleep.

###### Algorithm 1

SRC algorithm

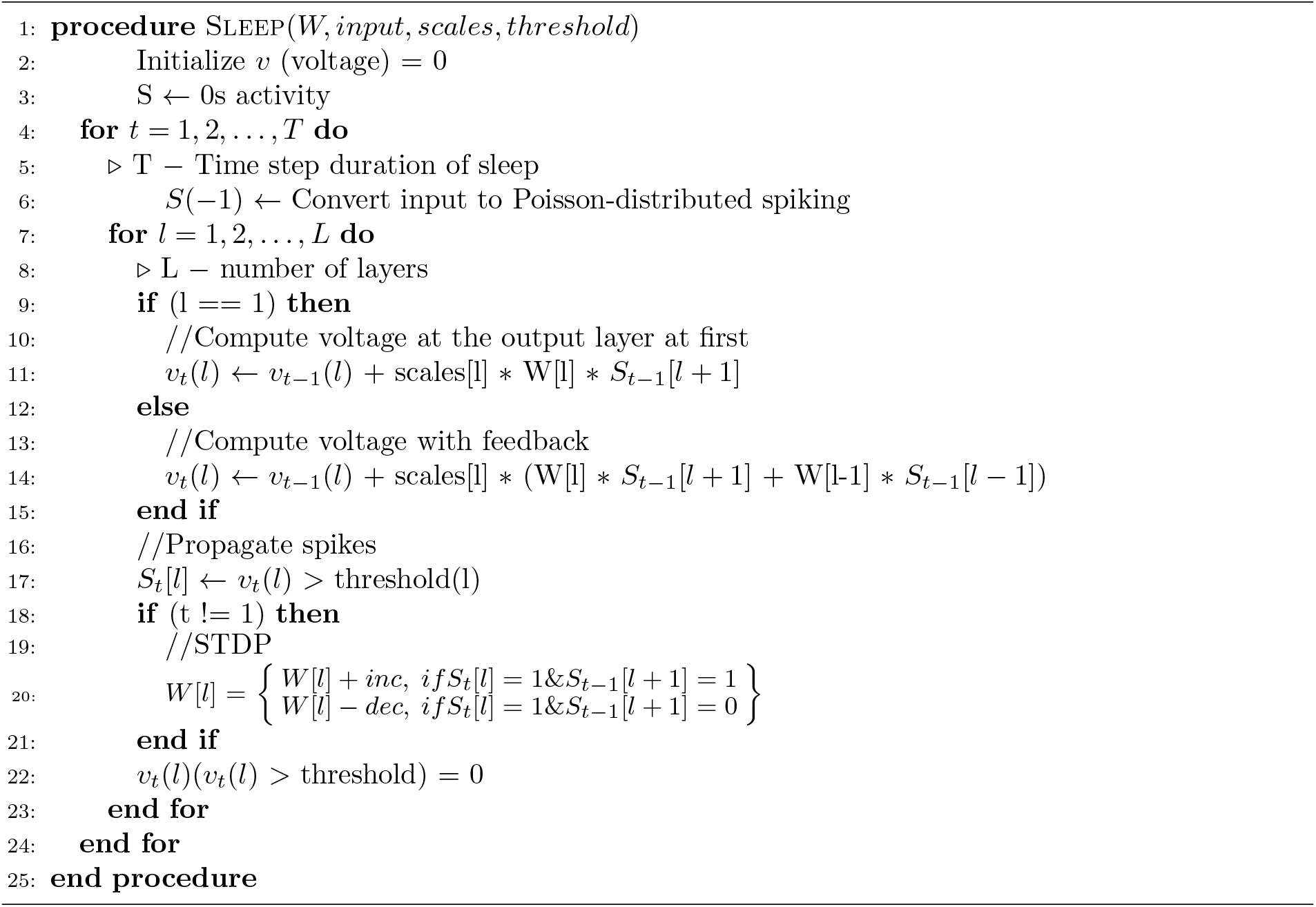

The feature extractor architecture includes the following components:

- Block 1: Two convolutional layers with 64 kernels (3 × 3), each followed by batch normalization [76] and ReLU activation.
- Block 2: Two convolutional layers with 128 kernels (3 × 3), each followed by batch normalization and ReLU activation.
- Block 3: Two convolutional layers with 256 kernels (3 × 3), each followed by batch normalization and ReLU activation.
- Block 4: One convolutional layer with 512 kernels (3 × 3), followed by batch normalization and ReLU activation.

The feature maps at the output of this network (512 × 4 × 4, flattened to 8192 values) were the input to MRNN-EP (8192–1024–10). The pretrained model in this experiment is from GitHub (VGG-13 with batch normalization at https://github.com/chenyaofo/pytorch-cifar-models/tree/master?tab=readme-ov-file). MRNN-EP was trained for 5 epochs per task, with 125 free-phase and 15 clamped-phase time steps, and each sleep episode replayed only the most recently trained task, as for MNIST. Sleep hyperparameters were optimized with the genetic algorithm described below, starting from the values of [26].

RNN-BPTT was trained using standard stochastic gradient descent (SGD) with a fixed learning rate of 0.01. The hyperparameters for our proposed model (MRNN-EP) are detailed in Supplementary Table S2. All models were trained using a batch size of 256. Training was performed for 3 epochs per task on MNIST and for 5 epochs per task on CIFAR-10.

### 4.5 Incremental Task Evaluation

To test the efficacy of SRC in mitigating catastrophic forgetting, we created five incremental tasks from each dataset. Each task included data for two distinct labels (e.g., task 1 contained labels 0 and 1, task 2 contained labels 2 and 3, etc.) for all the datasets. For MNIST, 10% of the training data was held out as a validation split and used for evaluation; for CIFAR-10, the official test set was used for evaluation. Each model was trained six times (seeds 0–5) on the same task order, and results are reported as mean ± SD over these six trials. On the official MNIST test set, which was used neither for training nor for hyperparameter selection, the final accuracy of the main model was 71.9 ± 2.9% (70.6 ± 3.2% on the validation split).

### 4.6 SRC Hyperparameter Optimization

The hyperparameters for SRC were optimized using a population-based Genetic Algorithm (GA) implemented in Python via the geneticalgorithm package [77]. The GA was configured with a population size of 100, uniform crossover with a probability of 0.75, and a mutation probability of 0.1 per parameter. Elitism was applied by carrying over the top 1% of individuals unchanged to the next generation, while 20% of the population was selected as parents for reproduction using fitness-proportionate (roulette-wheel) selection.

For MNIST, the fitness of each candidate solution was evaluated on a held-out validation split comprising 10% of the training data, and defined as the average classification accuracy across all tasks on this split. For CIFAR-10, sleep hyperparameters were initialized from ref. [26] and refined by a short GA; accuracy was insensitive to this refinement (82.7% with the initial parameters vs. 83.8% after refinement, for the same random seed).

### 4.7 Minimal Spiking Network

The minimal network (Fig. 6) had 10 input, 10 hidden and 10 output units, one per digit class. For each class *c*, we selected the 75 pixels whose intensities in that class’s average image most exceeded the corresponding average intensities in the other classes. At each time step, input unit *c* fired with a probability equal to the mean intensity of these pixels divided by 1.906, following the Poisson encoding used during sleep. Class-average images were computed from the training images for sleep and from the test images for testing. Input-to-hidden weights were 0.003 per synapse from each input unit onto its own hidden unit, 1.5 × 0.003 from each input unit of classes 0–7 onto hidden units 8 and 9, and 0 otherwise. Hidden-to-output weights were the mean weights between each class’s top-10% most-active set (as above) and each output unit of the trained MRNN-EP network (trial 0) immediately before Sleep 5. The drive of each input or hidden unit was multiplied by the size of the population it represents, and output-to-hidden feedback used the hidden-to-output weights, as in the full model. Sleep followed the spiking dynamics and plasticity rule of the full model (Algorithm 1) with that network’s Sleep-5 parameters (387 time steps, leak, layer scaling, inc and dec), driven by the Task-5 training mean image; firing thresholds were set to 10 (hidden) and 30 (output) times the model’s values. In the depression-only condition inc was set to 0. Testing used the same dynamics without plasticity: each class-average test image was presented for 387 time steps, and the output unit with the most spikes was taken as the network’s answer. Each condition was run 20 times with different random seeds; post-sleep weights were averaged over the 20 sleep runs. Per-class accuracy of the full network was obtained from its standard free-phase inference on the validation images of all five tasks.

### 4.8 Block-Matrix Analysis of Sleep-Induced Weight Changes

#### Task- and class-specific neuron/pixel groups

For the block-matrix heatmaps, we first defined fixed neuron and pixel groups for each of the ten digit classes, computed once from that class’s own task at its last waking training epoch (using only that task’s own validation data) and then reused across all subsequent sleep stages to track the same neuron identities over time. Per-class hidden-layer groups were the class-specific top-10% most-active sets: the 10% of hidden neurons with the highest mean activation on that class’s validation data, computed once at the end of that task’s training (its last waking epoch) and held fixed thereafter. Per-class input-layer groups were defined as the top 75 most discriminative input pixels for that class. This is distinct from the per-phase-epoch population analysis used for the Jaccard overlap measure, in which groups are instead redefined independently at every training and sleep epoch.

#### Block-matrix heatmaps

For the hidden-to-output (H → O) pathway, we computed Δ*W* = *W*_post_ − *W*_pre_ for every synapse connecting each of the ten hidden-layer class groups to each of the ten output units, then averaged Δ*W* within each (hidden group, output unit) pair to obtain a 10 × 10 block matrix per sleep stage. The input-to-hidden (I → H) pathway was treated analogously, with input pixel groups in place of hidden-layer groups. This was repeated independently for each of the five sleep stages, yielding the multi-panel heatmaps in Fig. 3c. Waking weight changes (Fig. 3b) were computed in the same way, with Δ*W* the weights at the end of a task’s training minus those at its start (for Task 1, the initial weights).

### 4.9 Specificity Indices, Intrusion Index and Manipulations of Sleep Weight Changes

#### Task populations

For each task, the hidden population was its fixed hidden-neuron group, the union of that task’s two class-specific top-10% most-active sets (see above), the input group was the union of the top-75 discriminative pixels of its two classes, and the output group consisted of its two output units. For sleep episode *k* (*k* = 2, …, 5), task *k* is the *recent* task and tasks 1, …, *k* − 1 are the *older* tasks.

#### Target and source specificity

For each pathway and sleep episode *k*, let *s* = −(*W*_post_ − *W*_pre_) be the per-synapse shift toward inhibition. Units shared between the recent group and the union of the older groups were removed from both. Target specificity compares the mean shift onto the recent versus the older postsynaptic groups, over all presynaptic units of tasks 1, …, *k*: TS = (*s*_recent_ − *s*_older_)*/*(|*s*_recent_|+ |*s*_older_|). Source specificity compares, onto the recent postsynaptic group, the mean shift from older versus own presynaptic groups: SS = (*s*_old src_ − |*s*_own src_|)*/*(|*s*_old src_| + |*s*_own src_|). Means are taken over all synapses of a block. Both indices range from −1 to 1; the value for a trial is the mean over Sleeps 2–5. For the Dale’s-law models, effective weights |*W*| × sign were used. Statistics: exact one-sided (TS *>* 0) and two-sided (SS ≠ 0) one-sample Wilcoxon tests over the six trials.

#### Intrusion index

For sleep episode *k* and older task *j < k, Q* is the mean hidden activity of task *j*’s own population divided by the mean hidden activity of task *k*’s population, both measured on task *j*’s validation images, at the end of task *k*’s training (before sleep) and after sleep *k. Q* was averaged over older tasks within each trial and sleep, and pooled over sleeps 2–5. A variant with the units shared by the two populations removed from both gave the same result.

#### Manipulations of sleep weight changes

All manipulations start from the pre-sleep weights of sleep *k* and apply a modified version of that sleep’s own weight change Δ*W* = *W*_post_ − *W*_pre_ in both the I → H and H → O pathways (feedback uses the transposed H → O weights, so each manipulation acts identically on the feedforward and feedback paths); biases are unchanged by sleep. *Target specificity removed* : for every presynaptic unit, its mean change onto the recent postsynaptic group and its mean change onto all other postsynaptic units were both replaced by their size-weighted pooled mean, keeping each synapse’s deviation from its group mean; this preserves every presynaptic unit’s total change. *Source specificity removed* : for every postsynaptic unit, its mean change from own-task inputs and from older-task inputs were both replaced by their pooled mean, keeping each synapse’s deviation; this preserves every postsynaptic unit’s total change. *Uniform*: each pathway’s Δ*W* replaced by its mean over all synapses of that pathway. Readouts were old-task accuracy (mean over tasks 1, …, *k* − 1), recent-task accuracy and *Q*, pooled over Sleeps 2–5; conditions were compared with real sleep by exact paired Wilcoxon tests over the six trials.

### 4.10 Dale’s-Law Models

In the Dale’s-law variant of MRNN-EP, every synapse of the input-to-hidden and hidden-to-output weight matrices was assigned a fixed excitatory or inhibitory sign at initialization (50% of each), with effective weight *W*_eff_ = |*W*_raw_| × sign at every forward pass and after every update. During sleep, inhibitory synapses used a sign-swapped magnitude rule relative to excitatory synapses (Supplementary Section S2). An additional sleep episode was applied before Task 1 training. In the excitatory-only and inhibitory-only variants, only excitatory or only inhibitory synapses, respectively, were modified during sleep, while the other synapses were held at their pre-sleep values. Sleep hyperparameters were optimized by genetic algorithm separately for each of the three models. Because these runs logged accuracy only after the final sleep, accuracies at intermediate stages were recomputed from the saved network states; the recomputed final accuracies matched the logged values for every task and trial.

## Acknowledgements

This study is supported by NSF (grant 2223839) and NIH (grants R01MH125557, R01NS132913 and R01AG099626).

## Author contributions

M.B. conceived and designed the study. Y.K. and M.B. implemented the models and performed the experiments. Y.K., J.E.D. and M.B. analyzed and interpreted the results and wrote and revised the manuscript.

## Data availability

The MNIST and CIFAR-10 datasets analyzed in this study are publicly available [72, 73]. All simulated data can be reproduced with the code described in the Code availability statement.

## Code availability

The code used in this study, including the network and sleep-replay simulations, the genetic-algorithm optimization of sleep parameters and the analysis scripts, will be made publicly available on GitHub and archived on Zenodo with a DOI upon publication. During peer review, it is provided to editors and reviewers as a supplementary software file.

