## Supplementary Information for "Firing Without Wiring: How REM-like Sleep Reactivation Rebalances Excitation and Inhibition to Orthogonalize Memories"

This Supplementary Information provides six additions to the main text. **Supplementary Section S1** reports additional representational, dynamical and synaptic analyses of the main MRNN-EP model: within- and between-class correlations, population overlap and exclusive coding, network activity and weight-change distributions during sleep, the full set of manipulations of sleep's weight changes that underlies main Fig. 4, and a comparison of the analytical model of main Fig. 5 with the full network. **Supplementary Section S2** reports the models with Dale's law imposed at the level of individual synapses, including the ablations in which only excitatory or only inhibitory synapses were plastic during sleep. **Supplementary Section S3** reports the recurrent network trained with backpropagation through time (BPTT), **Supplementary Section S4** MRNN-EP trained on CIFAR-10, **Supplementary Section S5** the feedforward networks consolidated with Sleep Replay Consolidation (SRC) [1] on MNIST and CIFAR-10, and **Supplementary Section S6** an MRNN-EP model in which each sleep episode replays a mixture of all tasks learned so far; **Supplementary Section S7** gives the network architectures and the MRNN-EP hyperparameters. For every model in Sections S2–S6, two figures reproduce the analyses of main Figs. 2 and 3 with identical definitions: (i) the class-level correlation matrices before and after the final sleep, the correlation ratio $R$, the population overlap (mean pairwise Jaccard index) and the intrusion index $Q$; and (ii) the reactivation of task populations during sleep and the block-averaged weight changes during training and during sleep.

#### S1 Additional analyses of the main MRNN-EP model

##### S1.1 Correlations, population overlap and exclusive coding

Supplementary Figure S1 expands main Fig. 2. Before the final sleep, the off-diagonal entries of the pairwise overlap matrix between class populations are uniformly high, indicating that all class populations share a large fraction of neurons; after sleep they drop towards 0.1–0.2, while the diagonal (each class vs. itself, identically 1 by construction) is unchanged (panel a). The fraction of each class's population that is not shared with any other class – the exclusive fraction – rises from $\sim 10\%$ at the end of training to $\sim 25\%$ after sleep on average (panel b). Class "1" is an exception: it maintains a high exclusive fraction even after each new task training, likely because digit 1 has the sparsest raw pixel representation of any MNIST digit and the lowest mean pixel-intensity correlation with every other digit. Finally, the correlation ratio $R$ of main Fig. 2b combines the mean within-class (diagonal) and between-class (off-diagonal) correlations of hidden activations; panel c shows the two components separately at every phase. Sleep reduces both, but the reduction is substantially larger for the between-class correlation, so that the representational gap between digit classes expands after every sleep episode.

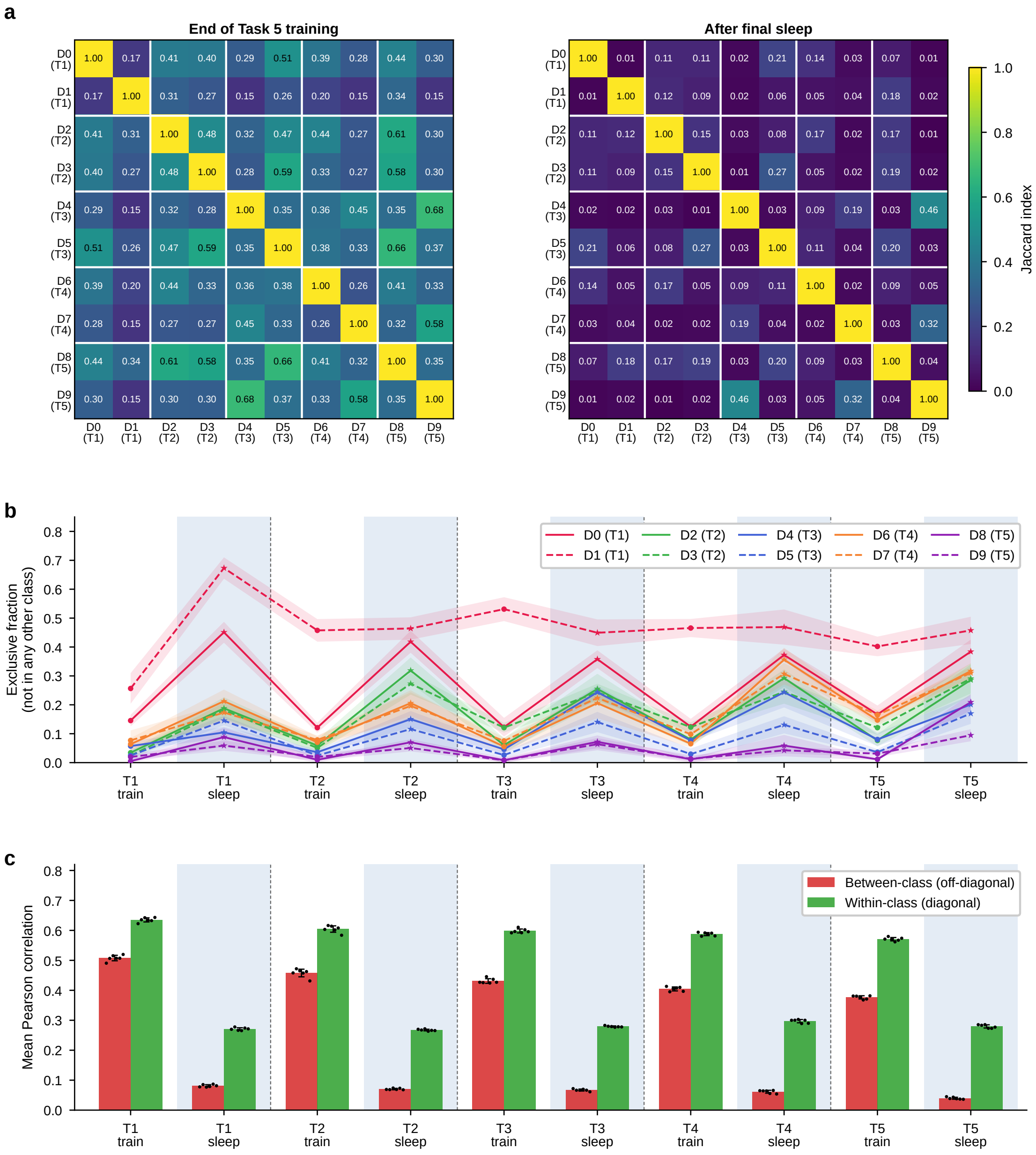


Figure S1: **Sleep carves out task-dedicated neuron populations and decorrelates classes.** **(a)** Pairwise population overlap at the end of Task 5 training (left) and after the final sleep (right): Jaccard index between the top-10% neuron populations of each class pair, mean over six trials (the mean of the off-diagonal entries across all phases is shown in main Fig. 2c); white lines separate tasks. **(b)** Per-class fraction of top-10% neurons that belong exclusively to that class's population, mean ± SD over six trials; circles, end of training; stars, after sleep. **(c)** Mean Pearson correlation of hidden activations for between-class (off-diagonal, red) and within-class (diagonal, green) pairs at the end of every training and sleep phase, mean ± SD over six trials; dots, individual trials; blue shading marks sleep phases.

#### S1.2 Network activity and weight changes during sleep

While the noisy input to the network remains constant during sleep, activity across the hidden and output layers progressively decreases over the course of the episode (Supplementary Fig. S2). The resulting weight changes are shifted in the negative direction: the large majority of synapses in both pathways decrease in strength – decreasing the strength of excitatory synapses and increasing the strength of inhibitory synapses – while only a small fraction is potentiated (Supplementary Fig. S3).

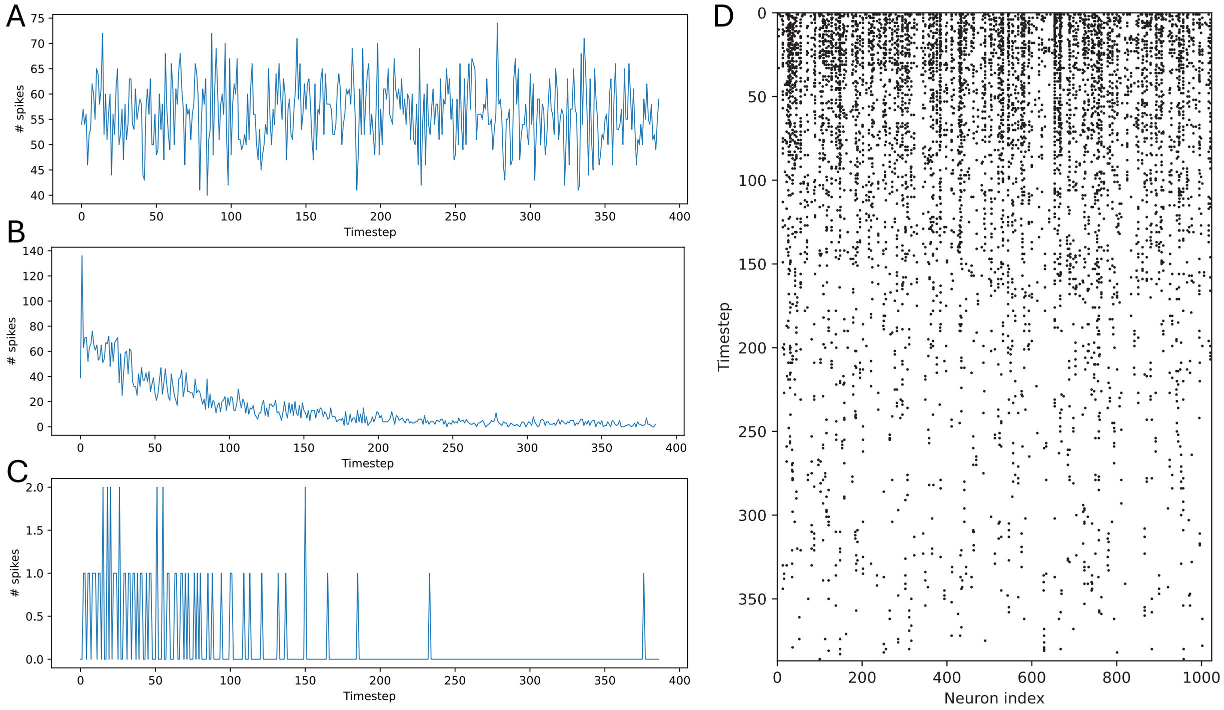


Figure S2: **Firing-rate dynamics over the final sleep episode** (trial 0, sleep after the last task). Number of spikes per timestep in the **(a)** input, **(b)** hidden and **(c)** output layer. **(d)** Raster plot of spiking activity in the hidden layer.

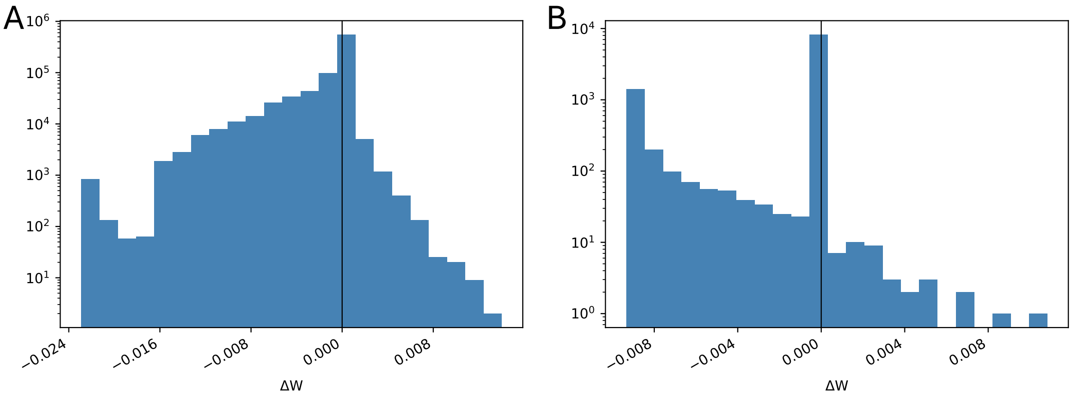


Figure S3: **Distribution of sleep-induced weight changes** (post- minus pre-sleep, final sleep episode, trial 0; log-scale counts). **(a)** Input-to-hidden connections. **(b)** Hidden-to-output connections.

#### S1.3 Manipulations of sleep's weight changes

Main Fig. 4 compares real sleep with two manipulations of sleep's own weight changes, each applied to both pathways. Supplementary Figure S4 shows the full set of conditions (definitions in main text, Methods). A uniform shift of every synapse by its pathway's mean change ("Global") restored nothing (old-task accuracy 0.17 vs. 0.16 before sleep and 0.74 after real sleep; pooled over Sleeps 2–5, six trials). Removing target specificity (column equalization) abolished recovery when applied to both pathways (0.21) or when the comparison set was restricted to the older tasks' postsynaptic units (0.20), and reduced it when applied to one pathway only (0.50 for H→O, 0.36 for I→H). Removing source specificity (row equalization) in both pathways left old-task accuracy (0.75) and $Q$ (1.97) as high as after real sleep (0.74, 1.89), and reduced only the accuracy of the most recent task (0.75 vs. 0.79).

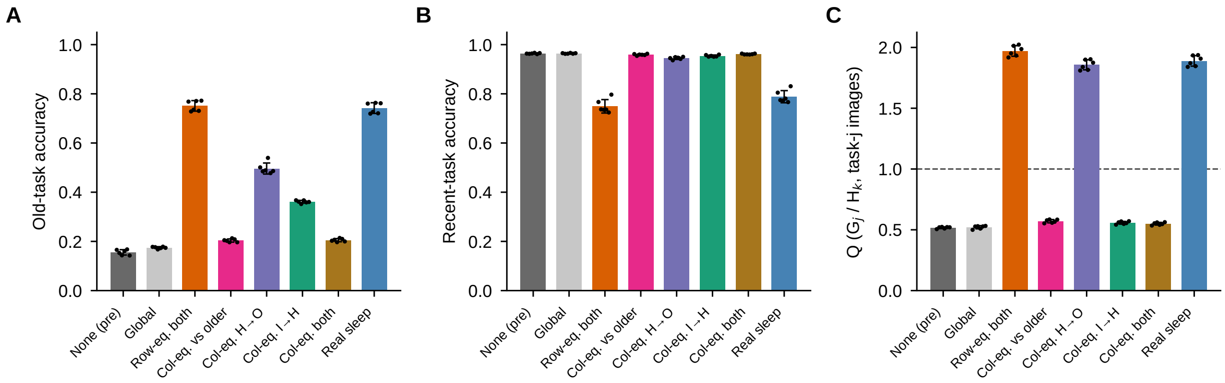


Figure S4: **Full set of manipulations of sleep's weight changes.** **(a)** Old-task accuracy, **(b)** recent-task accuracy and **(c)** intrusion index $Q$. Bar labels, from left: *None (pre)*, the pre-sleep network; *Global*, a uniform shift of every synapse by its pathway's mean change; *Row-eq. both*, source specificity removed in both pathways (main Fig. 4, "Sleep: SS removed"); *Col-eq. vs older*, a variant of target specificity removed in which each presynaptic unit's mean change onto the recent population is equalized only with its mean change onto the older tasks' populations, rather than onto all other units (neurons not yet assigned to any task are left unchanged); *Col-eq. H→O* and *Col-eq. I→H*, target specificity removed in that pathway only; *Col-eq. both*, target specificity removed in both pathways (main Fig. 4, "Sleep: TS removed"); *Real sleep*. "Row-eq." and "Col-eq." stand for row and column equalization of the weight-change matrix: equalization over each postsynaptic unit's inputs and over each presynaptic unit's outputs, respectively (main text, Methods). Pooled over Sleeps 2–5, mean ± SD over six trials; dots, individual trials.

#### S1.4 Shuffle control

As a further control, we randomly redistributed the real $\Delta W$ values produced by sleep across synapse positions (50 shuffles per stage), preserving the exact distribution of weight changes while destroying their targeting, first in the hidden-to-output pathway alone and then in all synapses. Shuffling only the hidden-to-output pathway substantially reduced recovery relative to real sleep (final stage: real $= 0.75$ vs. shuffled $= 0.38 \pm 0.01$, pre-sleep $= 0.03$; Supplementary Fig. S5); shuffling all synapses reduced performance to the pre-sleep level. The targeting of the weight changes, not merely their magnitude, therefore carries the recovery.

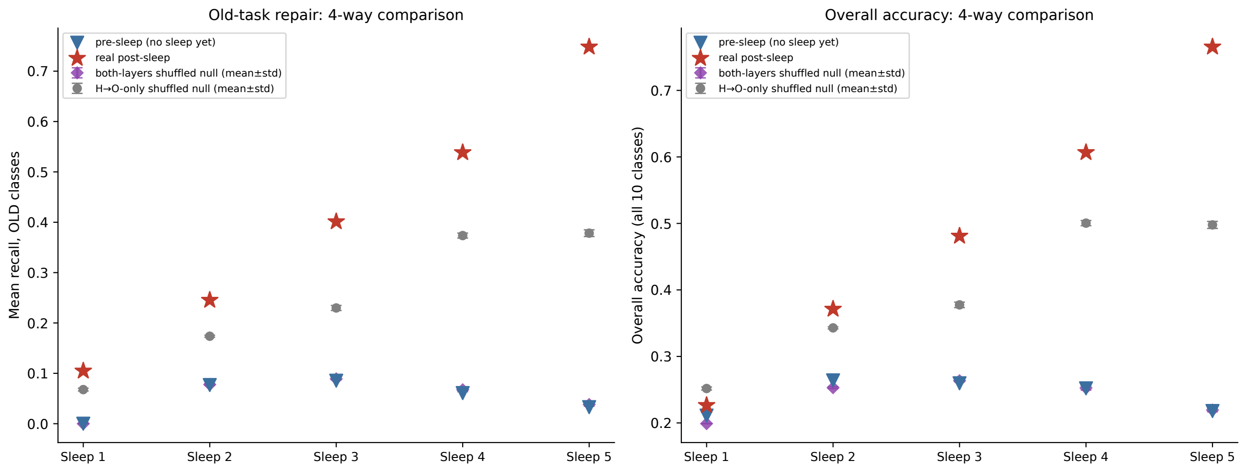


Figure S5: **Sleep's benefit depends on the targeting of weight changes, not merely their magnitude.** Old-task recall (left) and overall accuracy (right) at each sleep stage: pre-sleep (blue triangle), real post-sleep (red star), a shuffled control that redistributes sleep's own hidden-to-output $\Delta W$ values across synapses (gray circle, mean $\pm$ s.d. over 50 shuffles), and a second control that additionally shuffles the input-to-hidden pathway (purple diamond, mean $\pm$ s.d. over 30 shuffles).

#### S1.5 The analytical model and the full network

Eq. 1 of the main text gives the expected weight change of a synapse per replay step as

$$p_{\text{post}}\,[\,p_{\text{pre}}\,\text{inc} - (1 - p_{\text{pre}})\,\text{dec}\,],$$

assuming that pre- and postsynaptic spikes are independent. In the full network, the sleep plasticity rule updates a synapse whenever its postsynaptic unit spikes at step $t$, by $+\text{inc}$ if the presynaptic unit spiked at step $t-1$ and by $-\text{dec}$ otherwise (main text, Algorithm 1; $\text{inc} = 5.95 \times 10^{-4}$, $\text{dec} = 2.83 \times 10^{-4}$, $\text{dec}/\text{inc} = 0.475$). The weight change of a synapse over a sleep episode is therefore exactly $N_{\text{post}}\,[\,q\,\text{inc} - (1-q)\,\text{dec}\,]$, where $N_{\text{post}}$ is the number of postsynaptic spikes and $q$ the probability that the presynaptic unit was active in the step before a postsynaptic spike (main text, Eq. 3); independence corresponds to $q = p_{\text{pre}}$.

For every synapse of both pathways (input$\to$hidden and hidden$\to$output), in all six trials and Sleeps 2–5, we counted $N_{\text{post}}$, $q$ and the firing probabilities $p_{\text{pre}}$ and $p_{\text{post}}$ directly from the spike trains recorded during sleep, and compared the weight change predicted by Eq. 1 with $p_{\text{pre}}$ (independence) and with $q$ against the actual change (post- minus pre-sleep weights). Synapses were grouped with the same populations as in main Fig. 4: own (recent task $\to$ recent task), cross (older tasks $\to$ recent task), cross in the reverse direction (recent $\to$ older), and all others.

With $q$, Eq. 1 reproduced every synapse's weight change to within numerical precision (largest deviation $1.2 \times 10^{-7}$; Pearson $r = 1.00$ in both pathways; Supplementary Fig. S6e,f), as expected, since $q$ is the quantity the rule uses. The independence form performed differently in the two pathways (Supplementary Fig. S6c,d). For input$\to$hidden synapses, $q$ equaled $p_{\text{pre}}$ ($q/p_{\text{pre}} = 1.01$ in every synapse group), because the Poisson input does not depend on network activity; the independence form fit well ($r = 0.97$) and reproduced the own–cross difference in mean weight change ($-1.02$ vs. actual $-1.05 \times 10^{-3}$). For hidden$\to$output synapses, $q$ was about three times $p_{\text{pre}}$ ($q/p_{\text{pre}} = 3.1$), because hidden spikes drive the output units one step later. The independence form therefore fit less well ($r = 0.85$), overestimated the depression of own-pathway synapses two-fold ($-6.2$ vs. actual $-3.1 \times 10^{-3}$) and captured only a third of the own–cross difference ($-1.5$ vs.

actual $-4.5 \times 10^{-3}$). For cross synapses the correlation hardly matters: older-task hidden units are almost silent during sleep ($p_{\mathrm{pre}} = 0.003$, $q = 0.009$), so nearly every output spike depresses these synapses.

On the phase diagram of main Fig. 5a, all measured own and cross synapses lie on the depression side of the boundary (Supplementary Fig. S6a,b): with the network's dec/inc = 0.475, depression occurs whenever $q < \mathrm{dec}/(\mathrm{dec} + \mathrm{inc}) = 0.32$, and the mean $q$ was 0.20 (own) and 0.15 (cross) for input→hidden and 0.19 (own) and 0.01 (cross) for hidden→output synapses. Cross synapses lie further from the boundary than own synapses in both pathways and are therefore depressed more, as the analytical model predicts.

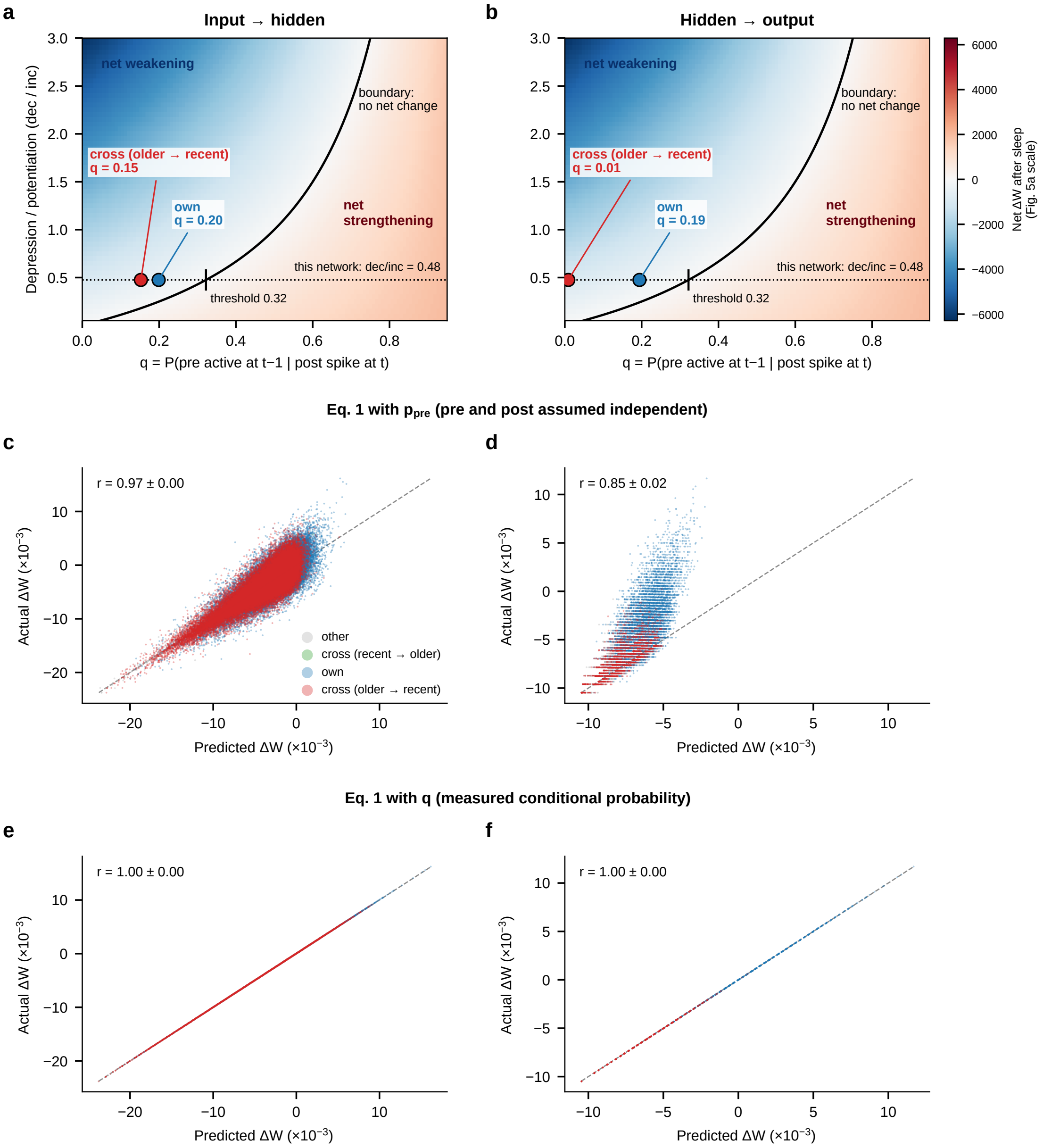


Figure S6: **The analytical model and the full network.** Left column: input→hidden synapses; right column: hidden→output synapses; six trials, Sleeps 2–5. **(a,b)** Mean conditional probability $q$ of own (blue) and cross (older → recent, red) synapses, placed on the phase diagram of main Fig. 5a at the network's dec/inc = 0.475 (dotted line). The background is Eq. 1 as in main Fig. 5a ($p_B = 0.7$, $T = 3000$); the horizontal axis starts at 0 so that the hidden→output cross synapses ($q = 0.01$) are visible. The boundary between net weakening and net strengthening does not depend on $p_{\text{post}}$; on the dotted line it lies at $q = 0.32$ (tick mark). **(c,d)** Weight change of each synapse over a sleep episode predicted by Eq. 1 with $p_{\text{pre}}$ (pre- and postsynaptic spikes assumed independent) versus the actual weight change. **(e,f)** The same with the measured $q$ in place of $p_{\text{pre}}$. (c–f) Random sample of synapses whose postsynaptic unit spiked during sleep (up to 3000 per synapse group, trial and sleep), coloured by group; dashed line, identity; $r$, Pearson correlation over all such synapses, mean ± SD over trials.

#### S2 Dale's-law-constrained MRNN-EP models

To test whether the mechanism survives a biologically motivated sign constraint on synaptic weights, we imposed Dale's law at the level of individual synapses (not individual neurons; see main text for why a neuron-level constraint is incompatible with EP's symmetric weights): each synapse in the input→hidden and hidden→output weight matrices was independently, randomly assigned a fixed excitatory or inhibitory identity at initialization, and its effective weight was computed as $W_{\text{eff}} = |W_{\text{raw}}| \times \text{sign}$ at every forward pass and after every plasticity update. During sleep, inhibitory synapses used a sign-swapped magnitude rule relative to excitatory synapses (the larger of the two STDP constants governs inhibitory strengthening rather than weakening), which outperformed the more naive implementation that reuses the excitatory update rule unchanged for inhibitory synapses. An additional sleep episode was applied before any task training (driven by noise with Task 1's input statistics); it is not shown in the figures. We report the model with a 50%/50% excitatory/inhibitory synapse ratio, which outperformed the more conventional 75%/25% ratio (68.9 ± 5.3% mean task accuracy); with GA-optimized sleep hyperparameters it reached $73.9 \pm 1.7\%$ mean task accuracy after the final sleep across six independently trained trials.

We also trained two ablations, identical in every respect except for which synapses were allowed to change during sleep: in the first, only excitatory synapses were plastic and inhibitory synapses were held fixed at their post-training value in every sleep episode; in the second, only inhibitory synapses were plastic and excitatory synapses were held fixed. Sleep hyperparameters were independently re-optimized by genetic algorithm for each ablation.

Supplementary Table S1 summarizes the three models with the definitions of main Fig. 7. All three recovered older tasks to the same extent and decorrelated the hidden layer. All three showed high target specificity in both pathways. The fully plastic and inhibitory-only models also showed source specificity, largest in the hidden→output pathway, whereas the excitatory-only model showed essentially none: all excitatory inputs onto the reactivated population were depressed to a similar extent, regardless of their source. Its recovery of older tasks was nevertheless undiminished, confirming that target specificity, not source specificity, carries recovery (main Fig. 4). Its lower final accuracy after the last sleep reflects weaker recovery of the older tasks at that last episode (Tasks 1–4: 0.63 vs. 0.76 for the fully plastic model), while the most recent task was retained better (0.83 vs. 0.65).

Table S1: **Dale's-law models: accuracy, decorrelation and specificity of sleep's weight changes**, computed as in main Fig. 7 (per trial, mean over Sleeps 2–5; mean ± SD over six trials). Final accuracy: mean over the five tasks after the last sleep. Old-task accuracy and $R$: before → after sleep. TS, SS: target and source specificity. The main MRNN-EP model is shown for reference.

| Model | Final acc. | Old-task acc. | $R$ | TS I→H | TS H→O | SS I→H | SS H→O |
|---|---|---|---|---|---|---|---|
| MRNN-EP (main) | $0.706 \pm 0.032$ | $0.16 \to 0.74$ | $0.71 \to 0.21$ | $0.92 \pm 0.01$ | 1.00 | $0.08 \pm 0.01$ | $0.46 \pm 0.02$ |
| Dale, fully plastic | $0.739 \pm 0.017$ | $0.11 \to 0.76$ | $0.67 \to 0.25$ | $0.94 \pm 0.00$ | 1.00 | $0.12 \pm 0.02$ | $0.62 \pm 0.01$ |
| Dale, excitatory only | $0.669 \pm 0.016$ | $0.08 \to 0.74$ | $0.71 \to 0.25$ | $0.89 \pm 0.01$ | 1.00 | $0.03 \pm 0.02$ | $0.01 \pm 0.04$ |
| Dale, inhibitory only | $0.727 \pm 0.021$ | $0.11 \to 0.76$ | $0.69 \to 0.26$ | $0.92 \pm 0.00$ | 1.00 | $0.10 \pm 0.01$ | $0.66 \pm 0.02$ |

##### S2.1 Fully plastic Dale's-law model

All signatures of the main model were reproduced (Supplementary Figs. S7, S8). $R$ fell from 0.62–0.74 after every training block to 0.17–0.31 after every sleep episode, the mean pairwise Jaccard overlap from 0.32–0.44 to 0.10–0.15, and $Q$ rose from 0.55–0.64 before to 1.40–2.12 after Sleeps 2–5. During each sleep, the population of the task just trained dominated reactivation (mean rate 0.140 vs. 0.023 for the other tasks' populations), and the weight changes were concentrated on the synapses onto that population.

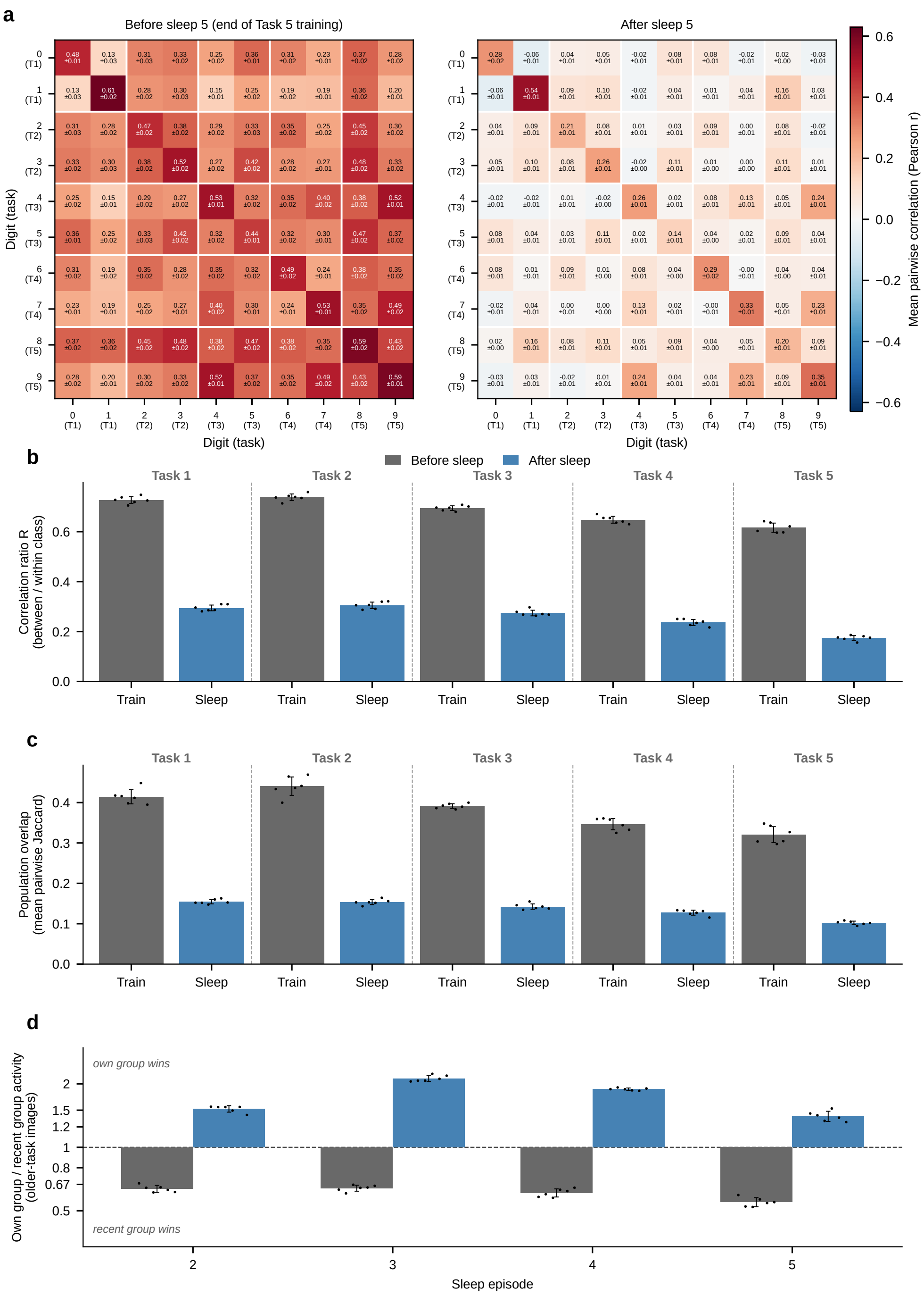


Figure S7: **Dale's-law MRNN-EP (50%/50% excitatory/inhibitory synapses, fully plastic): representations** (cf. main Fig. 2). **(a)** Class-level hidden-activation correlation at the end of Task 5 training and after the final sleep, mean ± SD over six trials. **(b)** Correlation ratio $R$ at the end of each training block (grey) and after the following sleep (blue). **(c)** Mean pairwise Jaccard overlap of the top-10% class populations. **(d)** Intrusion index $Q$ before and after Sleeps 2–5. (b–d) Mean ± SD over six trials; dots, individual trials. For all supplementary models, $Q$ was computed from the logged test-split activations; for the main model this gives values within $\sim 2\%$ of the validation-split $Q$ of main Fig. 2d.

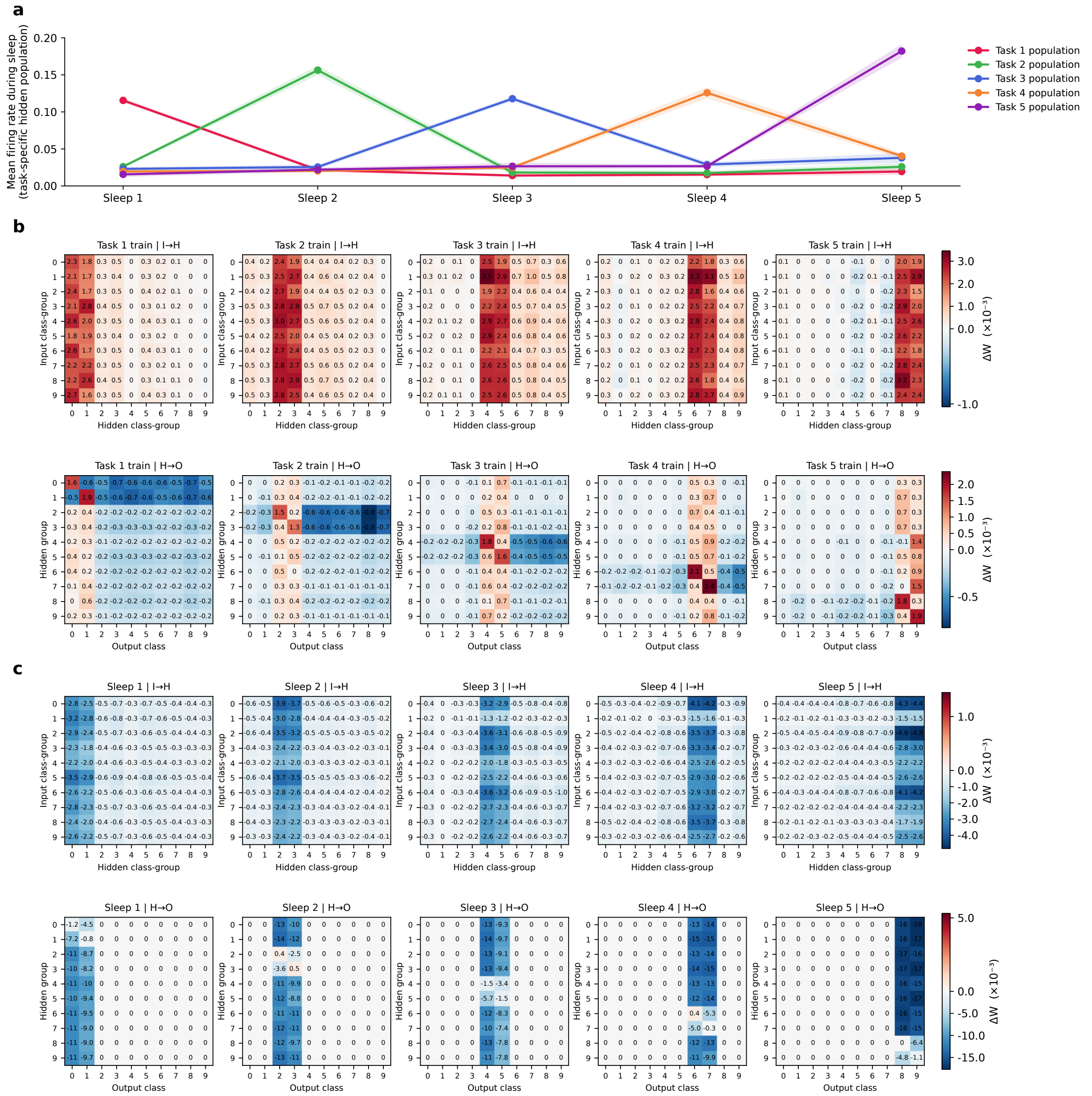

Figure S8: **Dale's-law MRNN-EP (fully plastic): reactivation and weight changes** (cf. main Fig. 3). **(a)** Mean firing rate of each task's hidden population during each sleep episode, mean ± SD over six trials. **(b)** Block-averaged weight changes during the training of each task (top: input→hidden; bottom: hidden→output). **(c)** Block-averaged weight changes during each sleep episode. (b,c) Trial 0; effective weights $|W| \times \text{sign}$; units of $\Delta W \times 10^{-3}$; the pre-training sleep episode is not shown.

#### S2.2 Excitatory-only sleep plasticity

With only excitatory synapses plastic during sleep, $R$ fell from 0.66–0.76 after every training block to 0.18–0.30 after every sleep episode, the Jaccard overlap from 0.35–0.44 to 0.10–0.16, and $Q$ rose from 0.47–0.60 to 1.26–2.01 (Supplementary Fig. S9). The task just trained again dominated reactivation (0.217 vs. 0.043), and the weight changes – here depression of excitatory synapses only – formed the same vertical bands onto its population, with little difference between older-task and own-task sources (Supplementary Fig. S10).

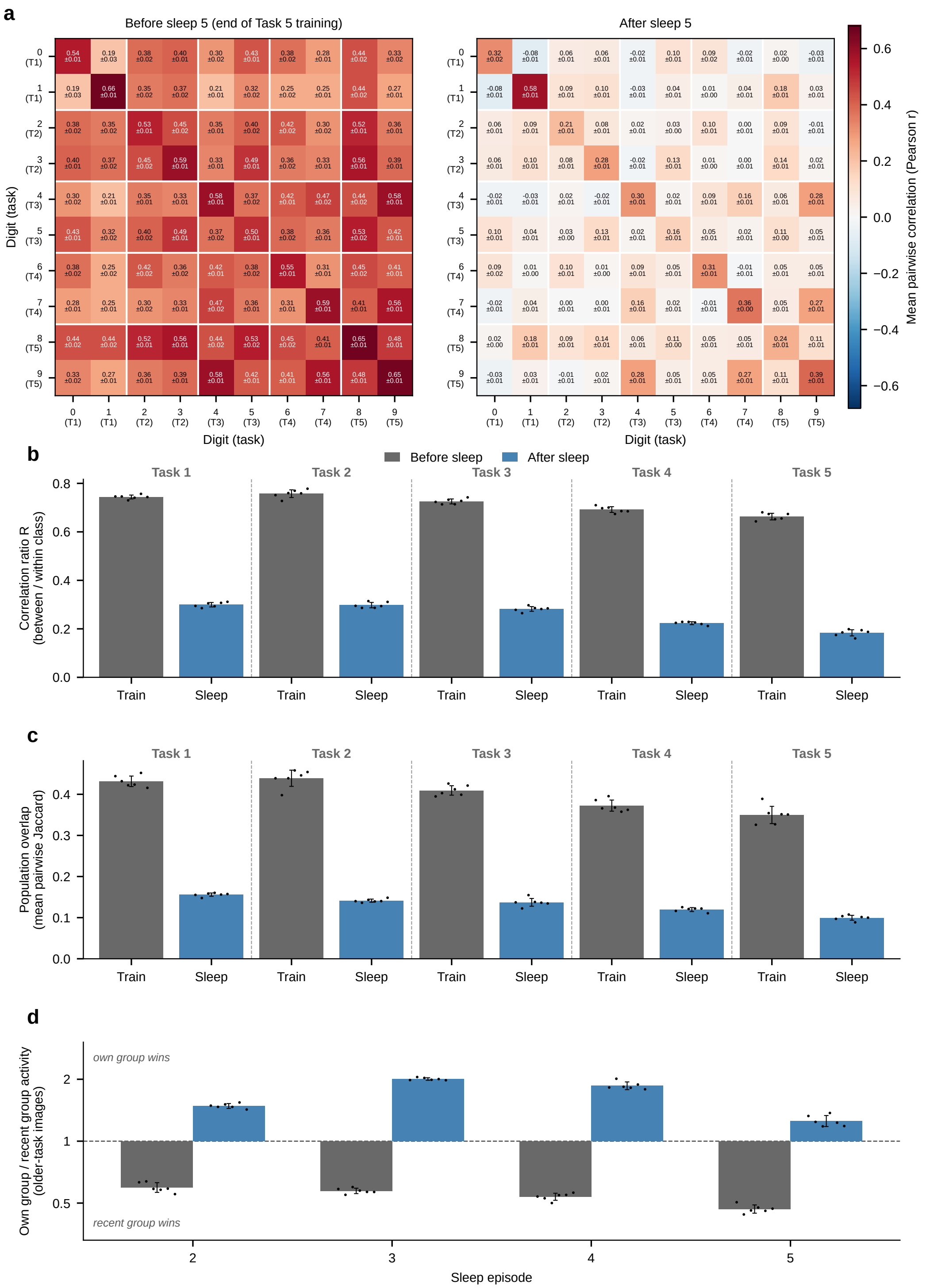


Figure S9: **Dale's-law MRNN-EP with excitatory-only sleep plasticity: representations.** Panels as in Supplementary Fig. S7.

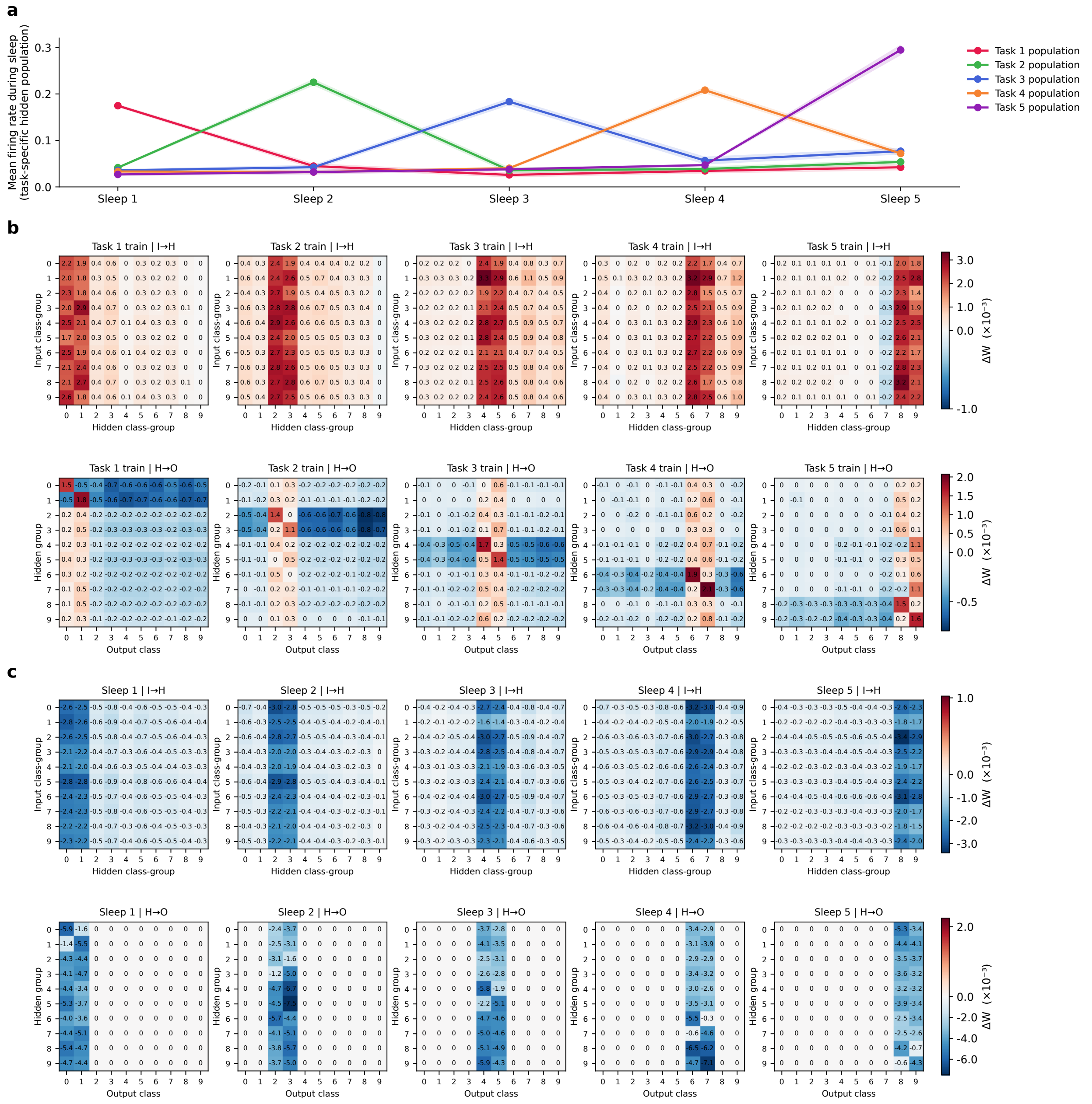


Figure S10: **Dale's-law MRNN-EP with excitatory-only sleep plasticity: reactivation and weight changes.** Panels as in Supplementary Fig. S8; inhibitory synapses do not change during sleep in this model.

#### S2.3 Inhibitory-only sleep plasticity

With only inhibitory synapses plastic during sleep, $R$ fell from 0.64–0.74 to 0.18–0.32, the Jaccard overlap from 0.33–0.43 to 0.10–0.16, and $Q$ rose from 0.50–0.60 to 1.36–1.88 (Supplementary Fig. S11); reactivation was again dominated by the task just trained (0.199 vs. 0.037), and the weight changes were concentrated on its population (Supplementary Fig. S12).

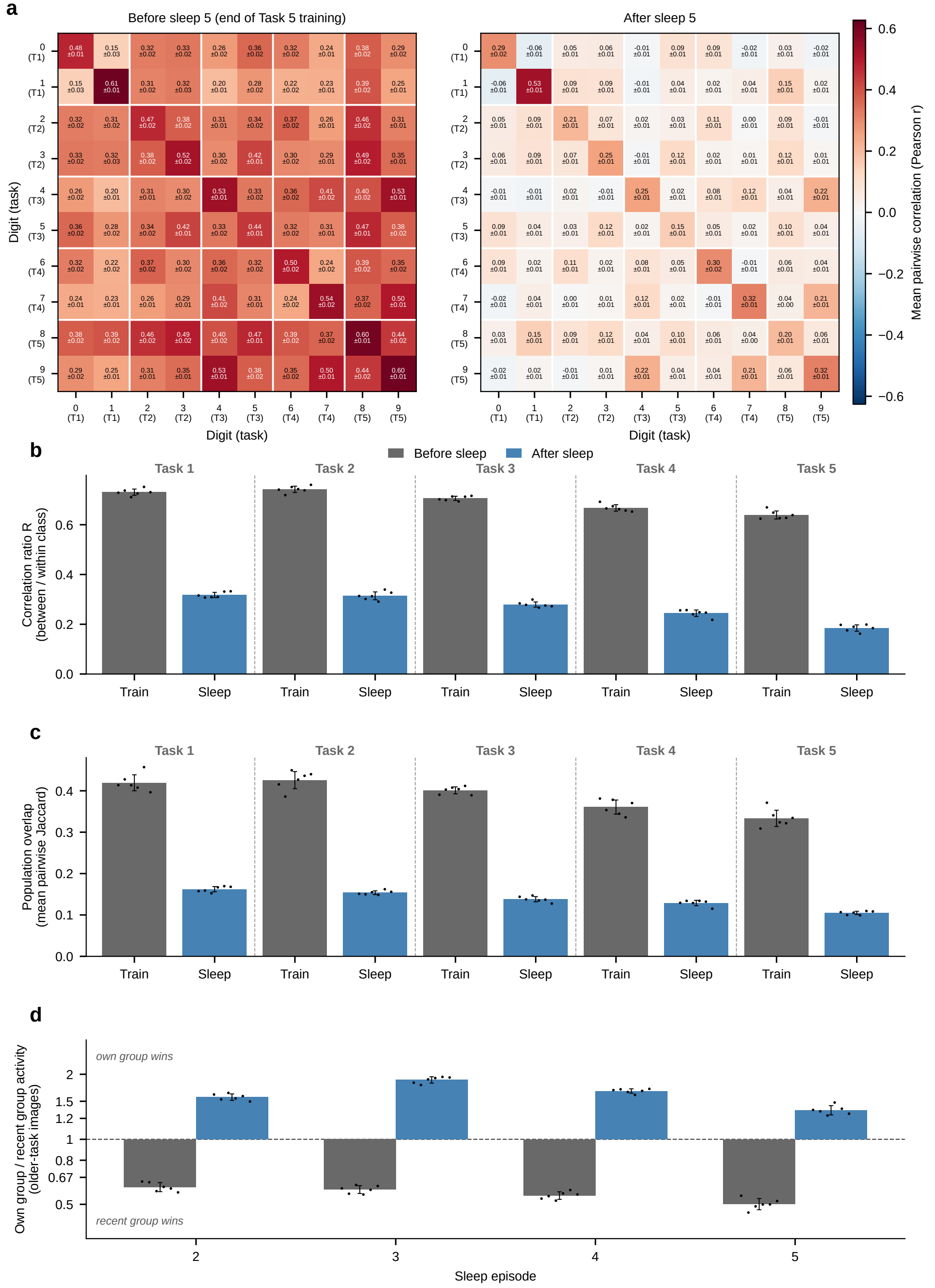


Figure S11: **Dale's-law MRNN-EP with inhibitory-only sleep plasticity: representations.** Panels as in Supplementary Fig. S7.

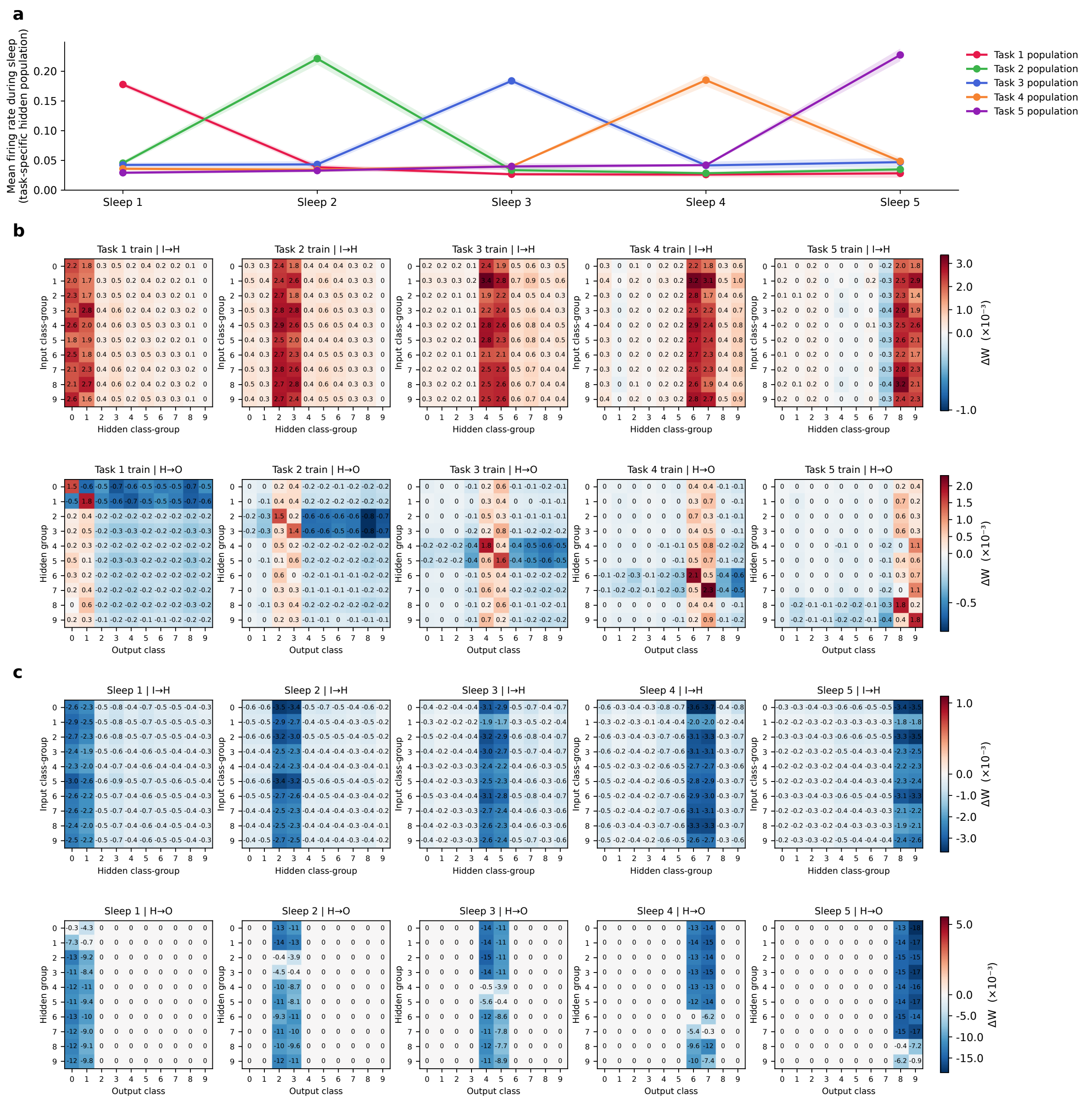


Figure S12: **Dale's-law MRNN-EP with inhibitory-only sleep plasticity: reactivation and weight changes.** Panels as in Supplementary Fig. S8; excitatory synapses do not change during sleep in this model.

### S3 Backpropagation-through-time (BPTT) trained RNN

To test whether the mechanism is specific to Equilibrium-Propagation (EP) training, we trained an architecturally matched recurrent network (same layer sizes, hidden-layer recurrence, same sleep input) using standard backpropagation-through-time (BPTT) instead of EP as the waking learning rule. Unlike the EP model, this network has no feedback pathway from the output layer back to the hidden layer during waking training; hidden-to-hidden recurrence is its only recurrent connection. Sleep hyperparameters (sleep duration, input gain, firing-rate scale, membrane decay, and the STDP increment/decrement constants) were re-optimized by genetic algorithm for this model, independently of the EP model's values. Over six independently trained trials, the model reached $62.6 \pm 2.9\%$ mean task accuracy.

$R$ fell from 0.55–0.73 after every training block to 0.14–0.35 after every sleep episode, the Jaccard overlap from 0.29–0.38 to 0.10–0.17, and $Q$ rose from 0.48–0.60 to 1.78–2.45 (Supplementary Fig. S13). The task just trained dominated reactivation (0.049 vs. 0.010), and in all three pathways – including the hidden$\rightarrow$hidden recurrence – the weight changes were concentrated on the synapses onto its population (Supplementary Fig. S14). In the recurrent pathway, connections from older-task hidden groups onto the just-trained group were depressed more than connections from the just-trained group onto itself (own$\rightarrow$own vs. older$\rightarrow$own: $-3.93$ vs. $-4.76 \times 10^{-3}$ after Task 2 sleep; $-3.10$ vs. $-3.58 \times 10^{-3}$ after Task 3; $-3.06$ vs. $-3.43 \times 10^{-3}$ after Task 4; $-4.90$ vs. $-5.87 \times 10^{-3}$ after Task 5), a 12–21% difference; this modest source specificity, also seen in the other two pathways of this model (main Fig. 7c), did not limit recovery (main Fig. 7d).

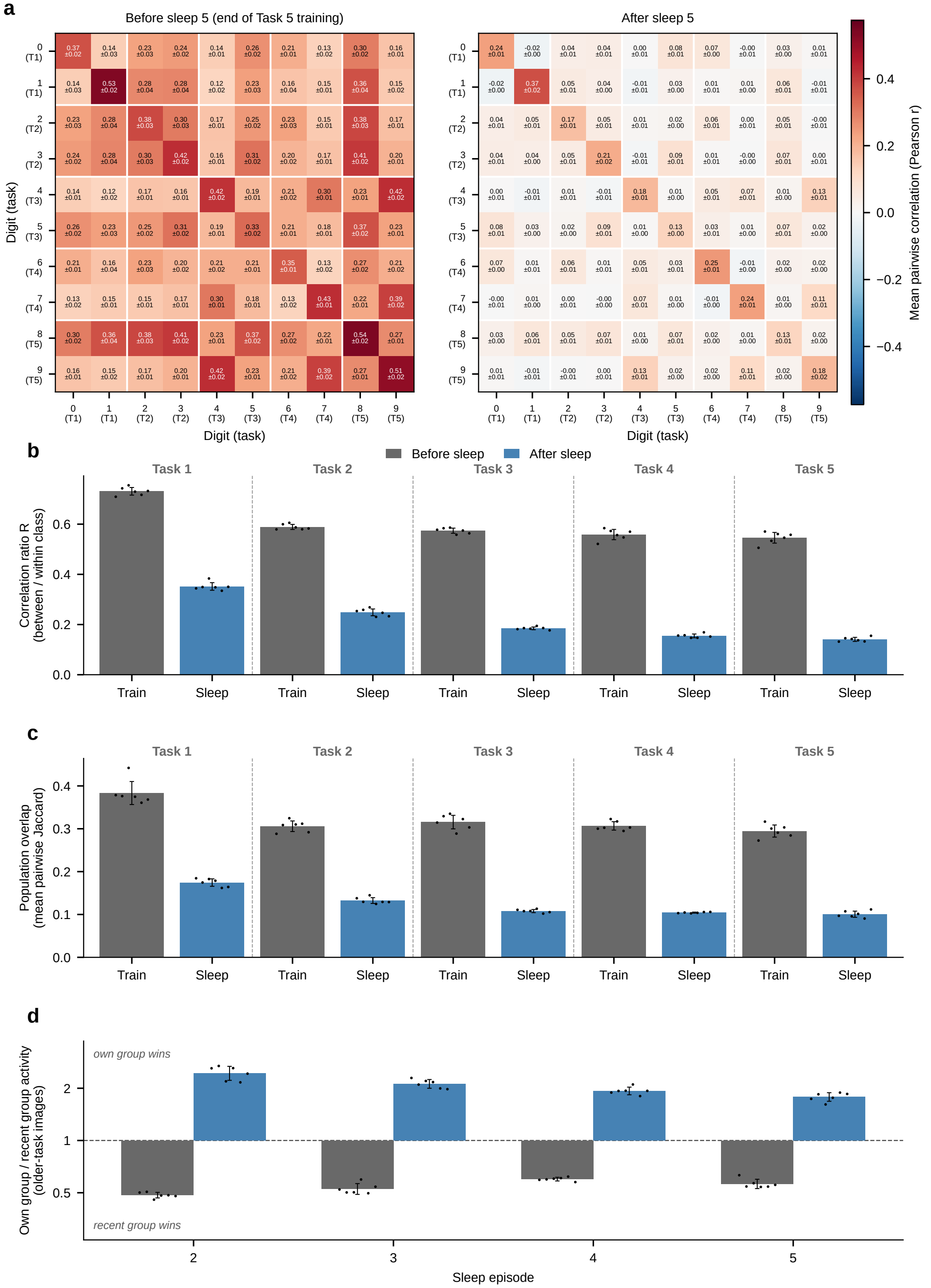


Figure S13: **RNN trained with BPTT: representations.** Panels as in Supplementary Fig. S7; mean ± SD over six trials.

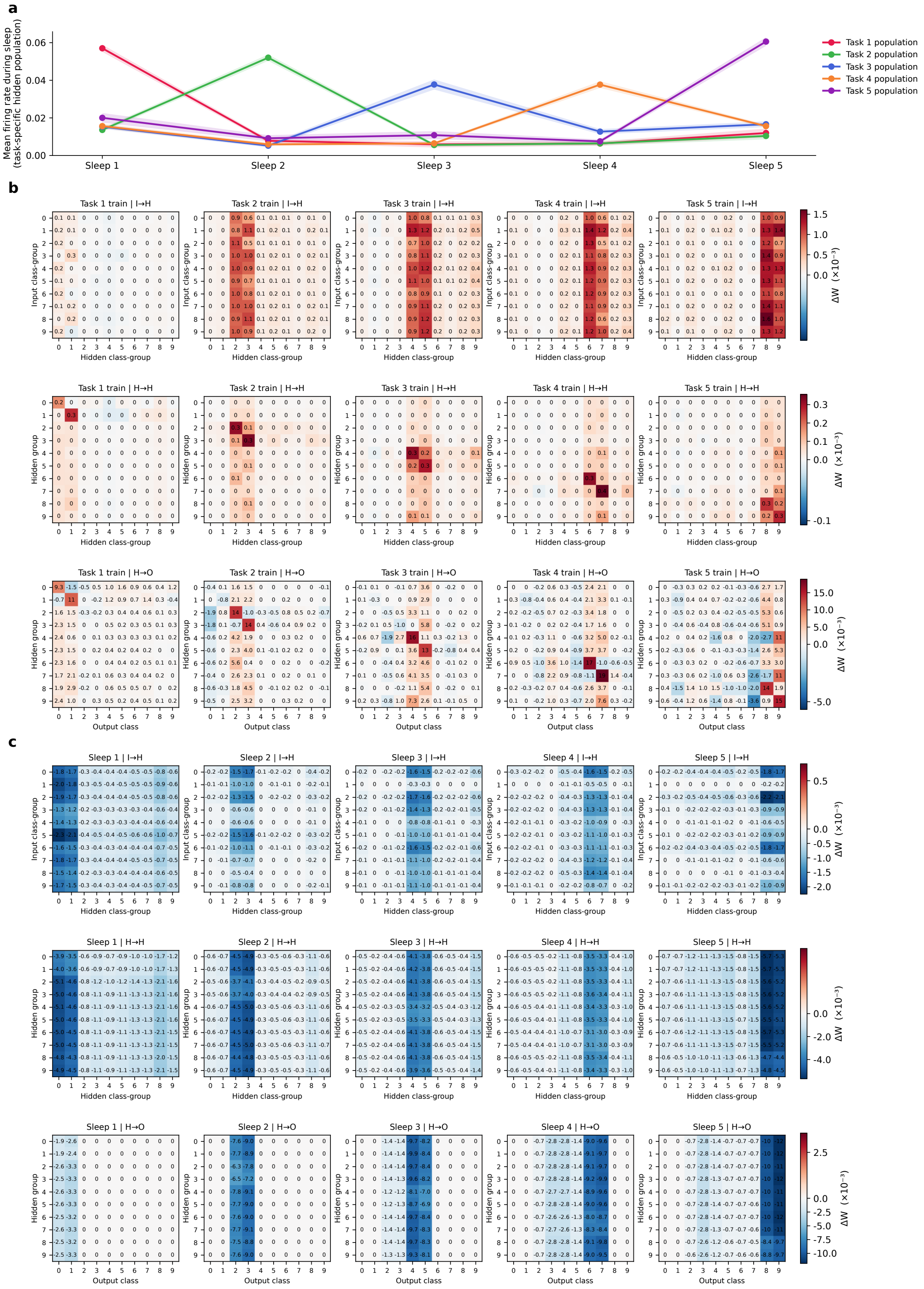


Figure S14: **RNN trained with BPTT: reactivation and weight changes.** Panels as in Supplementary Fig. S8, with an additional row for the hidden→hidden recurrent pathway (rows: presynaptic, columns: postsynaptic hidden class groups). (b,c) Trial 0.

#### S4 MRNN-EP trained on CIFAR-10

To test whether the mechanism depends on the simple statistics of MNIST, we trained MRNN-EP on five two-class tasks of CIFAR-10 natural images (classes 0–1, 2–3, . . . , 8–9). Images were passed through the convolutional layers of a pretrained VGG-13 network that were held fixed; their output (8192 features) was the input to MRNN-EP (8192–1024–10), and only the EP network learned during waking and changed during sleep. Training used 5 epochs per task and each sleep episode replayed only the most recently trained task, as for MNIST; sleep hyperparameters were optimized by genetic algorithm, starting from those of ref. [2] (main text, Methods). Over six independently trained trials, the model reached $83.5 \pm 0.7\%$ mean task accuracy after the final sleep. Input groups for the block matrices were the top-75 class-discriminative VGG-13 features rather than pixels.

$R$ fell from 0.77–0.91 after every training block to 0.18–0.61 after every sleep episode, the Jaccard overlap from 0.37–0.64 to 0.08–0.32, and $Q$ rose from 0.36–0.38 to 1.96–2.54 (Supplementary Fig. S15). The task just trained dominated reactivation (0.031 vs. 0.006 for older tasks), and the weight changes were concentrated on the synapses onto its population in both pathways (Supplementary Fig. S16; target specificity 0.79 for input→hidden and 1.00 for hidden→output, main Fig. 7b), with small source specificity that increased from the input to the output pathway (0.04 and 0.12; main Fig. 7c).

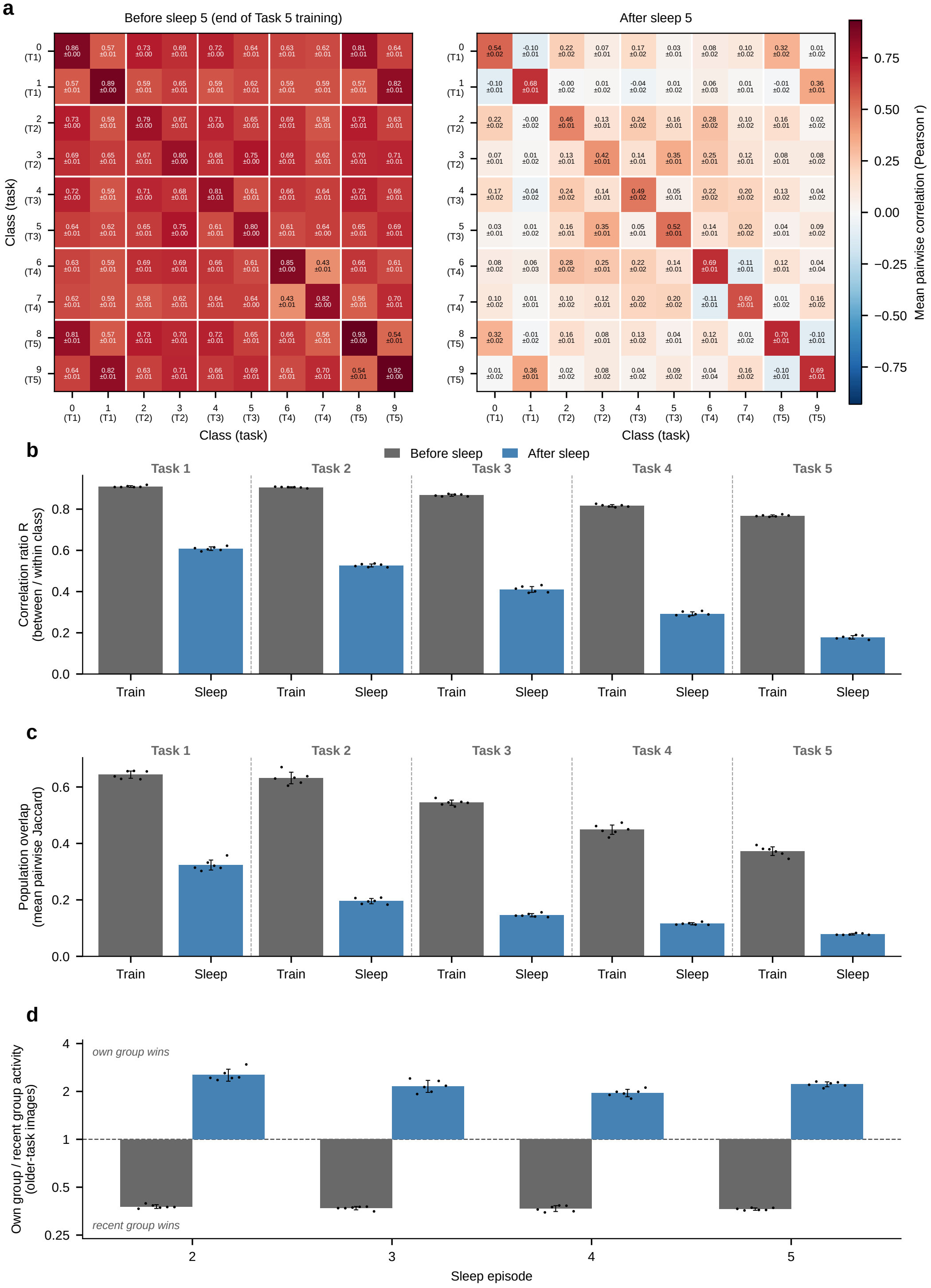


Figure S15: **MRNN-EP trained on CIFAR-10: representations.** Panels as in Supplementary Fig. S7; mean ± SD over six trials.

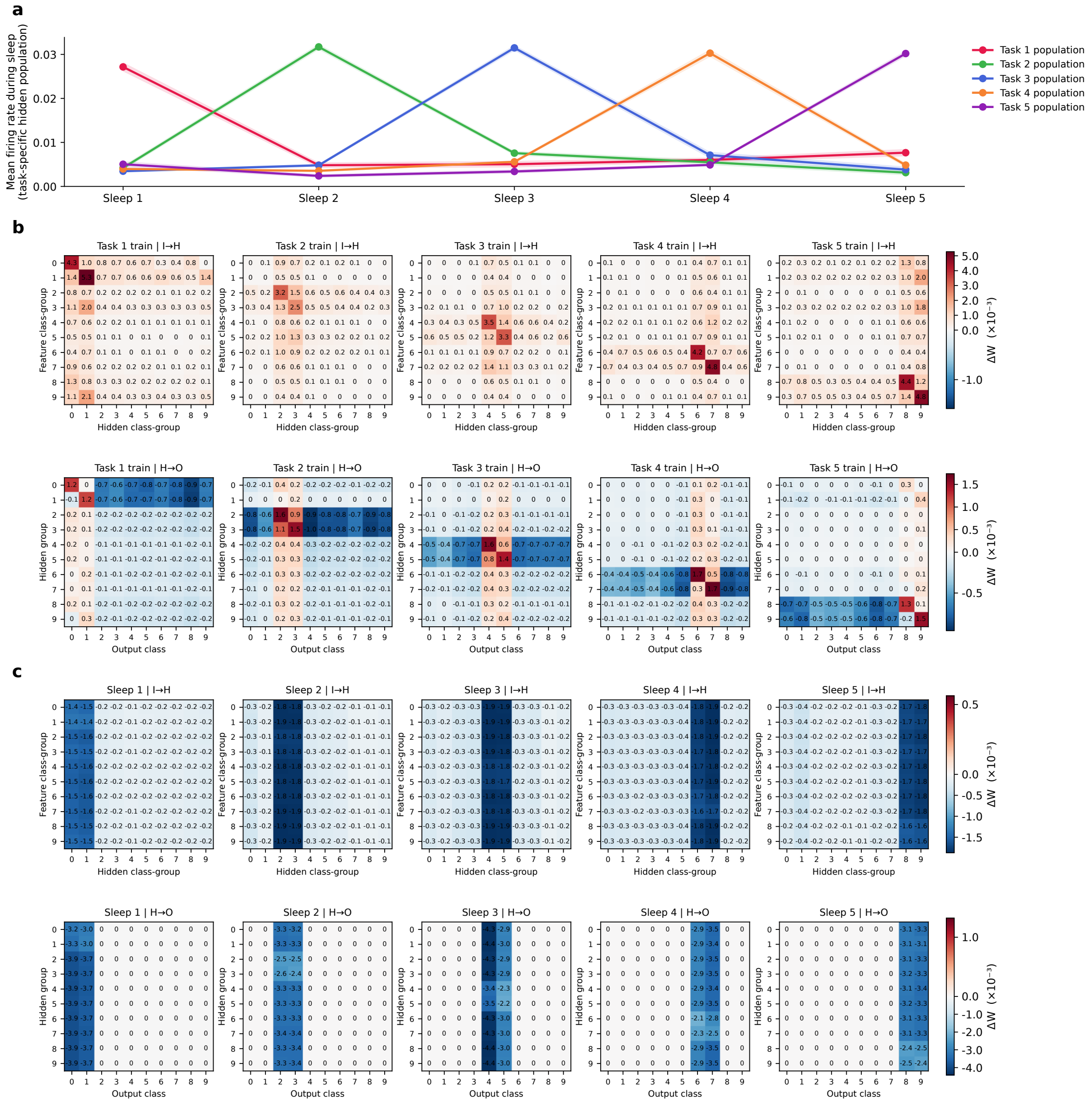


Figure S16: **MRNN-EP trained on CIFAR-10: reactivation and weight changes.** Panels as in Supplementary Fig. S8, with VGG-13 feature groups in place of pixel groups. (b,c) Trial 0.

### S5 Feedforward networks with Sleep Replay Consolidation

To test whether the signatures reported for the recurrent models generalize to feedforward architectures, we analyzed feedforward networks trained with backpropagation and consolidated with the original SRC procedure [1] on the same five-task, class-incremental curricula (two classes per task), with sleep driven, as in the main model, by the input statistics of the most recently trained task. Class-selective groups were defined with the same fixed-group convention as the main text: computed once per class from that class's own task at its last waking training epoch, then reused across all subsequent sleep stages.

#### S5.1 MNIST feedforward model

This network has three trainable layers – input→H1 (784→1200), H1→H2 (1200→1200), and H2→O (1200→10) – trained sequentially on the 5-task MNIST curriculum with SRC-driven sleep after each task, over six independently trained trials. Sleep restored older tasks (old-task accuracy 0.00 → 0.77 pooled over Sleeps 2–5; main Fig. 6d). In the second hidden layer, $R$ fell from 0.79–0.91 after every training block to 0.16–0.48 after every sleep episode, the Jaccard overlap from 0.49–0.63 to 0.12–0.28, and $Q$ rose from 0.16–0.23 to 1.73–4.99; the first hidden layer showed the same changes in weaker form ($R$: 0.67–0.81 to 0.35–0.45; $Q$: 0.70–0.76 to 1.11–1.47; Supplementary Fig. S17). In all three pathways, the weight changes were concentrated on the synapses onto the population of the task just trained (Supplementary Fig. S18), with source specificity increasing with depth (main Fig. 7c).

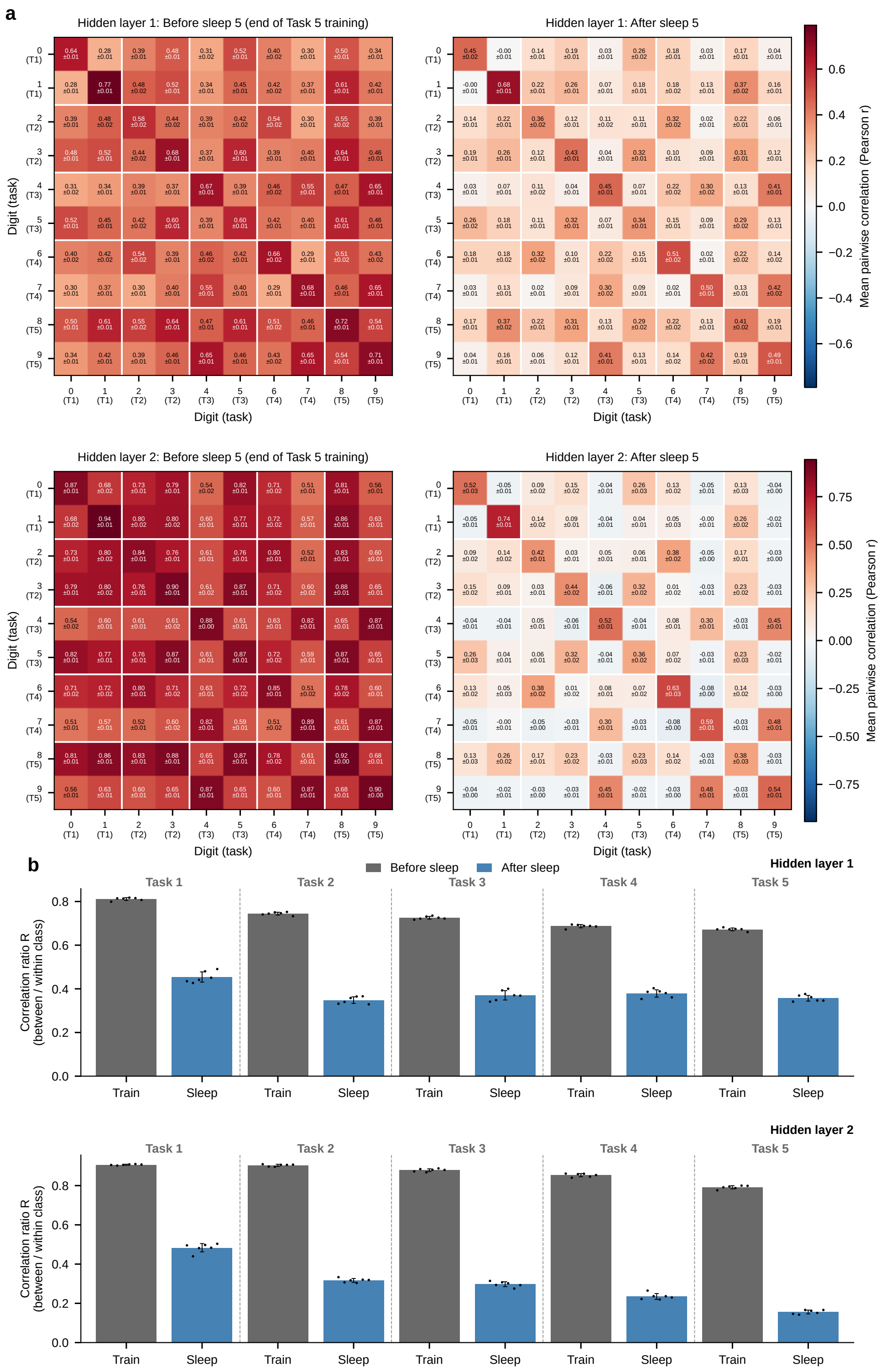


Figure S17: **MNIST feedforward SRC model: representations.** Panels as in Supplementary Fig. S7, shown separately for the two hidden layers (H1, H2); mean ± SD over six trials. Panels (c,d) continue on the next page.

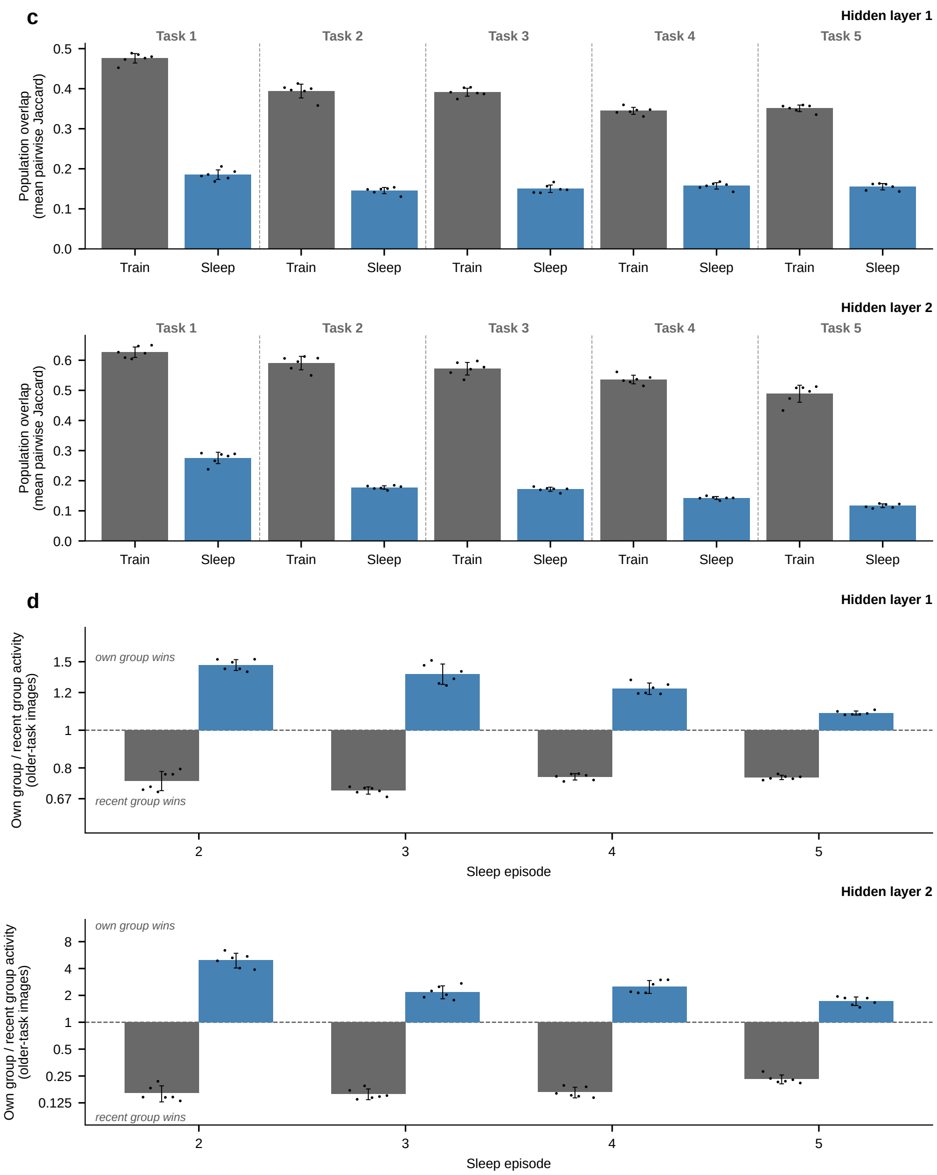


Figure S17: (continued).

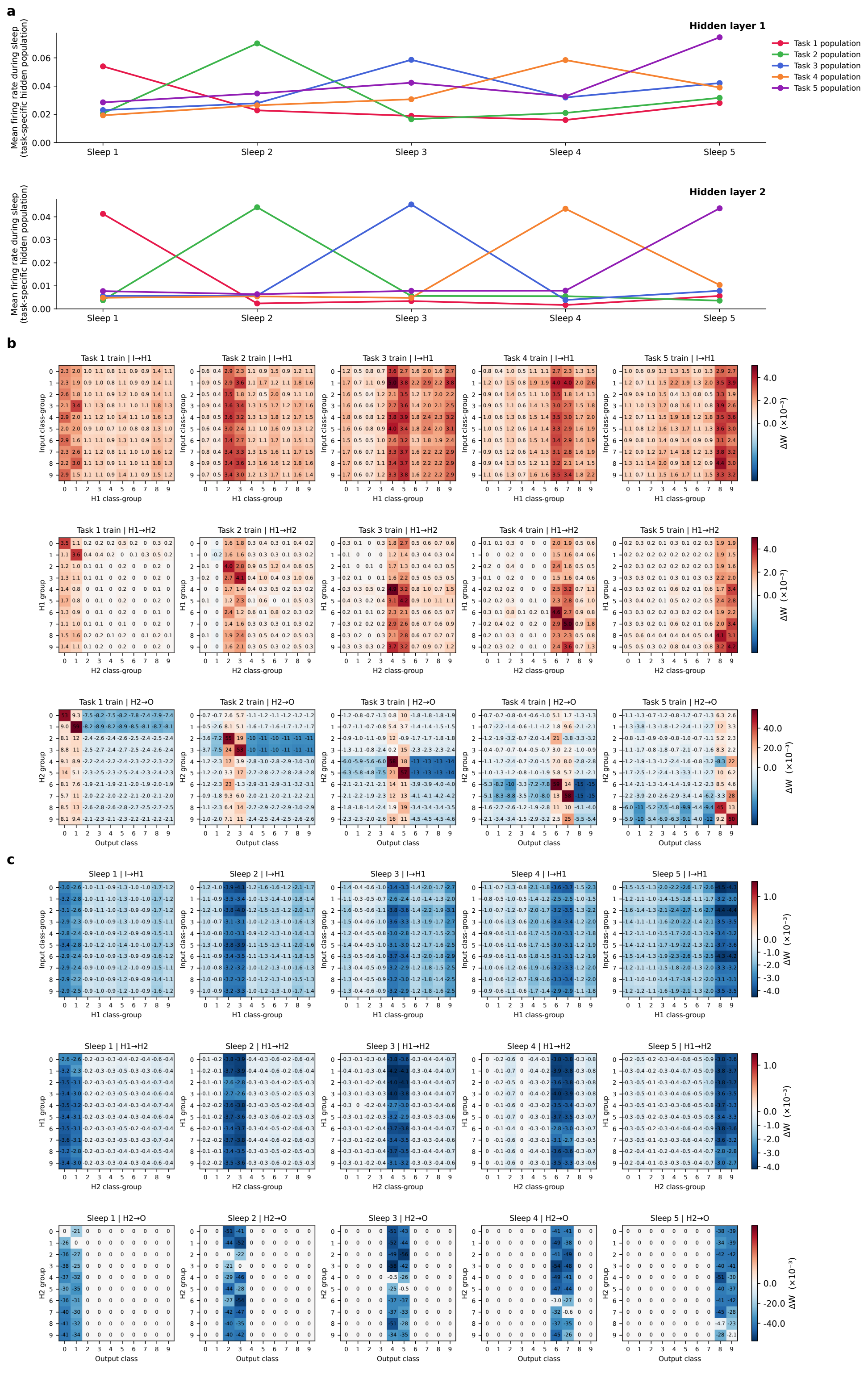


Figure S18: **MNIST feedforward SRC model: reactivation and weight changes.** Panels as in Supplementary Fig. S8, with one row per hidden layer in (a) and one row per pathway (input→H1, H1→H2, H2→O) in (b,c). Trial 0.

#### S5.2 CIFAR-10 feedforward model

This network is a small convolutional network (two convolutional layers followed by a fully connected head, 1800→1200→10) pretrained on all classes; its convolutional layers were then frozen, and the head was trained sequentially on the five CIFAR-10 tasks and consolidated with SRC-driven sleep of the head after each task (single trial). Input groups for the block matrices were the top-75 class-discriminative convolutional features rather than pixels. $R$ fell from 0.80–0.85 after every training block to 0.35–0.41 after every sleep episode, the Jaccard overlap from 0.36–0.43 to 0.14–0.19, and $Q$ rose from 0.46–0.56 to 1.64–2.57 (Supplementary Fig. S19); the weight changes again concentrated on the synapses onto the population of the task just trained (Supplementary Fig. S20).

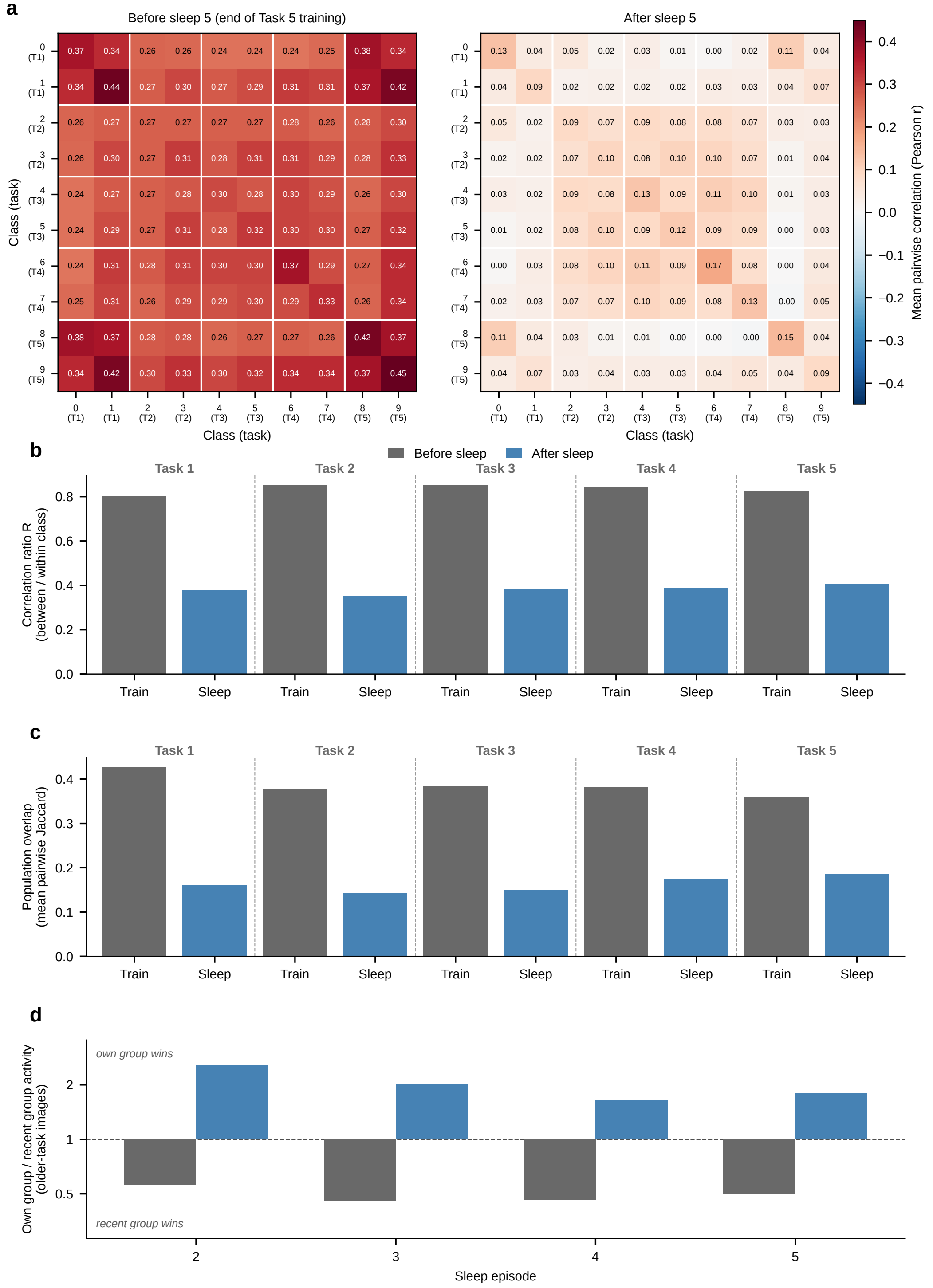


Figure S19: **CIFAR-10 feedforward SRC model: representations** (1200-unit hidden layer of the head). Panels as in Supplementary Fig. S7; single trial.

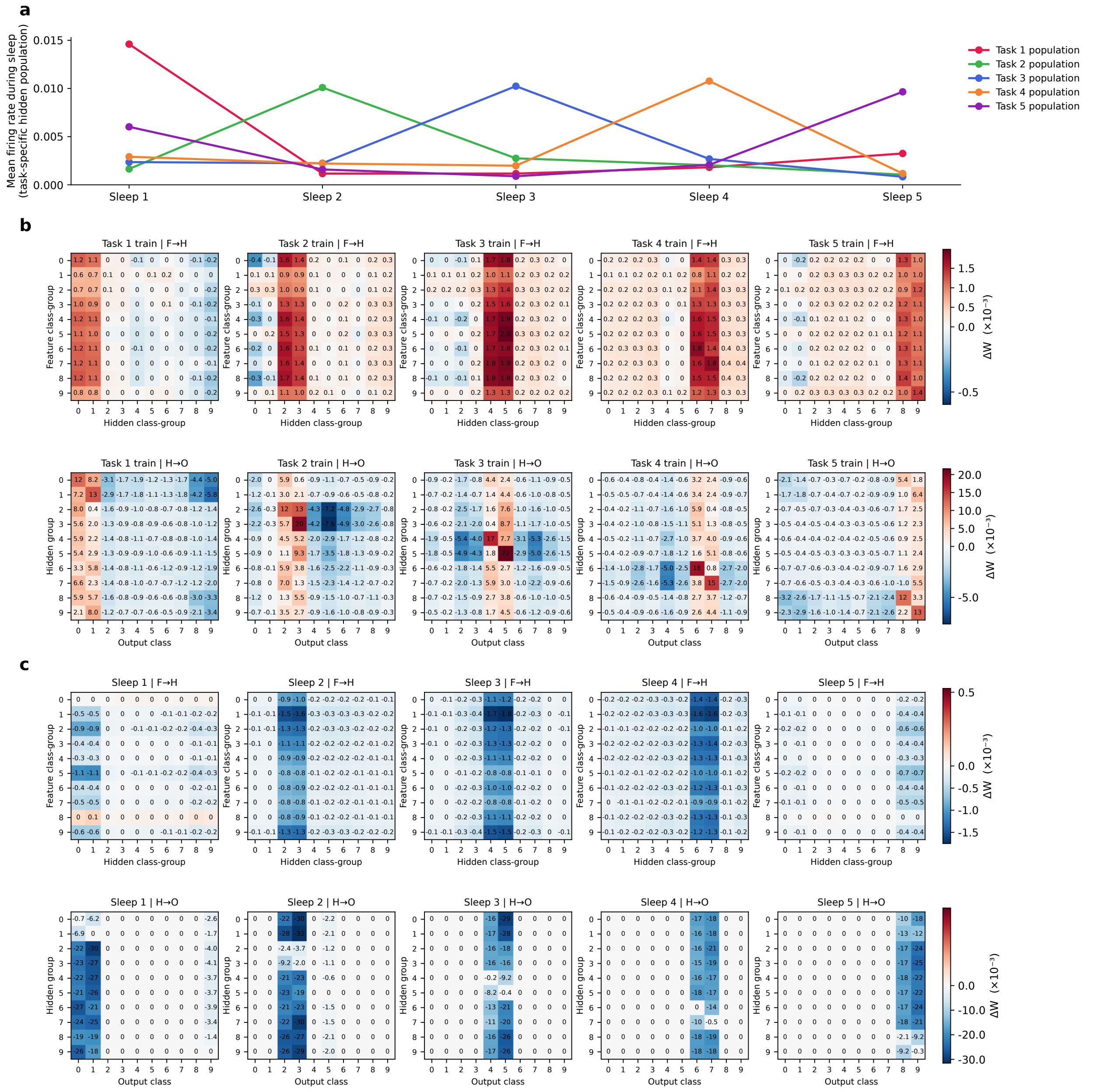


Figure S20: **CIFAR-10 feedforward SRC model: reactivation and weight changes.** Panels as in Supplementary Fig. S8, with convolutional-feature groups in place of pixel groups; single trial. Training of Task 1 starts from the end of the all-class pretraining.

#### S6 Sleep input based on all learned tasks

In the main model, each sleep episode is driven only by the input statistics of the task just trained. To test whether the mechanism depends on this choice, we trained an otherwise identical, unconstrained MRNN-EP model (3072 hidden units) in which sleep episode $k$ replays a mixture of tasks 1 through $k$. Sleep hyperparameters were independently optimized by genetic algorithm for this sleep input and network width. Over six independently trained trials, the model reached $65.4 \pm 4.7\%$ mean task accuracy after the final sleep.

Even though every learned task contributed to the sleep input, the most recent task's population still dominated reactivation, firing 2.0–3.3 times the mean rate of the older tasks' populations at Sleeps 2–5 (2.6 times on average, vs. 6.9 times with sleep input based on the most recent task only), and the older populations' reactivation declined with the age of the task (e.g., sleep 5: Tasks 1–5, 0.008, 0.011, 0.016, 0.023, 0.048; Supplementary Fig. S22a). $R$ fell from 0.50–0.76 after every training block to 0.29–0.38 after every sleep episode, the Jaccard overlap from 0.24–0.42 to 0.14–0.17, and $Q$ rose from 0.74–0.93 to 1.00–1.27 (Supplementary Fig. S21); the weight changes were concentrated on the synapses onto the population of the most recent task (Supplementary Fig. S22). The effects were weaker than with sleep input based on the most recent task only, consistent with the weaker dominance of the recent task during reactivation, but all signatures were present.

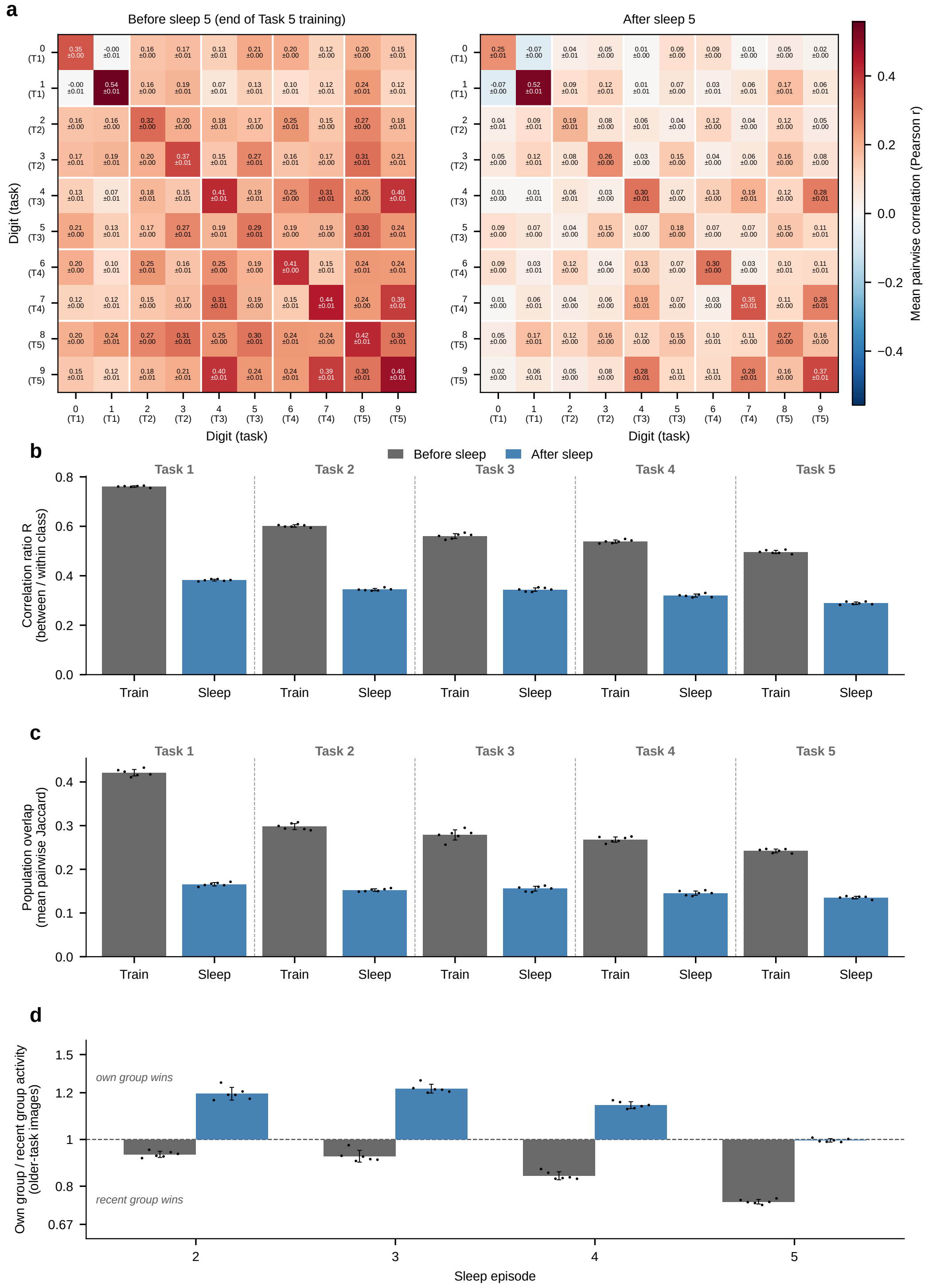


Figure S21: **MRNN-EP with sleep input based on all learned tasks (3072 units): representations.** Panels as in Supplementary Fig. S7; mean ± SD over six trials.

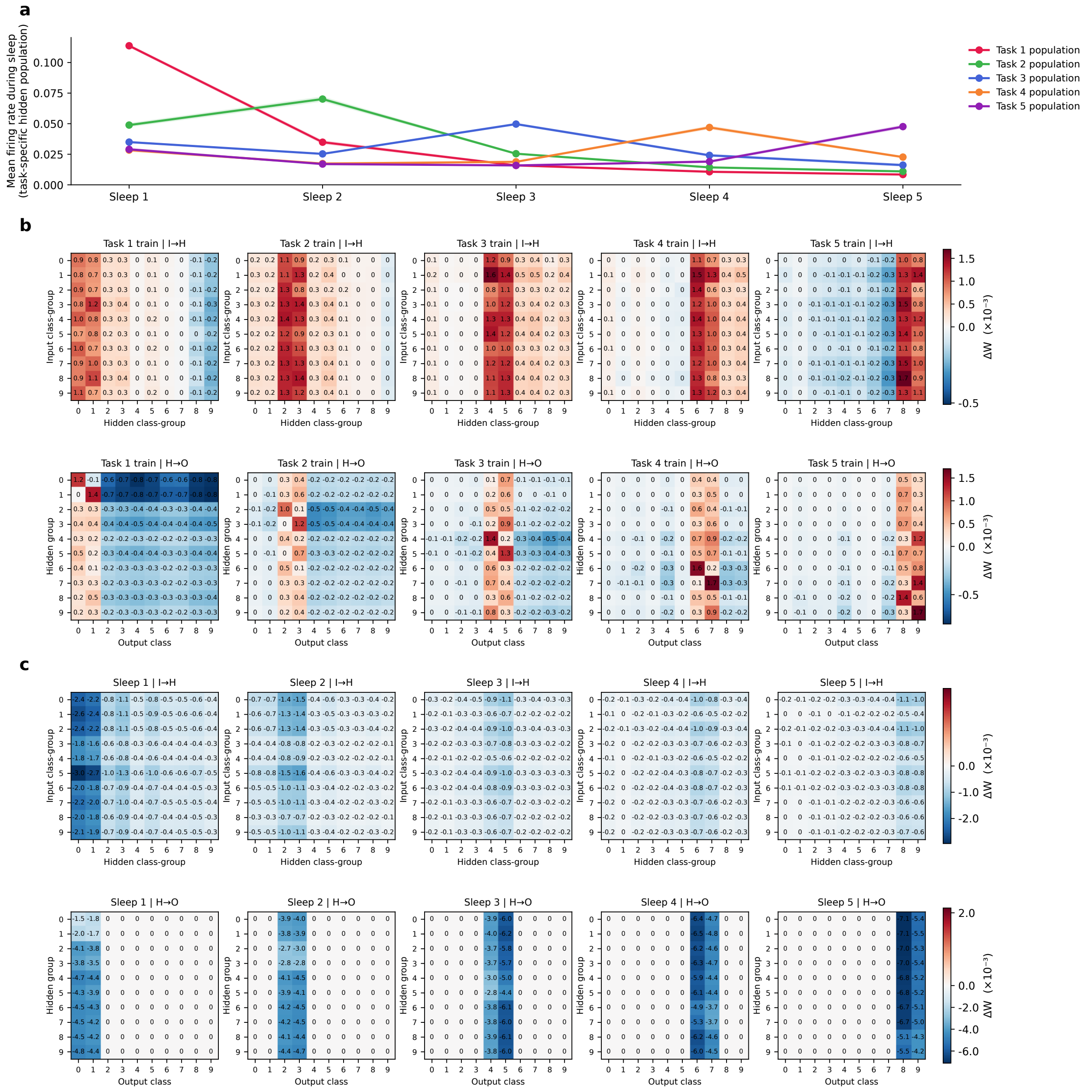


Figure S22: **MRNN-EP with sleep input based on all learned tasks: reactivation and weight changes.** Panels as in Supplementary Fig. S8; (a) mean ± SD over six trials, (b,c) trial 0.

### S7 Network architectures and MRNN-EP hyperparameters

Supplementary Figure S23 shows the architectures of MRNN-EP and RNN-BPTT, and Supplementary Table S2 lists the MRNN-EP hyperparameters for each dataset (main text, Methods).

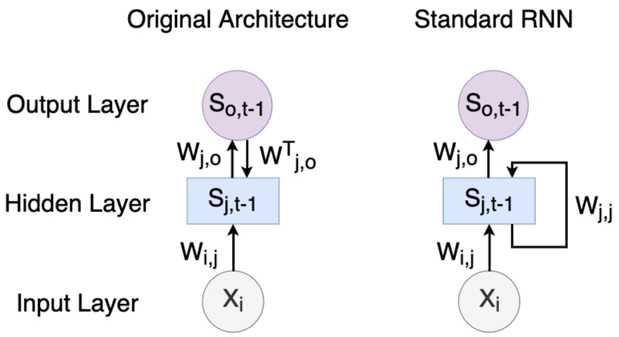


Figure S23: Architectures of MRNN-EP and RNN-BPTT. Left: MRNN-EP has feedback connections from the output layer to the hidden layer, which use the transposed hidden-to-output weights ($W^T$). Right: the standard RNN trained with BPTT (RNN-BPTT) has recurrent connections only within the hidden layer and no feedback from the output layer.

Table S2: The hyper-parameters for the MRNN-EP are summarized in this table. Here, $\alpha 1$ refers to the learning rate for updating the weights between the input and hidden layers, and $\alpha 2$ is the learning rate for updating the weights between the hidden and output layers. The 'Free Phase' and 'Clamped Phase' columns specify the number of time steps used during the free and weakly clamped phases, respectively. $\gamma$ is the feedback parameter, $\beta$ is a nudging parameter for the weakly clamped phase, and $dt$ is the Euler methods' time-step.

| Parameters | MNIST | CIFAR10 |
|---|---|---|
| $\alpha 1$ | 0.03 | 0.08 |
| $\alpha 2$ | 0.001 | 0.001 |
| $\beta$ | 1.0 | 1.0 |
| $dt$ | 0.2 | 0.2 |
| $\gamma$ | 1.0 | 1.0 |
| Free phase | 100 | 125 |
| Clamped phase | 10 | 15 |
| Hidden size | 1024 | 1024 |
| Batch size | 256 | 256 |
| Epoch per task | 3 | 5 |
